# Chromosomal Topological Domain Formation Modulates Transcription and the Coupling of Neighboring Genes in *Escherichia coli*

**DOI:** 10.1101/2025.11.23.690031

**Authors:** Nicolás N. Yehya, Christopher H. Bohrer, Qiwei Yu, David Hathcock, Boaz Goldberg, Sophia Leng, Frances E. Harris, Hunter Hendrix, Maxwell White, Fenfei Leng, Sam Meyer, Yuhai Tu, Jie Xiao

## Abstract

Chromosomal topology and transcription are tightly coupled, yet the quantitative impact of topological constraints on transcription, supercoiling, and the potential coupling between neighboring genes *in vivo* remains unclear. In this work, we constructed synthetic chromosomal domains in *Escherichia coli* that contained two genes inside a topology-controllable domain and a third gene outside. Using three-color single-molecule fluorescence *in situ* hybridization (smFISH), we measured transcription output from the three genes in individual cells under conditions in which gene orientation, domain formation state, and global chromosomal supercoiling density were varied. We found that topological domain formation repressed transcription, diminished gene orientation-dependent differences in transcription, and modulated the supercoiling sensitivity of genes located both within and near the domain. Relaxing global negative supercoiling through gyrase inhibition broadly repressed transcription; increasing global negative supercoiling level through topoisomerase I inhibition repressed highly expressed genes, while activating lowly expressed ones. Besides single-gene effects, we also observed an intrinsic coupling between neighboring genes with a non-monotonic dependence on the underlying supercoiling state, which shifted with domain topology and gene syntax. Our results establish chromosome topology as a major regulator of both transcription levels and the coupling between adjacent genes.

## Introduction

Classic transcription regulation holds that the binding of transcription factors (TFs) to gene regulatory sequences modulates the transcriptional activity of RNA polymerase (RNAP).^1,2^ While the protein-based transcription regulation mechanism remains central, recent studies have suggested that DNA supercoiling—the over- or under-winding of the double helix—coupled with the formation of chromosomal topologically isolated domains, could strongly influence transcription.^3–7^ The DNA mechanics-based influence thus adds another layer to transcription regulation in a complex cellular environment.

Previous studies have established that transcription and DNA supercoiling are tightly coupled.^4,6,8,9^ The twin-domain model describes that transcription generates positive supercoiling downstream and negative supercoils upstream of RNAP due to the rotational drag of the elongating transcription complex.^10,11^ These supercoils can spread over more than 2 kb along the DNA^12^ before they are enzymatically removed by topoisomerases.^13–15^ In *E. coli*, gyrase removes positive supercoils by catalyzing double-stranded (ds) DNA breakage and passage and introducing negative supercoils^13,16^; Topo I relaxes negative supercoils by catalyzing single-stranded (ss) DNA breakage and passage.^17,18^ The actions of the two enzymes maintain the average, global negative supercoiling density of the chromosome at ∼ -0.06 in exponentially growing cells.^19,20^

When accumulated, however, positive supercoils downstream of RNAP impede both transcription initiation and elongation, eventually stalling RNAP unless topological stress is relieved.^21^ Conversely, negative supercoils upstream of the transcription machinery can facilitate promoter opening for RNAP binding but also prevent RNAP from escaping the promoter or continuing elongation when the level is too high.^22–25^ As such, the interplay between transcription and DNA supercoils can lead to complex transcriptional behaviors, generating new dynamics and phenomena. For example, the successive dissipation and accumulation of supercoils have been proposed to cause transcriptional bursting — short transcription-on periods of rapid, successive production of multiple mRNA molecules when the torsional stress is released, interspersed by transcription-off periods when the torsional stress accumulates.^21,22^ Burstiness in gene expression generates phenotypic heterogeneity in mRNA and protein copy numbers even among genetically identical cells.^26^

Most interestingly, supercoils generated by one transcribing RNAP molecule can diffuse and influence neighboring RNAP molecules on the same or different genes, effectively coupling transcription units through shared DNA torsion.^12^ Depending on the transcription orientation (codirectional, divergent, or convergent), supercoiling can produce either cooperative or antagonistic effects on adjacent RNAP molecules and promoters.^12,27–29^ Modeling studies have shown that the annihilation of supercoiling between elongating complexes can produce emergent cooperative behavior of co-transcribing RNAP molecules, linking DNA torsional mechanics to collective transcriptional dynamics.^23,30–32^ These phenomena highlight that chromosomal supercoiling is not a mere byproduct of transcription but can act as a regulatory factor: the directionality of genes and their chromosomal context, collectively termed gene syntax^27,28^, can have substantial effects on transcription.

The impact of DNA supercoiling is especially relevant in the context of bacterial chromosomal organization, including large chromosomal interaction domains on the order of 100 kb^33^ and small topological domains on the order of 10-20 kb^7^. Because supercoils cannot diffuse across topological domain boundaries, the torsional stress they generate is shared by all genes within a domain. As such, chromosomal topological domain formation could isolate genes from the global chromosomal environment and alter their coupling and/or responses to perturbations via DNA mechanics, independent of protein-based TF regulation.^4,21,34,35^ Similar mechanisms could also be at play in eukaryotic chromosomes, where topological domains on the order of ∼ 200 kb to 2 Mb isolate and constrain supercoils, thereby impacting gene regulation within the domain differentially from genes outside the domain.^36–38^ This DNA mechanics-based transcription regulation is possibly a fundamental, universal mechanism for gene regulation, predating the more intricate, protein-based gene regulation that evolved over time, as supported by a recent comparative study.^39^

Despite growing interest in supercoil-mediated transcription regulation and several pioneering theoretical^23,27,28,30,32,40–44^ and experimental^12,21,28,35,45–48^ studies, direct, quantitative, and systematic measurements of how topological domain formation impacts the transcription of multiple genes in the chromosomal and cellular context are not available. Here, we address this gap by engineering a set of synthetic, topology-controllable chromosomal domains in *E. coli*, each containing three genes arranged in different orientations. Using single-molecule fluorescence *in-situ* hybridization (smFISH), we measured the transcriptional activity of each gene while controlling domain topology and modulating supercoiling with topoisomerase inhibitors. This synthetic-biology approach provides a precise, controllable *in vivo* system for probing the roles of topological domain formation in gene regulation.

Our approach revealed that topological domain formation profoundly affects gene expression. It represses transcription, modulates genes’ responses to supercoils, and, most interestingly, modifies the coupling between neighboring genes in a syntax-dependent manner and to different extents. Positive supercoils are essentially repressive for transcription, while negative supercoils can be activating for lowly expressed genes and repressive for highly expressed genes. Furthermore, we developed a minimal stochastic model to capture the quantitative coupling between different gene pairs and demonstrated that supercoiling can couple neighboring genes in a non-monotonic manner. These results reveal that chromosomal organization can directly shape transcription, independent of protein regulators. Our results provide a new framework for examining how chromosome architecture influences the emergence of transcriptional behaviors through supercoiling in a complex cellular environment. More broadly, these results highlight the impact of mechanical coupling between genes on expression, offering guidance for designing robust gene expression systems within their native chromosomal contexts.

## Results

### Design and construction of a controllable chromosomal topological domain in *E. coli*

To control topological domain formation at will in an otherwise identical chromosomal background and to minimize potential complications caused by native interactions, we constructed a synthetic domain of ∼ 6 kb devoid of any known TF-binding sequences (**Fig. 1A**, top left). The synthetic domain contained tandem repeats of the left and right operator sites of the λ repressor CI (3x*O_L_123* and 2x*O_R_123*) flanking two genes (*G1*, ∼ 3.4 kb, and *G2*, ∼ 1.7 kb) of identical synthetic constitutive promoters (*lacUV5*^49^) and arranged codirectionally (abbreviated as *codir* hereafter for simplicity). Each gene was terminated by strong ribosomal RNA (*rrn*) transcription terminators (*rrnB* T1 + *rrnD* for *G1* and *rrnD* for *G2*). A third gene (*G3*, ∼ 4 *kb*) with a synthetic constitutive *promoter (EM7*^50^) was placed outside *O_R_123* and terminated by the endogenous chromosomal terminator once integrated. The construct’s dimensions are listed in **Table S1**.

**Figure 1.**
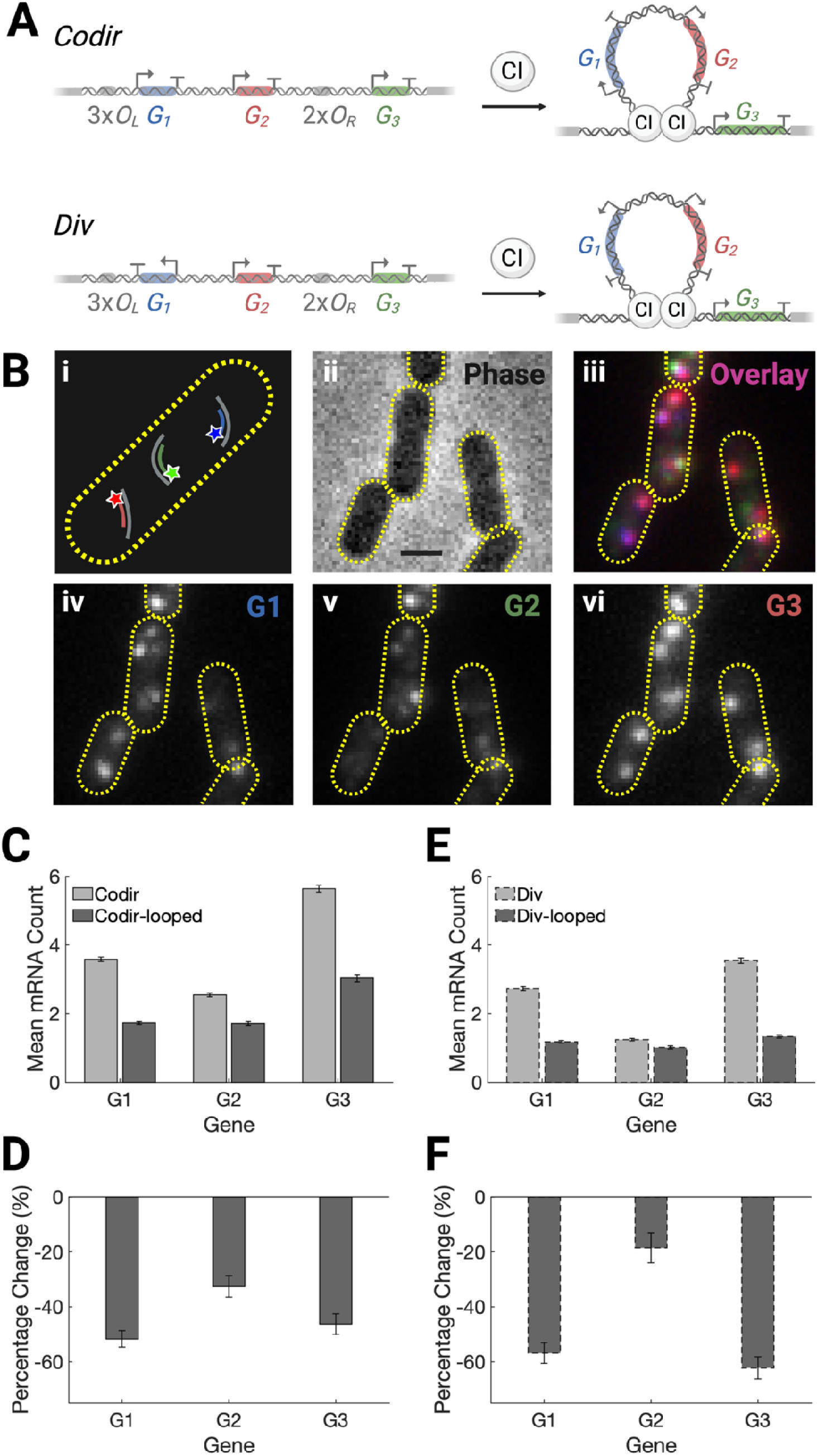
Topological domain formation represses transcription in the codirectional and divergent constructs. (A) Schematics of the chromosomal constructs. Each construct contains three genes (*G1, G2,* and *G3*) arranged either codirectionally (*codir*, top) or divergently (*div*, bottom). The binding and octamerization of the CI repressor on the flanking operator sites (3×*O_L_123* and 2×*O_R_123*) induce intramolecular looping, forming a topological domain encompassing *G1* and *G2* while leaving *G3* outside. (B) Schematics of single-molecule fluorescence *in situ* hybridization (smFISH) detection of dye-labeled probes for *G1* (blue), *G2* (green), and *G3* (red) transcripts, respectively, in a single *E. coli* cell with a yellow dashed outline (**i**), representative images of a view field of *E. coli* cells in phase-contrast (**ii**), fluorescence overlay (**iii**), and individual smFISH fluorescence channels for *G1* (**iv**), *G2* (**v**), and *G3* (**vi**), respectively. Scale bar: 1 μm. (C) Mean RNA copy number per cell for *G1, G2*, and *G3* in the codirectional construct (solid outline) under unlooped (light bars) and looped (dark bars) conditions. (D) Corresponding percentage changes of each gene in the codirectional construct upon looping. (E) Mean RNA copy number per cell for *G1, G2*, and *G3* in the divergent construct (dashed outline) under unlooped (light bars) and looped (dark bars) conditions. (F) Corresponding percentage changes of each gene in the divergent construct upon looping. Error bars represent the standard error of the mean (**Table S4**).

We integrated the domain into the *E. coli* chromosome at the *lacZ* operon location using a landing pad approach.^51^ We chose the *lac* operon location because it is in the *E. coli* right unstructured chromosomal macrodomain, which has minimal intra-macrodomain chromosomal interactions when compared to other structured macrodomains.^52^ We then grew cells in minimal medium (M9) at 30 °C and constitutively expressed the λ repressor CI from a low-copy plasmid in the cell (NY50, **Table S2**). The binding and octamerization of the λ repressor CI at the operator sites loop the intervening DNA, forming a topologically constrained domain of 6,082 bp that encompasses *G1* and *G2* but not *G3* (**Fig. 1A**, top right). As a control for the unlooped condition, we expressed an otherwise identical plasmid without the λ repressor *cI* gene in the same strain background, which does not form the synthetic topological domain (NY51, **Table S2**).

We previously used this method to investigate the effect of chromosomal DNA looping on CI-mediated transcription regulation.^53^ We verified that the expression of CI indeed led to the looping of the intervening DNA sequences in live *E. coli* cells using single-molecule imaging.^53^ Furthermore, using purified CI and the corresponding DNA construct in an *in vitro* plasmid DNA nicking topology assay^54,55^, we verified that loop formation via binding and oligomerization of CI retained supercoils in the topologically constrained segment of the plasmid (**Fig. S1**). Therefore, the synthetic platform with or without CI expression enables us to control the formation of the topological domain and subsequently examine its impact on the transcription of the three genes.

### Three-color smFISH measurements reveal differential basal transcription levels

To measure the transcription activity of all three genes simultaneously in the same cells, we employed three-color single-molecule fluorescence *in situ* hybridization (smFISH)^56^ and quantified the corresponding RNA copy numbers for *G1*, *G2*, and *G3* in exponentially growing cells based on the integrated fluorescence intensity of individual smFISH spots in cells (**Fig. 1B**, STAR Methods, **Table S3**). Under the unlooped condition (without CI expression, strain NY51), the *codir* construct expressed on average 3.6 ± 0.1, 2.6 ± 0.1, and 5.6 ± 0.1 transcripts/cell for *G1*, *G2*, and *G3* respectively (μ ± *sem*, mean ± standard error of the mean, *N* = 2,791 cells from 8 independent replicates; statistical significance assessed by two-sample Kolmogorov-Smirnov test here and hereafter unless otherwise noted, **Fig. 1C**, light gray bars, **Tables S4**, **S5**). These expression levels represent the basal expression of the three genes in the native *E. coli* chromosomal environment. Note that although *G1* and *G2* have identical synthetic promoters, their expression levels differed significantly, suggesting a potential effect due to the gene syntax.

### Topological domain formation represses transcription and diminishes gene syntax effect in a codirectional construct

We next investigated the effect of chromosomal topological domain formation by expressing CI in the same strain background and compared transcription of the three genes (looped, strain NY50) with that of the unlooped strain (NY51) under identical growth conditions. Interestingly, domain formation repressed transcription and diminished the gene syntax effect as it reduced the transcription of *G1* and *G2* to similar levels at ∼ 1.7 ± 0.1 RNA/cell, respectively (μ ± *sem*, *N* = 2523 cells from 7 independent replicates, **Fig. 1C**, **D**, dark gray bars, **Table S4**). The reduced transcription is unlikely to have resulted from the direct repression by CI-binding because the *G1* promoter is 229 bp away from the nearest *O_L_* operator, and *G2*’s promoter is in the middle of the domain, ∼ 2-4 kb away from CI’s binding sites (**Table S1**). Furthermore, we have previously shown that CI binds to its operator sites specifically and does not spread on the chromosome at our low, constitutive expression level.^53^ Therefore, these observations suggest that the formation of the topological domain represses transcription, independent of any known protein factors.

Notably, *G3*, which lies outside the loop, also exhibited a substantial reduction in transcription (3.0 ± 0.1, μ ± sem, *N* = 2523 cells from 7 independent replicates, **Fig. 1C, D**, **Table S4**). As the *G3* promoter is even further away from the nearest *O_R_*123 operator (595 bp), this observation suggests that the creation of a topological boundary can affect nearby genes over long distances, consistent with what would be expected from the DNA-mediated supercoiling effect.^21,35^

### Topological domain formation represses transcription and diminishes gene syntax effect in a divergent construct

To investigate whether the observed effects are dependent on gene orientation in the domain, we constructed a divergent (*div*) domain, in which *G1* is reversed from its orientation in the *codir* construct (**Fig. 1A**, bottom left, **Table S1**). Under the unlooped condition (Strain NY92, **Table S2**), the three genes were, in general, expressed differentially and at lower levels than they were in the *codir* construct (2.7 ± 0.1, 1.3 ± 0.1, and 3.5 ± 0.1 transcripts/cell, for *G1*, *G2*, and *G3* respectively, μ ± sem, *N* = 2,791 cells from 7 independent replicates, **Fig. 1E**, light gray bars with dashed outlines, **Table S4**). Given the identical promoter and gene sequences of the three genes between the *div* and *codir* constructs, this result suggests that relative gene orientation to each other impacts their expression levels in the native chromosomal environment.

Next, we measured transcriptional outputs of the *div* construct under the looped condition (**Fig. 1A**, bottom right; Strain NY91; **Table S2**). We found that looping also reduced *G1*, *G2*, and *G3* expression significantly (1.2 ± 0.1, 1.0 ± 0.1, and 1.3 ± 0.1 transcripts/cell, respectively, μ ± sem, *N* = 2,051 cells from 7 independent replicates, **Fig. 1E, F**, dark bars with dashed outlines, **Table S4**) in a similar fashion compared to that in the *codir* construct. In particular, *G1* and *G2* had a large difference in their expression levels under the unlooped condition but decreased to comparable levels upon domain formation (**Fig. 1E**, compare the light- and dark-gray bars of *G1* or *G2*). These observations suggest that topological confinement exerts a general repressive effect on transcription and diminishes the gene-syntax effect, regardless of gene orientation within the domain.

### Gyrase inhibition represses transcription

We reason that the above effects are likely mediated by accumulated supercoils, which can propagate along the DNA and be constrained by topological barriers. To probe this possibility, we assessed the impact of perturbing the global chromosomal negative supercoiling state by treating cells with a gyrase inhibitor, novobiocin (**Fig. 2A**). Novobiocin inhibits gyrase activity by abolishing ATP binding to the ATPase domain in the GyrB subunit^57,58^ and lowers the global negative supercoiling density (Llσ>0).^19,20^

**Figure 2.**
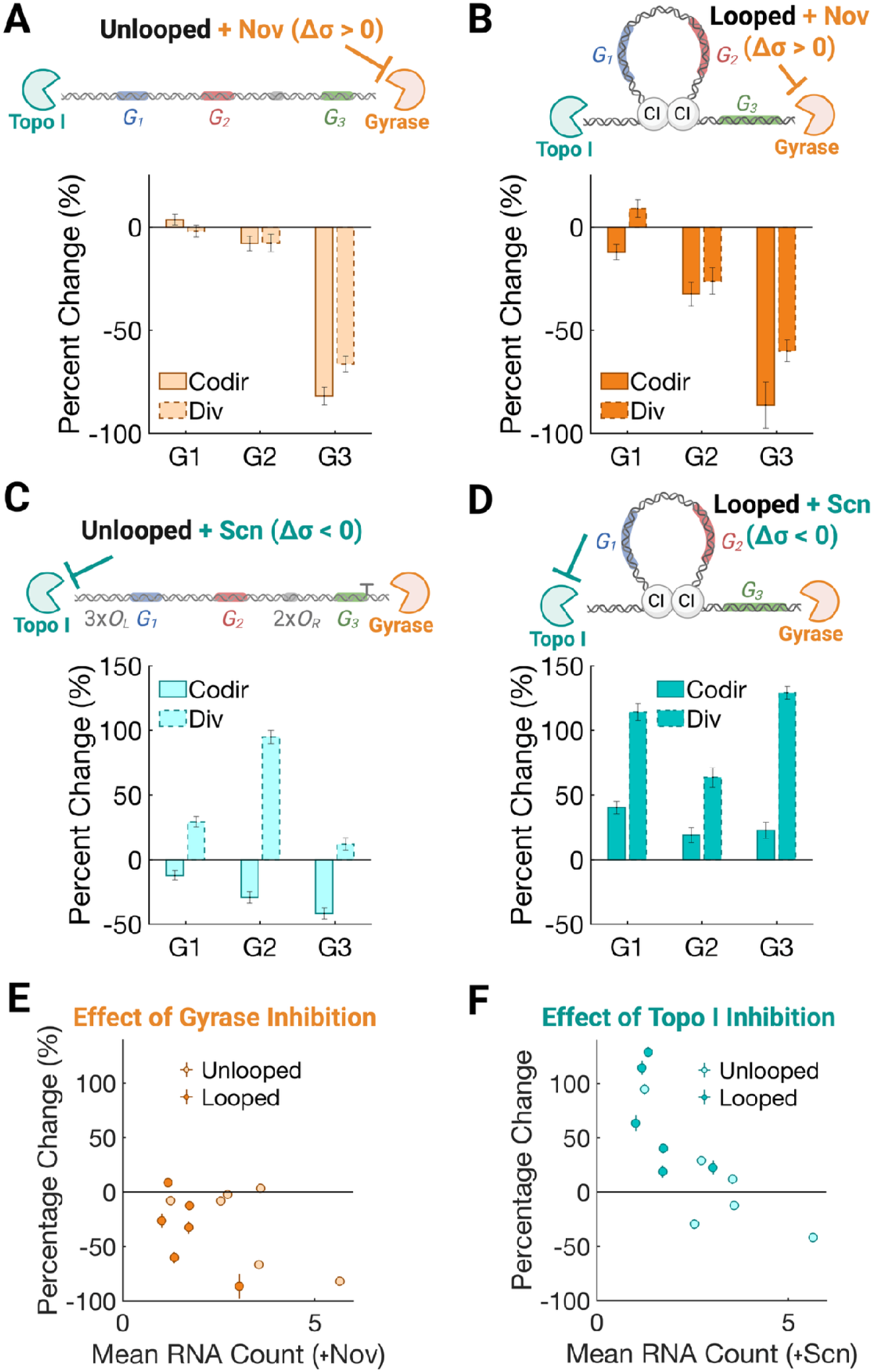
Genes respond to topoisomerase inhibition differentially in a context-dependent manner. (A) Percentage change in the RNA copy numbers for genes *G1*, *G2*, and *G3* for unlooped constructs due to gyrase inhibition by novobiocin (+Nov, top schematics). Bars with solid/dashed outlines represent codirectional/divergent constructs. (B) Same as (A) for looped codirectional (solid) and divergent (dashed) constructs. (C) Percentage change in the RNA copy numbers for genes *G1*, *G2*, and *G3* for unlooped constructs due to Topo I inhibition by seconeolitsine (+Scn, top schematics). Bars with solid/dashed outlines represent codirectional/divergent constructs. (D) same as (C) for looped codirectional (solid) and divergent (dashed) constructs. (E) Scatter plots of the baseline expression levels (x-axis, untreated) and percent change (y-axis) after novobiocin treatment. Light and dark markers represent unlooped and looped conditions, respectively. (F) Scatter plots of the baseline expression levels (x-axis, untreated) and percent change (y-axis) after seconeolitsine treatment. Light and dark markers represent unlooped and looped conditions, respectively. For all the bar and scatter plots, error bars represent the standard error of the mean.

We treated cells with 300 μg ml^-1^ (489.7 μM) novobiocin for 15 minutes before fixation under the unlooped condition. We previously used this treatment concentration and duration to avoid dsDNA breaks.^59^ Interestingly, for both the *codir* and *div* constructs (strains NY51 and NY92, respectively, **Table S2**), *G1* and *G2* were only slightly affected (∼ < 10% reduction compared to the untreated conditions, **Fig. 2A**, **Table S4**). This observation suggests that, under this unlooped condition, gyrase inhibition and the relative orientations of *G1* and *G2* have minimal effects on their responses. In contrast, *G3* expression was significantly reduced by 82 ± 4%, (*N* = 2,673 cells from 8 independent replicates), and 66 ± 4%, (*N* = 3,055 cells from 6 independent replicates) for *codir* and *div* constructs, respectively (**Fig. 2A**, **Table S4**). As *G3* has the highest expression level under the untreated condition, is the longest gene in the construct, and resides downstream of *G2*, it may experience the highest level of accumulated positive supercoils among the three genes, hence requiring a higher level of Gyrase activity to maintain its transcription. This possibility is also consistent with previous observations that the accumulation of positive supercoils often represses transcription of strong genes.^60^

### Topological domain formation sensitizes genes to gyrase inhibition

We next examined how topological domain formation modifies gene responses to gyrase inhibition. Surprisingly, under the looped condition (strains NY50 and NY91, respectively, **Table S2**), we observed enhanced responses of *G1* and *G2* but essentially unchanged responses of *G3* (**Fig. 2B**). For example, *G2* transcription was reduced by ∼ 30% for both the *codir* and *div* constructs (**Fig. 2B**, *N* = 1,999 cells from 6 independent replicates for *codir* and *N* = 2,301 cells from 6 independent replicates for *div*, **Table S4**), up from a repression level of < 10% under the unlooped condition (compare with **Fig. 2A** *G2* bars of the same styles). In contrast, *G3* was repressed at comparable high levels, regardless of whether the condition was looped or unlooped (**Fig. 2A, B**, *G3* bars). Most interestingly, under the looped condition, novobiocin produced slight, but apparently opposite effects on *G1* transcription in the two constructs where *G1* has opposite orientations: it reduced *G1* transcription by ∼ 12% in the *codir* construct, but increased *G1* transcription by ∼ 10% in the *div* construct (**Fig. 2A, B**, *G1* bars). Note that under the unlooped condition, novobiocin only had a slight effect on *G1* transcription in both constructs (**Fig. 2A**). These observations suggest that topological domain formation enhances the sensitivity of genes within the domain to gyrase inhibition, likely due to the accumulation of local positive supercoils, and that this effect depends on gene orientation.

**Figure 3.**
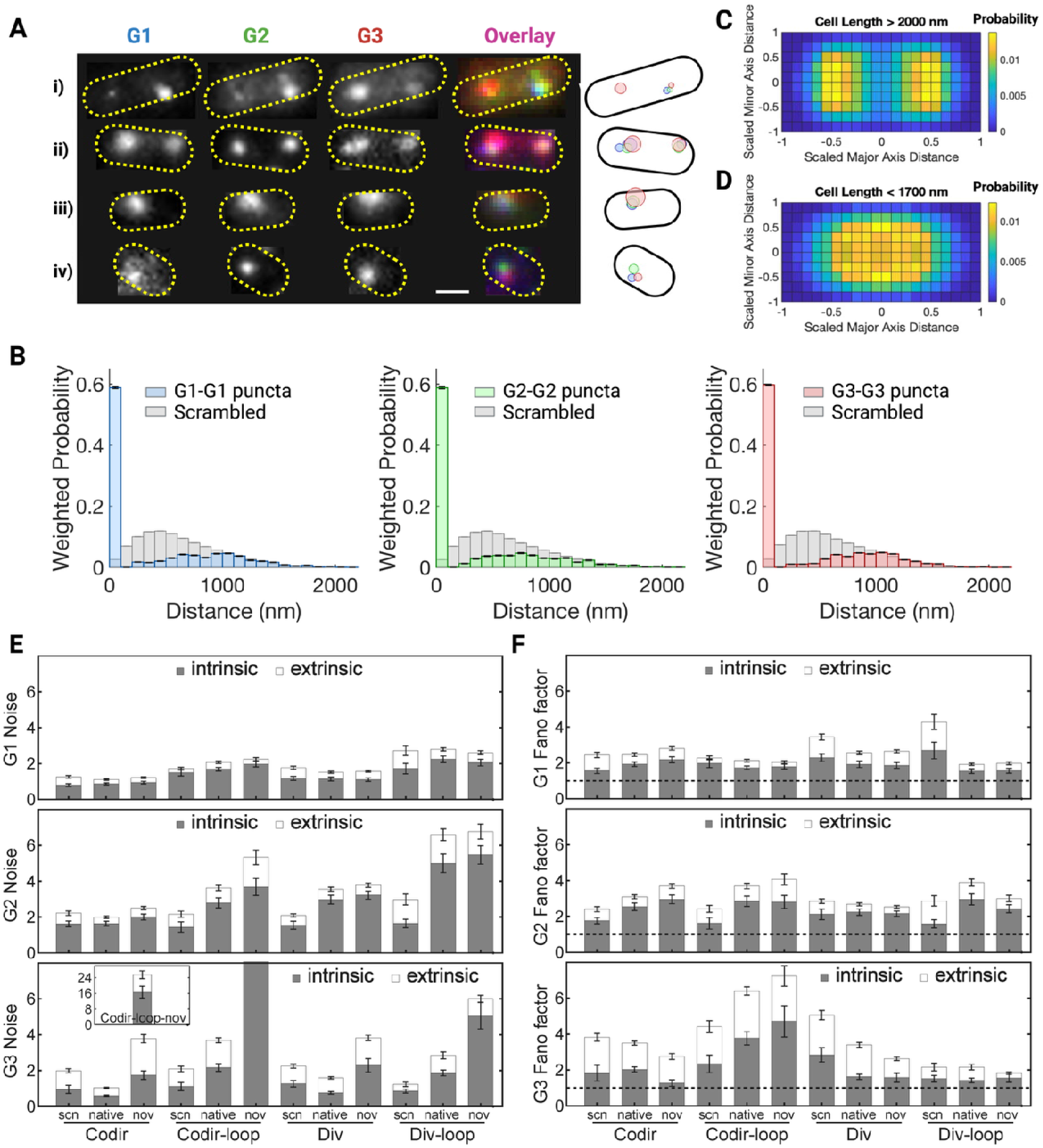
RNA puncta likely represent nascent transcription sites on the chromosome. **(A)** Representative three-color smFISH images (left) of four *E. coli* cells showing *G1* (blue), *G2* (green), and *G3* (red) RNA puncta with overlays. Cells are outlined in yellow dashed lines. The corresponding spatial maps of all puncta from these cells are shown in the right, where each punctum was colored by RNA type (blue: *G1*, green: *G2*, red: *G3*) and scaled in size according to the RNA copy number in that punctum. Scale bar: 1 μm. **(B)** Pairwise distance distributions for *G1–G1*, *G2–G2*, and *G3–G3* puncta in the codirectional construct. Distances are weighted by the number of RNA molecules in each punctum. Grey bars represent control where the puncta location is scrambled within the same cell. **(C)** Two-dimensional probability distribution of puncta localizations for long cells (> 2.0 µm) across all conditions and all RNAs. The cell size is rescaled to a standard size of 1 μm x 3 μm. **(D)** Same as (C) for short cells (< 1.7 µm). **(E)** Decomposition of intrinsic and extrinsic noises for the three genes across all conditions. Intrinsic noise (gray bars) dominated, and looping in general increased the intrinsic noise of the corresponding condition. Error bars are estimated via bootstrap resampling. **(F)** Decomposition of intrinsic and extrinsic Fano factors for the three genes across all conditions. Nearly all conditions had intrinsic Fano factors significantly > 1 (dashed lines). Error bars are estimated via bootstrap resampling.

### Topoisomerase I inhibition alters transcription in a syntax-dependent manner under the unlooped condition

Having examined the impact of inhibiting gyrase—which led to decreased negative supercoiling densities (Δσ>0)—we next investigated how increased negative supercoiling densities (Δσ<0) affect transcription. Specifically, we shifted the global chromosomal supercoiling state in the opposite direction by treating cells with a Topoisomerase I inhibitor, seconeolitsine.^61^ Topoisomerase I relaxes negative supercoiling, so its inhibition leads to a higher level of accumulated negative supercoiling in the chromosome.^60^

We treated cells with 25 μM seconeolitsine for 15 minutes before fixation.^60^ In *codir* cells, seconeolitsine reduced *G1*, *G2*, and *G3* expression by 12 ± 4%, 29 ± 5%, and 42 ± 4%, respectively (*N*= 1,081 cells from 3 independent replicates, **Fig. 2C**, bars with solid outlines, **Table S4**), indicating that for the codirectional arrangement, bias toward both more and less negative supercoiling density repressed transcription. However, treating the *div* construct with seconeolitsine under the unlooped condition enhanced the transcription significantly for all three genes but to different extents (29 ± 4%, 92 ± 4 %, and 12 ± 5% for *G1*, *G2*, and *G3* respectively, *N* = 1,046 cells from 3 independent replicates, **Fig. 2C**, bars with dashed outlines, **Table S4**). These results suggest that the gene syntax under the unlooped condition may significantly impact individual gene expression and lead to drastically different responses to topoisomerase I inhibition.

### Topoisomerase I inhibition activates transcription under the looped condition

Next, we investigated how the three genes respond to Topo I inhibition under the looped condition. Interestingly, treating cells with seconeolitsine under the looped condition enhanced transcription for both the *codir* and *div* constructs; the expression of *G1*, *G2*, and *G3* were enhanced by 40 ± 5%, 19 ± 6%, and 23 ± 6% (*N*= 618 cells from 3 independent replicates) respectively for the *codir* constructs, and 114 ± 7%, 66 ± 8%, and 129 ± 5%, respectively (*N* = 605 cells from 3 independent replicates) for the *div* construct (**Fig. 2D**, **Table S4**). As Topo I inhibition leads to the accumulation of more negative supercoils, these observations suggest that, under the looped condition, accumulated negative supercoils activate transcription, in contrast to the effect of accumulated positive supercoils when Gyrase was inhibited (compare **Fig. 2B** and **D**). Note that *G3*, which is outside of the domain boundary, also responded differentially to seconeolitsine under the unlooped and looped conditions, indicating an impact of domain formation for genes near the boundary.

### Supercoiling sensitivity depends on the basal transcription level

The differential, nonuniform responses of the three genes in the *codir* and *div* constructs under the looped and unlooped conditions suggest that other factors, in addition to gene orientation or topological domain formation, may impact the sensitivity of gene transcription to supercoils. As transcription also generates supercoils, we reasoned that genes with different transcription levels may be differentially sensitive to topoisomerase inhibition. To examine this possibility, we calculated the percent change in RNA copy number between drug-treated and untreated conditions across all constructs, genes, and looping states, and plotted the percent change relative to the mean RNA copy number in the untreated condition for each matched pair (**Fig. 2E, F**).

For novobiocin-treated conditions (**Fig. 2E**), we observed a general repression trend (percentage change < 0) regardless of the basal expression levels, with no statistically significant dependence on the expression level or the looping state (*p_Spearman_* = -0.329, *p* = 0.3). The repression suggests that accumulated positive supercoiling resulting from gyrase inhibition imposes a general transcriptional penalty, likely by preventing transcription initiation or by stalling RNAP during elongation. In contrast, seconeolitsine treatment produced a significant anti-correlated trend (*p_Spearman_* = -0.811, *p* = 0.002, **Fig. 2F**); genes with low expression levels showed higher levels of activation, while genes with higher expression levels were generally repressed after seconeolitsine treatment. This observation suggests that negative supercoils can be either activating or repressive, depending on the initial transcriptional level. Hence, negative supercoils may act as a homogenizing force, bringing genes with too low or too high expression levels back into a nominal range.

### An individual gene’s transcription level depends on its syntax

So far our experiments have shown that the transcription levels of the three genes varied in a large range from ∼ 0.4 copies per cell to ∼ 5.6 copies per cell under twelve different conditions (**Table S4**), each being a unique combination of three factors, the construct orientation (*codir* or *div*), looping state (looped or unlooped), and drug condition (untreated, +nov, or +scn). To assess the impact of each factor on the transcription levels of the three genes, we applied a three-way analysis of variance^62^ to the measured RNA copy numbers for each gene across all conditions (STAR Methods). We found that nearly all factors and their combinations significantly affected transcription levels (F statistics >> 1), but the degree of impact varied across genes (**Table S6**). Specifically, *G1* transcription was most strongly affected by the looping state (the greatest F-statistics), *G2* by construct orientation, and *G3* by drug condition. These results suggest that, for genes located within a topological domain such as *G1* and *G2*, transcription is highly sensitive to domain formation and relative gene orientation, demonstrating a gene syntax effect. In contrast, a gene outside a domain, such as *G3*, may be most sensitive to changes in the chromosomal supercoiling state, even though it still responds to neighboring domain formation.

### RNA puncta likely represent nascent transcription sites on the chromosome

In smFISH images, we often observed that many cells showed discrete RNA puncta that contained more than one RNA molecule from the same gene (**Fig. 3A**). Here, we define an RNA punctum as a diffraction-limited fluorescent spot (radius *r* of ∼ 100 nm) detected above the cell background (STAR Methods). The average number of RNA molecules per punctum varied depending on different conditions and genes, but was generally in the range of 1-10 copies (**Fig. S2**, **Table S7**). The average number of puncta per cell varied but was generally in the range of 0-4 (**Fig. S3**, **Table S7**).

We reason that these RNA puncta likely represent nascent transcription sites, because the likelihood of detecting newly produced RNA molecules still attached to or near the transcription site before they diffuse away or are degraded should be significantly higher than that of randomly diffusing RNA molecules colocalizing with each other by chance. To examine this possibility, we calculated the pairwise distance distributions of all RNA molecules from the same gene using the *codir* construct as an example (STAR Methods, **Fig. 3B**, colored bars). We then compared them with those calculated by computationally scrambling the spatial coordinates of RNA molecules in the same cells (**Fig. 3B**, gray bars). We observed distinct peaks centered at the first 100 nm bin for all the three RNAs, in contrast to broad peaks around 500 nm in the scrambled controls, which reflected the cell radius (**Fig. 3B**). In STAR Methods, we show that the mean displacement, *r*, of an RNA molecule away from its chromosomal transcription site is only related to its diffusion coefficient *D* and degradation rate *λ* by < r^2^ > = 6D/*λ*. Given the relatively fast mRNA degradation rates in *E. coli* cells (on average ∼ 1/min^63^), the distinct peaks at the diffraction-limited resolution strongly suggest that these RNA molecules diffuse slowly inside the cells (*D* <∼ 10^-5^ µm^2^.s^-1^). As such, the corresponding RNA puncta likely represent nascent transcription sites where transcribed RNAs are still attached to the chromosomal DNA. We observed similar pairwise distance distributions for RNA molecules from the same genes or different genes under all other conditions (**Fig. S4**).

Besides the prominent peaks at the first 100-nm bin, we also observed a minor, broader peak centered ∼ 1000 nm for the pairwise distance distributions (**Fig. 3B**). As this distance is on par with that between two segregated nucleoids, we reasoned that it likely arose from two nascent transcription sites, one on each replicated chromosome. To examine this possibility, we used the *codir* construct as an example and analyzed the relationship between cell length (a proxy for cell cycle time) and the number of RNA puncta per cell, as longer cells are more likely to have replicated and segregated their chromosomes. We observed that there was a significant correlation for all the three genes: cells with only one RNA punctum centered at shorter lengths while cells with two or more RNA puncta shifted to longer lengths (**Fig. S5A-F**, *G1* *p_Spearman_* = 0.38, *p* <<0.001, *G2* *p_Spearman_* = 0.28, *p* <<0.001, *G3* *p_Spearman_* = 0.39, *p* <<0.001). The copy number of RNA molecules per punctum did not exhibit such a cell-length dependence (**Fig. S5G-I,** *G1* *p_Spearman_* = 0.03, *p* = 0.18, *G2* *p_Spearman_* = -0.01, *p* = 0.53, *G3* *p_Spearman_* = 0.01, *p* = 0.73). Based on these observations, we aggregated all puncta across all conditions in long cells (> 2 μm) and plotted the average, normalized two-dimensional (2D) histograms for a cell of standard size (1 μm x 3 μm). We observed a typical two-lobed shape (**Fig. 3C**), reminiscent of replicated and segregated nucleoids as we and others previously observed.^59,64^ In contrast, short cells (< 1.7 μm) exhibited a single-lobed distribution (**Fig. 3D**). Normalized 2D histograms of RNA puncta in all other individual conditions showed similar patterns (**Fig. S6**). Taken together, these observations strongly suggest that RNA puncta are likely nascent transcription sites on the chromosome, where RNA molecules are still attached to or in proximity to their chromosomal gene locus.

### Intrinsic variations of RNA copy number per cell suggest bursty transcription

Previous studies have established that for a random, birth-and-death transcription process, the distribution of RNA copy number per cell follows a Poissonian distribution, with the Fano factor (*f* = σ2/µ) equaling one.^65,66^ A super Poissonian distribution with *f* > 1 often indicates a bursty transcription process, in which a gene is only switched on for a short period of time to produce multiple transcripts before it is switched off.^67,68^ Therefore, we compiled distributions of RNA copy number per cell in different conditions to examine the corresponding transcription mode. To avoid the complication of transcription from two replicated chromosomes in the same cell, we used data from short cells (< 1.7 µm) where most cells only contained one unreplicated chromosome. We observed that most conditions showed non-Poissonian distributions and were best fit with a negative binomial distribution, indicative of transcriptional bursting ^69,70^ (**Fig. S7**). However, previous studies, including ours, have shown that a non-Poissonian distribution can also result from extrinsic noise, *i.e*., cell-to-cell variation.^71–73^ Specifically, fluctuations in RNAP levels and the cellular environment across different cells, which we collectively termed the extrinsic variable E, could lead to heterogeneous transcription rates and, consequently, significant variations in RNA copy number distributions, in addition to intrinsic variations caused by the inherent stochasticity of the corresponding transcription mode.

To separate the influence of extrinsic noise and identify the intrinsic transcription mode, we made use of the observation that in long cells (> 2.0 μm), RNA puncta were well separated between the two halves and likely represented nascent transcription sites on two replicated chromosomes (**Fig. 3C**). As the same gene on the two replicated chromosomes shared the same extrinsic RNAP level and cellular environment, the RNA copy number variation between the two halves of the cell reflects the intrinsic stochastic nature of transcription at the same extrinsic variable E, while that across different cells (different *E* values) reflects the influence of both the intrinsic and extrinsic stochasticity. Therefore, the total noise (variation across all cells, 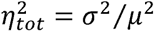) for each gene can be decomposed into intrinsic and extrinsic contributions, 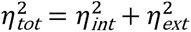, similar to what was previously performed on two different genes in the same chromosome (STAR Methods).^72,74^

Using this approach, we computed the extrinsic and intrinsic noise of each gene based on its RNA numbers in the two halves of long cells (length > 2.0 μm, STAR Methods). We observed that, across all conditions, while extrinsic noise contributed significantly (**Fig. 3E**, white bars), the total noise remained dominated by intrinsic noise (**Fig. 3E**, gray bars; **Table S8A**), suggesting a difference between transcription and translation, with the latter often dominated by extrinsic noise.^71,73,75^ Topological domain formation increased intrinsic noise, in general, most likely due to the repressed transcription levels (compare the intrinsic noise levels of the looped *v.s.* unlooped conditions in **Fig. 3E**). Indeed, plotting the decomposed intrinsic and extrinsic noise components against the mean RNA expression levels showed the expected trends that the intrinsic noise decreased as expression levels increased. In contrast, extrinsic noise was largely independent of expression levels (**Fig. S8A, B**). Most importantly, the intrinsic Fano factors of all three genes were, under most conditions, still significantly larger than one and independent of expression levels, demonstrating a non-Poissonian, bursty-like transcription mode (**Fig. 3F**, gray bars; **Table S8B**; **Fig. S8C, D**).

We note that the approach of using the two halves of long cells as proxies for two chromosomal gene copies relies on the assumption that individual transcripts diffuse slowly across the two halves of the cell, so that there is minimal mixing before the transcripts are degraded. The observation of diffraction-limited RNA puncta with multiple RNA copies in many puncta, as in **Fig. 3A-C**, suggests that this assumption is reasonable. As a further control, we computationally scrambled the number of RNA molecules across the two halves of the cells and recomputed the corresponding intrinsic and extrinsic noise. We observed that the resulting intrinsic noise diminished significantly to nearly zero, whereas extrinsic noise increased (**Fig. S9A**, STAR Methods). This computational control validated the minimal mixing assumption of transcripts between the two cell halves and also suggested that the calculated intrinsic noise was the lower bound of the true intrinsic noise, as any mixing between the two halves of the cells would only reduce the apparent intrinsic noise (STAR Methods).

### Correlation analysis suggests coupled transcription between gene pairs

In smFISH imaging, we observed that RNA puncta from different genes often colocalized (**Fig. 3A, Fig. S4**). This observation not only supported the interpretation that they likely represented nascent transcription sites from the corresponding chromosomal locus but also raised an interesting question of whether the expression of different genes could be correlated in time. We reasoned that if two genes from the same chromosomal locus transcribed independently from each other, their RNA copy numbers should be uncorrelated. However, if one gene’s transcription were coupled to another gene, for example, *G1* is only on when *G2* is on (or off), their RNA copy numbers would be correlated (or anticorrelated).

To examine this possibility quantitatively, we calculated the Spearman correlation coefficients for all gene pairs under all conditions (**Table S9**). To avoid potential complications brought by two replicated chromosomes, we only used cells shorter than 1.7 μm. We observed varied but significant correlations between gene pairs in most conditions, except for a few (**Table S9**). For example, *G1* and *G2* transcription correlated significantly with each other in ten out of twelve different conditions (all except the *codir-scn* and *div-loop-scn* conditions). Similarly, *G2* and *G3* transcription also showed significant correlations or anticorrelations in all except the *div-nov* condition. As a control, when we computationally scrambled RNA copy numbers for a single gene across all cells within the same condition, its correlation coefficient with the other genes dropped to nearly zero (**Table S9**). These results strongly suggest that genes within the same local chromosomal context could be coupled in transcription, even though each gene has its own independent promoter and terminator.

### Covariance analysis demonstrates intrinsic gene-gene coupling

What could lead to the observed correlations among different gene pairs? We reason that neighboring genes could influence each other’s transcription through the supercoils that they generate and share during transcription on the same chromosomal DNA. If so, analyzing the intrinsic correlation between different gene pairs under various conditions may identify important factors modulating the DNA mechanics-based gene-gene interactions. However, as we described above for single-gene noise analysis, gene-gene correlation can also arise from shared external factors, such as RNAP levels and cellular environments. For example, if the RNA counts of two genes (*m*_1_, *m*_2_) both depend on a shared extrinsic variable *E* (**Fig. 4Ai**), any variation in *E* could modulate the apparent correlation between *m*_1_ and *m*_2_ (**Fig. 4Aii**)

**Figure 4:**
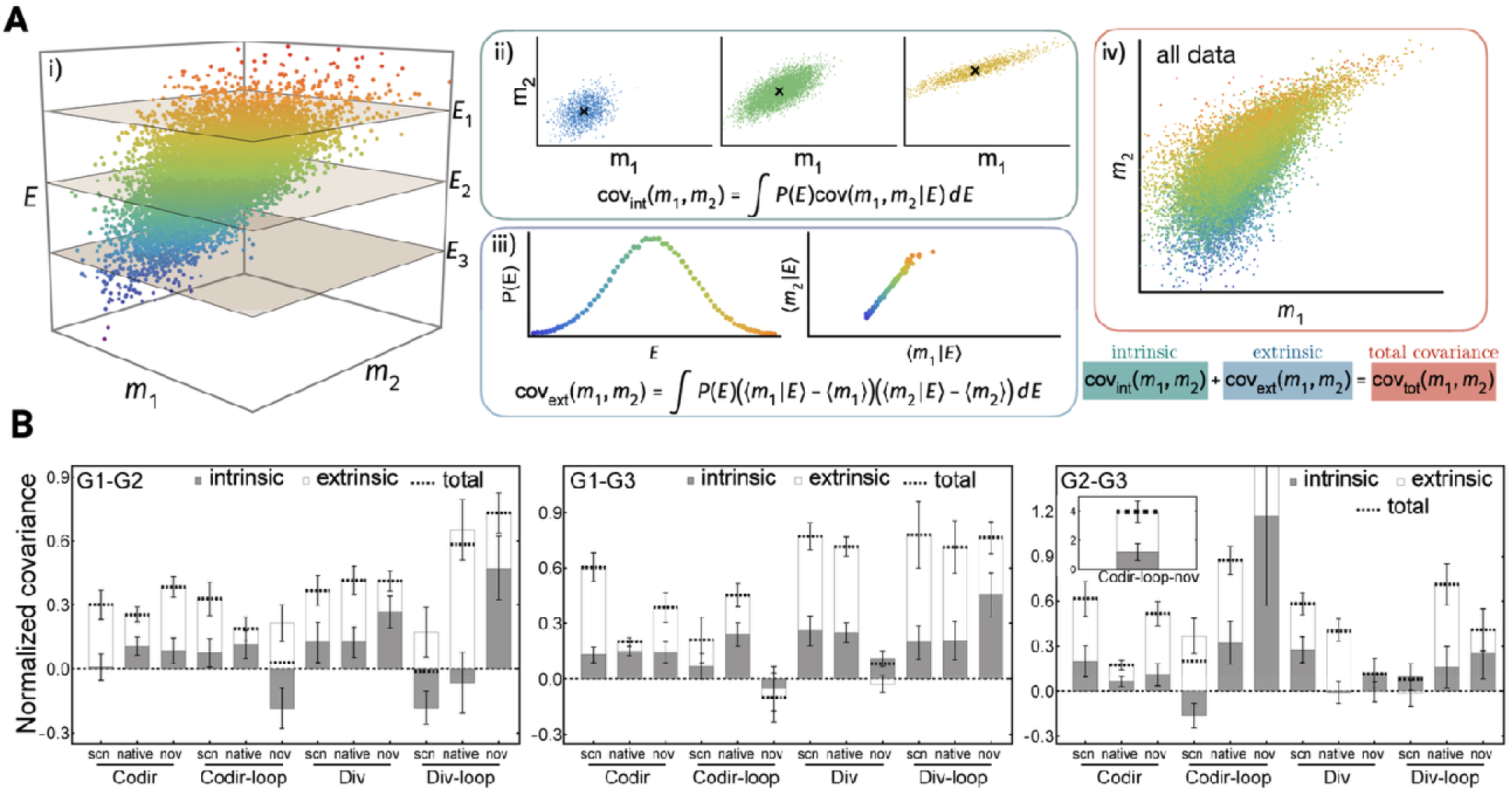
Intrinsic-extrinsic decomposition of covariances of gene pairs across all conditions reveals gene-gene coupling. (A) Illustration of the effect of extrinsic variability on the covariance between the expression of two genes. All data in (**A**) were synthetically generated to demonstrate the decomposition procedure. (**i**) The distribution of measured RNA copy numbers, and, may depend on extrinsic, or cell-to-cell variability . (**ii**) The intrinsic covariance be-tween and is defined as the covariance at fixed averaged over the extrinsic variable distribution . (**iii**) The extrinsic covariance is the covariance between condi-tional means, and, over the extrinsic distribution . (**iv**) The total co-variance, computed across all observed data, is equal to the sum of intrinsic and extrinsic components. (B) Intrinsic-extrinsic covariance decompositions of *G1-G2*, *G1-G3*, and *G2-G3* gene pairs, normalized by the product of the corresponding mean expression levels. While extrinsic variation (white bars) contributes more significantly to the apparent total covariance, the intrinsic components (gray bars) are significantly non-zero for most condi-tions.

To distinguish the intrinsic and extrinsic contributions to the observed gene-gene correlations, we extended a previous mathematical framework that separates the two components from the expression variations of a single gene.^76^ This new framework allowed us to decompose the total covariance *cov_tot_*(*m*_1_, *m*_2_) of expressed RNA molecules from two genes into intrinsic *cov*_*int*_(*m*_1_, *m*_2_) and extrinsic *cov*_*ext*_(*m*_1_, *m*_2_) components (**Fig. 4A**, STAR Methods). Here, the transcript numbers (*m*_1_, *m*_2_) depended on extrinsic variables collectively denoted by *E* (**Fig. 4Ai**). Using the same strategy of approximating the two halves of long cells (length > 2.0 μm) as two replicated chromosomes, we obtained from two halves of the same cell two samples of (*m*_1_, *m*_2_) under the same value of E, while different cells provide samples for different values of E. We defined the intrinsic covariance as the covariance between transcripts from the same gene pair in two halves of the same cell (fixed value of the extrinsic variable E), averaged over the distribution of *E* (**Fig. 4Aii).** The extrinsic covariance was defined as the covariance between the mean expression levels (*m*_1_|E) and (*m*_2_|E) due to variations in the extrinsic variable *E* (**Fig. 4Aiii**). The sum of intrinsic and extrinsic covariances is equal to the total covariance, which can be computed equivalently by lumping the data across all cell halves (all *E* values, **Fig. 4Aiv)**. As such, the extrinsic covariance in this decomposition reflects global co-regulatory mechanisms that simultaneously impact both genes, while intrinsic covariance isolates the local, mechanics-based gene-gene coupling.

Using this framework, we computed the covariances of different gene pairs across all conditions (**Fig. 4B**). We observed that while gene pair covariances were dominated by extrinsic contributions, the intrinsic components were significantly non-zero under most conditions and even negative in some conditions, demonstrating the presence of local gene-gene coupling independent of the shared cellular environment. Most interestingly, the *G1-G3* pair, which was not immediately adjacent on the chromosome, also showed significant intrinsic covariances (**Fig. 4B**, middle panel), strongly suggesting a coupling mechanism mediated by long-distance interactions, such as supercoils.

### A gene-gene coupling model describes joint RNA distributions quantitatively

The covariance analyses so far demonstrated that transcription of adjacent genes is indeed coupled through an intrinsic mechanism. However, it does not quantify the coupling strength or provide insights into the coupling mechanism. Additionally, it is unclear how different conditions impacted the intrinsic covariance. To address these questions, we constructed a gene-gene coupling model that describes the joint distribution of a gene pair’s transcription quantitatively (**Fig. 5A**). In this model, a gene i (=1,2) switches on and off stochastically with rates *k*_*on*,i_ and *k*_*off*,i_ respectively. In the *on* state, the gene produces transcripts at a rate µ_i_, which are degraded at a constant rate d_i_ independent of the gene state. The switching rates, however, are dependent on the state of the other gene. For example, when *G2* is on, the switching on and off rates of *G1* are modified from (*k*_*on*,1_, *k*_*off*,1_) to (*k*^’^_*on*,1_, *k*^’^_*off*,1_), and *vice versa*. The observed positive covariance in transcription arises because, for example, when *G2* is active, *G1* is more likely to turn on (see STAR Methods for mathematical model definition and analysis of relevant parameter regimes). Conversely, negative covariance occurs when *G2* is active, *G1* is more likely to be turned off. This model is therefore a two-gene generalization of the transcriptional bursting process as previously described^77^, but it goes beyond the transcription kinetics of individual genes to describe the joint distributions of a gene pair’s transcription, which result from gene-gene coupling.

**Figure 5:**
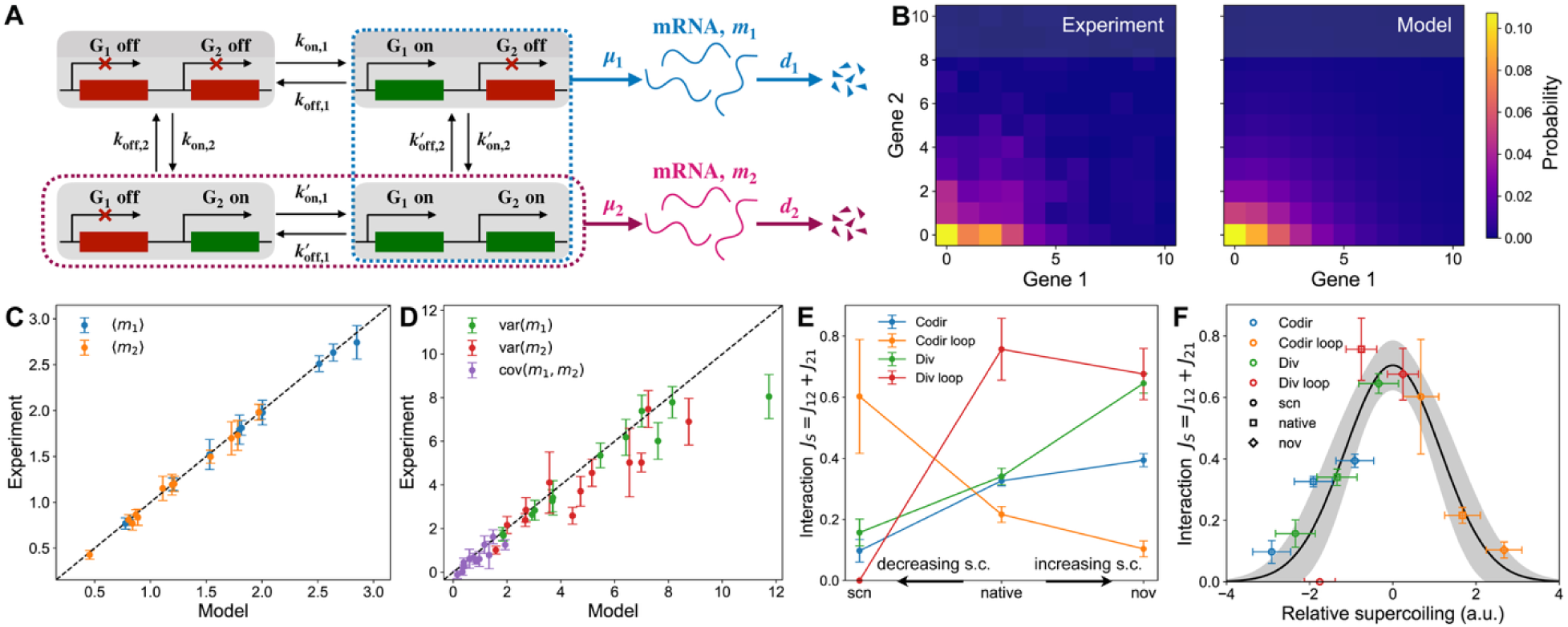
A four-state theoretical model captures coupled stochastic gene expression. **(A)** The four-state model of coupled transcription of two genes (*G1, G2*). Each gene stochastically switches between on and off states, with transition rates dependent on the state of the other gene. Transcripts are produced at a rate µ_i_ during the *on* state and degraded at a rate d_i_. **(B)** Experimentally measured joint RNA count distribution P(*m*_1_, *m*_2_) (left) and the model fit (right) of the codirectional construct. **(C)** Mean expression levels of *G1* and *G2* from the model (x value) agree with experiments (y value) across all conditions. Error bars are bootstrapped errors. **(D)** Variance and covariance of G1 and G2 from the model (x value) agree with experiments (y value) across all conditions. Error bars are bootstrapped errors. **(E)** The total coupling strength parameter *J*_S_ = *J*_12_ + *J*_21_ varies due to drug treatments by seconeolitsine (scn) and novobiocin (nov). Arrows indicate whether drug treatments increase (Δσ > 0) or decrease (Δa < 0) overall supercoiling density. Colors represent different constructs and looping conditions. Error bars are uncertainties in fitting gene expression to the four-state model. **(F)** The construct-dependent response in **(E)** can be explained by a non-monotonic dependence (here demonstrated by a bell curve) of the coupling strength *J*_S_ as a function of relative supercoiling. Colors represent constructs, and shapes indicate drug treatment conditions. Horizontal error bars and the grey shading represent uncertainties associated with placing points on the bell curve, and vertical error bars represent uncertainties in fitting gene expression to the four-state model. All results in this figure were generated by including extrinsic variations in µ_i_ with an amplitude equal to 50% of the upper bound estimated from long cells (see STAR Methods for details and justification).

To account for the extrinsic variability observed in experiments, we incorporated into the model a distribution of transcription production rates, µ_i_, commonly used to represent the effects of extrinsic fluctuations, such as RNAP concentration and cellular resources.^73,74^ The experimentally determined extrinsic variance (noise) and covariance (coupling) can then be attributed to fluctuations in the production rates. Because our estimate of extrinsic noise from long cells is an upper bound, we fixed the distribution of production rates to match 50% of the observed extrinsic noise and covariance amplitudes (see STAR methods for details). Thus, our gene-gene coupling model captured both intrinsic and extrinsic statistics of transcription.

Using a maximum-likelihood approach, we fit the experimentally measured joint RNA copy-number distributions of gene pairs under each condition to the theoretical model at half the maximal extrinsic noise level (see one example of *G1*-*G2* in **Fig. 5B**; other pairs and associated model parameters are in **Fig. S10** and **Table S10**). To avoid accounting for transcripts produced by replicated chromosomes, we fit only data from short cells (< 1.7 µm). We found that the model successfully captured the joint distribution of transcript counts P(*m*_1_, *m*_2_) across all conditions, including the mean (**Fig. 5C**), variances, and covariances (**Fig. 5D**). As the joint distribution fitting took into account the extrinsic covariance, it provides intrinsic transcriptional parameters (**Table S10**). Reducing extrinsic noise and covariances to zero or increasing them to the experimentally measured maximal level produced modest differences in kinetic parameters, but the trends across conditions were essentially the same (**Fig. S10B, Table S10**).

### Coupling strength between gene pairs depends on supercoiling non-monotonically

The gene-gene coupling model enabled us to quantify the coupling strength between a gene pair and to analyze how it was affected by different conditions. We defined an coupling strength parameter *J*_ij_ as the relative change of the burst, or switch-on, frequency of G j due to G i activation: *J*_ij_ = f_*on*,i_ (f_j_ ’/f_j_ - 1), where f_*on*,i_ = *k*_*on*,i_/(*k*_*on*,i_ + *k*_*off*,i_) is the fraction of time that gene i is on; f_j_ ’ = *k*^r^_*on*,j_/(*k*_*off*,i_ + *k*^r^_*on*,j_) and f_j_ = *k*_*on*,j_/(*k*_*off*,i_ + *k*_*on*,j_) are the probabilities that gene j switches on while gene i is on for coupled and uncoupled activation, respectively (STAR Methods). This coupling can be interpreted as frequency of coupling, quantified by f_*on*,i_, times the magnitude of the coupling. As our model does not constrain the directionality of coupling (see Discussion), we used the overall coupling strength *J*_S_ = *J*_12_ + *J*_21_between the two genes to characterize their coupling strength. We calculated the]_S_ values for all pairs across all conditions (**Table S10**) and found that the coupling strength varied widely across conditions and was independent of joint expression levels (**Fig. S11**). Instead, the coupling strength appeared to be increased by novobiocin (nov) treatment while decreased by seconeolitsine (scn) treatment (see *G1-G2* pair as an example in **Fig. 5E**, blue and green lines). However, this trend was reversed or became non-monotonic when the constructs were looped (**Fig. 5E**, red and orange lines). For the other two gene pairs, we also observed that both drug treatments and looping state modulated the coupling strength (**Fig. S12**).

Since novobiocin and seconeolitsine influence supercoiling in opposite ways, and the local supercoiling level of any gene experiences may depend on its syntax, we hypothesized that the coupling strength between two genes may be modulated by the local supercoiling density non-monotonically, in that too high or too low a supercoiling density would decrease the coupling strength. To illustrate this point, we constructed a model depicting the relationship between supercoiling and coupling strength as follows.

As we cannot measure the local supercoiling density in our experiments, we assumed that each construct (*codir* or *div*) under the looped or unlooped condition has a native supercoiling density σ = *x_c_*, which is shifted one arbitrary unit (*a.u*.) to σ = *x_c_* +1 by novobiocin and σ = *x_c_* -1 by seconeolitsine, respectively. We then considered a bell curve with a height of *J*_max_ and width of σ_c_to fit each construct’s *x_c_* on this curve based on its measured *J*_S_ value (STAR Methods). We chose a bell curve to describe the relationship because of its mathematical simplicity for non-monotonic trends (only two parameters are required), but other qualitatively similar non-monotonic curves could also capture the relationship between coupling and supercoiling density.

Using this model, we observed that *J*_S_ of *G1-G2* for all twelve conditions was well described by a single bell curve (**Fig. 5F**). Fitting *J*_S_ of other gene pairs produced similar results (**Fig. S12**). The conformity of all *J*_S_ values for each gene pair around a bell curve provides a qualitative estimate of the native supercoiling level *x_c_* underlying the joint transcriptional response of each gene pair under each condition. For example, the *G1-G2* pair in unlooped conditions (**Fig. 5F**, blue and green markers) exhibited more negatively supercoiled levels irrespective of the gene orientations (left side of the bell curve) compared to their looped conditions (**Fig. 5F**, orange and red markers), consistent with the observation that topological domain formation isolates genes from the globally negatively supercoiled chromosomal DNA and represses their expressions (**Fig. 1C-F**). The modeled *G1-G2* coupling strengths of a few conditions, including *div*-loop (red square), *div*-loop-nov (red diamond), *div*-nov (green diamond), and *codir*-loop-scn (orange circle), were the highest among other conditions, placing their native supercoiling levels midway on the bell curve. Interestingly, the bell curves estimated differential native supercoiling levels of gene pairs *G2-G3* and *G1-G3* compared to that of the *G1-G2* pair even under identical unlooped conditions (**Table S10, Fig. S12**), suggesting that each gene may experience a unique, local supercoiling density related to its own syntax and differential coupling with neighboring genes, in contrast to the common assumption that the same supercoiling level rapidly equilibrates and is commonly shared by all genes on the same chromosomal DNA.

## Discussion

In this work, we investigated how chromosomal topological domain modulates transcription in *E. coli*. Using a controllable domain-formation approach that alters the chromosomal domain topology in the native cellular environment without altering gene or promoter sequences, combined with single-cell, single-molecule RNA quantification and modeling, we demonstrated that topological domain formation significantly impacts transcription levels of individual genes and modulates gene-gene coupling. Our results establish that chromosome topology is a major transcription regulator, independent of protein transcription factors.

We observed that domain formation consistently repressed transcription of genes enclosed within the domain (*G1* and *G2*), regardless of their relative orientations (**Fig. 1C-F**). These results are consistent with the model in which torsional stress accumulates within topologically constrained regions, reducing promoter accessibility or preventing RNAP escape from the initiation complex.^47^ Interestingly, *G3*, which is located outside the domain, also showed repressed transcription upon domain formation, suggesting that the formation of a new topological domain boundary could influence neighboring transcription units, regardless of whether it is part of an existing topological domain. As *G3* is separated from the nearest domain boundary (*O_R_* operator site) by nearly 600 bp, the repression effect is most likely mediated by accumulated negative DNA supercoils between the promoter and the boundary, which can propagate on the DNA through long distances, as previously observed within a topological domain.^35^

Perturbations to DNA supercoiling levels provided further insight into the functional consequences of torsional stress. Inhibition of gyrase with novobiocin, which increases positive supercoiling, led to pronounced repression of *G3* transcription compared to that of *G1* and *G2,* regardless of the domain formation (**Fig. 2A, B**). This repression is likely because *G3* is the most highly transcribed gene among the three, which requires significantly higher gyrase activities to remove positive supercoils generated by its transcription. However, topological domain formation appears to exacerbate the repressive effect of gyrase inhibition on *G1* and *G2*, suggesting that topological domains enhance gene sensitivity to accumulated supercoils (**Fig. 2A, B**).

In contrast, inhibition of Topoisomerase I with seconeolitsine, which leads to increased negative supercoiling, produced context-dependent outcomes. The divergent construct showed seconeolitsine broadly enhancing transcription regardless of looping (**Fig. 2C, D**). This result can be related to the established activation of divergent promoters by transcription-induced supercoils in *topA*-deficient strains.^78^ The codirectional construct showed repressed transcription in the absence of the domain formation but enhanced transcription in the presence of the domain (**Fig. 2C, D**). This inversion suggests that additional negative supercoils can either restore or hamper promoter accessibility, depending on the gene’s syntax.

Analyzing the correlation between the percentage transcription change of each construct after drug treatment and its basal expression level revealed that accumulated positive supercoils repressed transcription, whereas accumulated negative supercoils had a homogenizing effect, repressing highly expressed genes while activating lowly expressed genes (**Fig. 2E, F**). It is possible that highly expressed genes generate a high level of negative supercoils behind RNAP in a naturally negatively supercoiled chromosomal environment; thus, further accumulation of negative supercoils caused by Topo I inhibition represses transcription, likely by trapping RNAP at promoters. Lowly expressed genes, especially under the looped, constrained conditions where genes are insulated from the naturally negatively supercoiled chromosomal DNA, required more negative supercoils to be activated. These observations support a model in which positive and negative supercoils exert differential regulatory effects, shaped by domain architecture and gene orientation.

Beyond modulating the transcriptional output of individual genes and their responses to global supercoiling levels, topological domain formation also impacts transcription coupling between neighboring genes in complex ways. Despite strong apparent correlations between the expressed RNA copy numbers of different genes (**Table S9**), the presence of extrinsic noise (e.g., due to fluctuations in RNAP levels)^71–73^, prevented us from isolating the intrinsic correlations that resulted solely from the transcription coupling between genes. To address this problem, we leveraged the observation that distinct RNA puncta containing multiple RNA molecules most likely represent nascent transcription sites on the chromosome (**Fig. 3B, Fig. S4**). We also noted that long cells (>2.0 µm) contain well-separated transcription sites in the two halves of the cell, most likely corresponding to two replicated chromosomes (**Fig. 3C**). Based on these observations, we developed a decomposition analysis to separate the influence of extrinsic noise on individual gene transcription from the correlations between different genes.

We showed that at the individual gene level, intrinsic noise dominated and that topological domain formation increased intrinsic noise (**Fig. 3E**), most likely due to the repressed expression levels (**Fig. S8**). The intrinsic Fano factors of the three genes were all significantly larger than one (**Fig. 3F**), indicating a non-Poissonian transcription process and most consistent with transcriptional bursting. Here, topological domain formation did not appear to alter the intrinsic transcription mode, suggesting that transcription bursting may be a common transcription mechanism in bacteria.

Extending our decomposition analysis to the expression of genes on the same chromosome (**Fig. 4A**), we isolated the intrinsic and extrinsic covariances between any two genes in their transcribed RNA copy numbers. Here, intrinsic covariance refers to the internal coupling of two genes due to their shared chromosomal DNA state, whereas extrinsic covariance refers to external factors, such as RNAP levels and cellular resources, that are shared by genes in the same cells. We observed that extrinsic covariances dominated across all conditions, but significant intrinsic covariances persisted (**Fig. 4B**). In most conditions, transcription of a gene pair was positively correlated (positive intrinsic covariances), largely independent of topological domain formation. However, *G1* and *G2* showed significant negative covariances in two opposing conditions, *codir-*loop-nov and *div*-loop-scn, which, when compared to the positive covariances under the corresponding unlooped conditions, suggest that topological domain formation indeed impacts gene-gene coupling in a syntax-dependent manner, here possibly due to different gene orientations.

What could lead to the observed intrinsic correlations or anticorrelations between different genes? We reasoned that transcription-generated supercoils, along with the native supercoiling state of the shared chromosomal DNA, could create different local supercoiling environments with varying DNA mechanics, through which neighboring genes modulate each other’s transcription. This possibility is supported by the observation of significant intrinsic covariances among all the three gene pairs including the *G1-G3* pair, which were nearby but not immediately adjacent to each other on the chromosome like the *G1-G2* and *G2-G3* pairs (**Fig. 4B**). As such, one gene’s transcription could influence another gene’s transcription in a context-dependent manner, creating complex transcription dynamics independent of protein transcription factors. For example, one gene’s switching-on rate could depend on the on- or off-state of a neighboring gene, and *vice versa*, because the local supercoiling shared by the two genes may differ under different combinations of expression states.

To understand quantitatively how neighboring genes modulate each other’s transcription, we developed a minimal theoretical model to capture the joint distributions of RNA copy numbers for gene pairs across all conditions (**Fig. 5A-D**). The model showed that the overall coupling strength between gene pairs is independent of their expression levels but instead dependent on gene syntax (**Fig. S11**). Interestingly, for the same construct, shifting the global negative supercoiling levels from high (treated with seconeolitsine) to native (untreated) and to low levels (treated with novobiocin) produced varied, sometimes opposing, and non-monotonic responses (**Fig. 5E, Fig. S12**). Based on these observations, we proposed a model in which gene-gene coupling exhibits a non-monotonic dependence on supercoiling levels, with extremely high or low negative supercoils reducing coupling (**Fig. 5F**), just as extreme positive and negative supercoils decrease the mean expression levels. Importantly, this model also indicated that each construct’s local supercoiling density is modulated by its unique gene syntax (**Fig. 5F, Fig. S12**), in contrast to the commonly assumed rapid diffusion and equilibration of supercoils on chromosomal DNA. It is possible that supercoils along the DNA are maintained locally between any two transcribing RNAP molecules, which form topological barriers, but are not uniformly shared along the DNA. These local supercoils, when combined with stochastic transcription initiation, gene orientations, and topological domain formation, gave rise to complex gene-gene coupling behaviors.

One limitation of our study is that, from the steady-state RNA distributions measured by smFISH, we could only determine the overall coupling strength but not its directionality (see STAR Methods and **Fig. S10B**). Our model predicts that the directionality of coupling would lead to asymmetries in temporal correlations (**Fig. S10B**). Therefore, future experiments measuring the dynamics and temporal correlation of gene expression would help determine the dominant direction of the gene-gene coupling, which would provide additional information about the microscopic mechanisms underlying supercoiling-mediated gene-gene coupling. Such measurements may be explored using the DuTrAC single-molecule gene expression reporters that we developed previously.^79^ The temporal information would reveal more information about the correlations and coupling mechanisms present in topologically interacting genes and key parameters for expanding our model.

Another limitation of our study is that we did not account for the effects of co-transcriptional translation. As all three genes contained ribosomal binding sites, mRNA-associated polysomes during co-transcriptional translation likely contributed significantly to the observed supercoiling sensitivity of the genes due to the additional torsional stress. Thus, translation could also alter transcription dynamics through the act of topological constraint. This possibility will be explored in future work.

Finally, despite multiple attempts, we were not able to obtain a convergent chromosomal construct in which *G1* and *G2* transcribe toward each other. We suspected that the convergent arrangement of *G1* and *G2* at their expression levels may create a high level of chromosomal DNA torsional stress due to the accumulated positive supercoils in between^22,80^, leading to chromosomal instability.^81,82^

In summary, our findings established that chromosomal topology and DNA supercoiling constitute an intrinsic mechanical layer of transcriptional regulation. Topological domains can modulate gene expression levels and couple neighboring genes, providing a physical mechanism for context-dependent coordination across the chromosome. These results highlight how mechanical effects and constraints work together with canonical regulatory networks to control transcription in complex chromosomal environments.

