## Supplemental File for "Chromosomal Topological Domain Formation Modulates Transcription and the Coupling of Neighboring Genes in *Escherichia coli*"

### STAR Methods

| **Key resources table** | | |
| --- | --- | --- |
| **REAGENT or RESOURCE** | **SOURCE** | **IDENTIFIER** |
| **Bacterial Strains** | | |
| See **Table S2** for bacterial strains | N/A | N/A |
| **Chemicals, peptides, and recombinant proteins** | | |
| Kanamycin sulfate salt | Sigma Aldrich | Cat #: 60615-5G; CAS: 70560-51-9 |
| Streptomycin sulfate salt | Sigma Aldrich | Cat #: S9137-25G; CAS: 3810-74-0 |
| Chloramphenicol | Fisher Scientific | Cat #: BP904-100; CAS: 56-75-7 |
| Tetracycline | Sigma Aldrich | Cat #: 87128-100G; CAS: 60-54-8 |
| Novobiocin sodium salt | Sigma Aldrich | Cat #: N1628-1G; CAS: 1476-53-5 |
| Seconeolitsine | Universite de Lyon, INSA Lyon | Labs of Florence Popowycz and Laurent Soulère^1^ |
| EZRDM - MOPS EZ Rich Defined Medium Kit | Teknova | Cat #: M2105 |
| IPTG, dioxane-free | Thermo Scientific | Cat #: R0392; CAS: 367-93-1 |
| L-(+)-Arabinose | Sigma Aldrich | Cat #: A3256-25G; CAS: 5328-37-0 |
| PBS (1x), pH 7.4 | Quality Biological | Cat #: 114-058-101 |
| Formaldehyde Solution | Sigma Aldrich | Cat #: FX0410-5 |
| UltraPure™ 20x SSC Buffer | Invitrogen | Cat #: 15557-044 |
| Tris Base | Fisher Scientific | Cat #: BP152-500 |
| Acetic Acid | Fisher Scientific | Cat #: A38SI-212 |
| Potassium-Acetate | Sigma Aldrich | Cat #: 236497-2.5KG |
| Magnesium-Acetate | MD Chemicals | Cat #: MX0025-1 |
| DTT | GoldBio | Cat #: DTT50 |
| Nt.BbvCI | New England Biolabs | Cat #: R0632L |
| T4 DNA ligase | Florida International University | Lab of Fenfei Leng |
| ATP | Amersham Pharmacia Biotech, Inc | Cat #: 27-1006-01 |
| λ CI | Clemson University | Lab of Laura Finzi^2^ |
| **Oligonucleotides** | | |
| smFISH probes, see **Table S3** | This study (Stellaris, LGC Biosearch Technologies) | N/A |
| 5′CGGCATGGCGGCCCTATGCTGAGGACTCGGCCGACGCGCT-3′ | Eurofins Genomics | A double-stranded oligonucleotides containing the Nt.BbvCI recognition site |
| **Software and Algorithms** | | |
| MATLAB (2024b) | Mathworks | <https://www.mathworks.com/products/matlab.html> |
| Mathematica 13.1 | Wolfram | [https://www.wolfram.com/mathematica/](https://nam02.safelinks.protection.outlook.com/?url=https%3A%2F%2Fwww.wolfram.com%2Fmathematica%2F&data=05%7C02%7Cxiao%40jhmi.edu%7Cb7dc1b3dbdb84ec79f3d08de2704a906%7C9fa4f438b1e6473b803f86f8aedf0dec%7C0%7C0%7C638991098189054507%7CUnknown%7CTWFpbGZsb3d8eyJFbXB0eU1hcGkiOnRydWUsIlYiOiIwLjAuMDAwMCIsIlAiOiJXaW4zMiIsIkFOIjoiTWFpbCIsIldUIjoyfQ%3D%3D%7C0%7C%7C%7C&sdata=nAt0k2KkQXzFlVbICkChVmSLky%2FSZ%2FAnTAcOZ9yLIDA%3D&reserved=0) |
| Python 3.12.2 | Python Software Foundation | <https://www.python.org/downloads/release/python-3122/> |

### Experimental Methods

#### Bacterial strains and plasmids construction

Strains NY45 (codirectional construct) and NY89 (divergent construct) were constructed from the parental strain XF001 that contains a landing pad at the *lac* operon site on the chromosome.^3^ The landing pad method^4^ was used to insert large constructs (> 1 Kbp) into the chromosomal DNA. Specifically, the codirectional construct donor plasmid (pSHAY3a) and the divergent construct donor plasmid (pSHAY5a) were first constructed using In-Fusion cloning (Takara). Plasmid pSHAY3a or pSHAY5a was electroporated into the XF001 strain containing the helper plasmid pTKRED (From Dr. Thomas E. Kuhlman). Several small cultures of the XF001 strain containing pTKRED and pSHAY3a or pSHAY5a were grown in 1μL of Lysogeny Broth (LB) at 30°C for 4 hours. These small cultures were supplemented with 200μL of EZRDM rich media with 50 μg ml^-1^ Streptomycin and grown again at 30°C for 4 hours. Finally, the Landing Pad reaction was induced by adding 2mL EZRDM media containing 2 mM IPTG, 0.4% Arabinose, and 10 μg ml^-1^ Kanamycin to each culture, and the cultures were allowed to grow overnight at 30°C. The following morning, the cultures were diluted 1:1000 in LB media; 100 μL of each dilution was plated on LB plates containing 50 μg ml^-1^ Kanamycin and incubated overnight at 37°C).

Colonies grown on overnight plates were tested for insertion by colony PCR and whole-genome sequencing. Once colonies with successful insertion of the construct were identified, the strains were cured of the pTKRED plasmid by growing cultures in 3 mL of LB with 10 μg mL^-1^ Kanamycin at 42°C for 7 hours and diluting 1:3000 into 3 mL of LB with 10 μg mL^-1^ Kanamycin to grow at 42°C overnight.

The resulting strains, NY45 (codirectional construct – pSHAY3a) and NY89 (divergent construct – pSHAY5a), were further verified by sequencing and then transformed with a plasmid pACL18, which expresses the CI λ-repressor gene under a constitutive promoter, to produce strains NY50 and NY91, respectively. As a control, NY45 and NY89 were also transformed with the plasmid pACL18noCI, which was constructed by removing the CI λ-repressor gene from pACL18 via In-Fusion cloning (Takara), to produce strains NY51 and NY92, respectively. Strains NY50, NY51, NY91, and NY92 were used for smFISH experiments.

#### DNA Nicking Topology Assay

The DNA topology nicking assay plasmid pNY1 was cloned from plasmid pSHAY3a, where a BbvCI nick site was added to the portion of the plasmid that lies opposite the *G1*/*G2* loop. Assays were performed in a mixture (320 μL) containing 0.156 nM of negatively supercoiled DNA template and 170 nM of wild-type λ repressor (CI) in 20 mM Tris-acetate (pH 7.9 at 25 °C), 50 mM potassium acetate, 10 mM magnesium acetate, and 1 mM DTT was assembled on ice and incubated for 30 min at 37 °C. Then, the supercoiled DNA templates were digested by 20 units of Nt.BbvCI at 37 °C for 15 s. Next, a large excess (289 nM) of a double-stranded oligonucleotides containing the Nt.BbvCI recognition site (top strand, 5′ CGGCATGGCGGCCCTATGCTGAGGACTCGGCCGACGCGCT-3′) was added to the reaction mixtures to inhibit further plasmid digestion. The nicked DNA templates were ligated using 50 nM of T4 DNA ligase in the presence of 1 mM ATP at 37 °C for 5 min, and the reaction was terminated by extraction with an equal volume of phenol. Plasmids were precipitated in ethanol and dissolved in 25 μL of 10 mM Tris·HCl buffer (pH 8.5). The ligated DNA products were separated using 0.8% agarose gel electrophoresis, and the superhelicity of topoisomers was calculated from the gel images stained with ethidium bromide or SYBR Green.

**Single Molecule Fluorescent *in situ* Hybridization (smFISH)**

All DNA oligos used for smFISH were ordered from Biosearch Tech: *lacZ* (Gene 1) transcripts were labeled with 48 oligonucleotides conjugated with Fluorescein Dye, *luc2* (Gene 2) transcripts were labeled with 48 oligonucleotides conjugated with CAL Fluor Red 590 Dye, and *NPTII* (Gene 3) transcripts were labeled with 30 oligonucleotides conjugated with Quasar 670 Dye. See **Table S3** for probe sequences.

Cultures of NY50 and NY51 were grown overnight in LB media with 25 μg ml^-1^ Chloramphenicol at a temperature of 30°C while shaking. Control cultures of XF001 were grown overnight in 3mL LB media with 6μL Tetracycline at a temperature of 30°C while shaking. The following morning, cultures were diluted to an optical density (OD) below 0.4 and allowed to grow until the OD reached 0.5. Once an OD of 0.5 was reached, the cultures were treated with either 300 μg ml^-1^ (489.7 μM) novobiocin or 25 μM seconeolitsine for 15 minutes before fixation. Cultures were fixed in a 3.7% (vol/vol) formaldehyde solution for 30 minutes. As a wash step, samples were spun down and resuspended in 1x PBS 2 times. The fixed cells were permeabilized in 70% (vol/vol) ethanol for 1 hour. Samples were spun down and resuspended in 1x PBS 1 time. The samples were spun down again, resuspended in hybridization buffer (40% (wt/vol) formamide, 2x SSC) containing the oligonucleotide probes, and incubated overnight. The following day, samples were spun down, resuspended in 1x PBS 2 times, and stored on ice. This procedure was adapted from Skinner *et al*.^5^

Fixed and labeled cells were imaged using brightfield conditions and cycles of excitation at 488 nm (Coherent, obis), 561 nm (Coherent, sapphire), and 647 nm (Coherent, obis). Fluorescent images were collected at exposure times of either 200 ms or 100mes with a power density of approximately 200 W/cm^2^. Each field of view was imaged at four *z*-planes evenly spaced by 250nm.

**FISH Data Analysis**

The maximum projection of the four z-stack images for each field of view was used to count individual RNA molecules in each cell. Corrections were applied for registration between fluorescence channels, photobleaching between subsequent images, and saturation correction as previously described^6^. Spots were detected as local maxima and discriminated using a negative control image set (target genes absent) and shape/symmetry parameters. The total fluorescence intensity of each spot in the images was divided by the intensity of a single mRNA molecule, and the result was rounded to calculate the RNA copy number in that spot. The intensity of single RNA molecule for each imaging channel was determined by a multi-Gaussian fit of the fluorescent spot intensity distribution, as done in other FISH protocols.^5^ Image analysis was done using custom MATLAB scripts.

#### Statistical analysis of RNA expression levels

The distributions of RNA copy number per cell are non-normally distributed as shown in **Fig. S7**. Therefore, we applied a two-sample Kolmogorov-Smirnov test using the built-in MATLAB function kstest2. The results were reported in **Table S5**. Similarly, we used the built-in MATLAB function anova to analyze the three-way interactions among looping state, construct orientation, and drug treatment for each RNA’s expression. **Table S6** lists the ANOVA analysis results. We reported both the *F* statistics and *p-values*, but did not interpret the *p-values because* the data were not normally distributed. Instead, we relied on the *F* values to determine the most significant factor.

#### Diffusion of RNA with degradation

Diffusion of an mRNA molecule with coefficient $D$:

$$P\left( \vec{x},t \right)=\frac{1}{\sqrt{\left( 4\pi Dt \right)^{3}}}\exp\left( -\frac{\vec{x}.\vec{x}}{4Dt} \right)$$

resulting in a mean squared displacement:

$$\langle\vec{x}\left( t \right)^{2}\rangle=6Dt$$

We now include an exponential lifetime distribution of parameter $\lambda$:

$$P\left( \tau\right)=\lambda e^{-\lambda\tau}$$

with $\lambda=1/T=log\left( 2 \right) t_{1/2}$. For a typical mRNA of $t_{1/2}=60 s$, we have $\lambda\simeq0.7/60\simeq0.01 s^{-1}$.

Mean displacement away from the transcription site accomplished by an mRNA molecule before degradation:

$$\langle r^{2}\rangle=\int_{\tau=0}^{\infty} P\left( \tau\right)\langle\vec{x}\left( \tau\right)^{2}\rangle d\tau=\int_{\tau=0}^{\infty} \lambda e^{-\lambda\tau} 6D\tau d\tau$$

$$\boxed{\langle r^{2}\rangle=\frac{6D}{\lambda}}$$

For the degradation rate $\lambda$ above, and a diffusion coefficient $D={2.10}^{-5} \mu m^{2} . s^{-1}$, each RNA diffuses to a mean distance $\langle r\rangle\simeq0.1 \mu m$, and all RNAs produced from a locus (assuming no overall movement) appear as a single punctum if this is the spatial resolution. If the diffusion coefficient is much larger, RNAs quickly diffuse to the entire cell and no multi-RNA puncta are visible.

Assuming a transcription with bursts of size $b$ and frequency $f$ (thus an average time between bursts of $1/f$), the average number $\langle n\rangle$ of RNAs within the punctum is

$$\langle n\rangle=\frac{bf}{\lambda}$$

Poissonian expression corresponds to $b=1$ where $f$ is the transcription initiation rate. The mode of expression does not affect the diffusion distance, but changes the distribution of mRNA counts within puncta.

#### Master equation for the four-state model

Here, we provide further detail on a general four-state model describing the correlated expression of two genes (see **Fig. 5A** and the associated main text). The model has 12 parameters, including two production rates $\mu_{i}$, two degradation rates $d_{i}$, and eight gene state switching rates $k_{on,i}$, $k_{off,i}$, $k_{\text{on},i}^{'}$, and $k_{\text{off},i}^{'}$.

The system is described by a joint distribution of gene states and mRNA counts $P(s_{1},s_{2},m_{1},m_{2},t)$. $s_{i}=0,1$ are binary variables describing gene $i$’s on/off state at time *t*. $m_{i}$ is the number of transcripts for gene $i$. The joint distribution follows dynamics governed by the following master equation

$$\begin{aligned} \begin{matrix} \frac{dP\left( s_{1},s_{2},m_{1},m_{2},t \right)}{dt}=\left( \mathcal{L}_{\mathrm{switch}}+\mathcal{L}_{\mathrm{mRNA}} \right)P\left( s_{1},s_{2},m_{1},m_{2},t \right), \end{matrix}\#\left( 1 \right) \end{aligned}$$

where $\mathcal{L}_{\mathrm{switch}}$ describes the switching dynamics of gene states, and $\mathcal{L}_{\mathrm{mRNA}}$ describes the mRNA production and degradation. The production/degradation dynamics $\mathcal{L}_{\mathrm{mRNA}}$ are given by

$$\begin{aligned} \mathcal{L}_{\mathrm{mRNA}}P(s_{1},s_{2},m_{1},m_{2},t)=&s_{1}\mu_{1}P(s_{1},s_{2},m_{1}-1,m_{2},t)+s_{2}\mu_{2}P(s_{1},s_{2},m_{1},m_{2}-1,t) \\ &+d_{1}(m_{1}+1)P(s_{1},s_{2},m_{1}+1,m_{2},t) \\ &+d_{2}(m_{2}+1)P(s_{1},s_{2},m_{1},m_{2}+1,t) \\ &-(s_{1}\mu_{1}+s_{2}\mu_{2}+d_{1}m_{1}+d_{2}m_{2})P(s_{1},s_{2},m_{1},m_{2},t). \end{aligned}$$

Since there is no feedback from mRNA production to gene-state switching, the switching dynamics depend only on $(s_{1},s_{2})$ and not on $(m_{1},m_{2})$. The switching dynamics $\mathcal{L}_{\mathrm{switch}}P(s_{1},s_{2},t)$ are given by

$$\begin{aligned} \begin{aligned} \mathcal{L}_{\mathrm{switch}}P\left( 0,0,t \right)=k_{off,1}P\left( 1,0,t \right)+k_{off,2}P\left( 0,1,t \right)-\left( k_{on,1}+k_{on,2} \right)P\left( 0,0,t \right), \\ \mathcal{L}_{\mathrm{switch}}P\left( 1,0,t \right)=k_{on,1}P\left( 0,0,t \right)+k_{\text{off},2}^{'}P\left( 1,1,t \right)-\left( k_{off,1}+k_{\text{on},2}^{'} \right)P\left( 1,0,t \right), \\ \mathcal{L}_{\mathrm{switch}}P\left( 0,1,t \right)=k_{on,2}P\left( 0,0,t \right)+k_{\text{off},1}^{'}P\left( 1,1,t \right)-\left( k_{off,2}+k_{\text{on},1}^{'} \right)P\left( 0,1,t \right), \\ \mathcal{L}_{\mathrm{switch}}P\left( 1,1,t \right)=k_{\text{on},1}^{'}P\left( 0,1,t \right)+k_{\text{on},2}^{'}P\left( 1,0,t \right)-\left( k_{\text{off},1}^{'}+k_{\text{off},2}^{'} \right)P\left( 1,1,t \right). \end{aligned}\#\left( 2 \right) \end{aligned}$$

Solving the master equation $dP(s_{1},s_{2},m_{1},m_{2},t)/dt=0$ gives the steady-state distribution $P(s_{1},s_{2},m_{1},m_{2})$, which can be used to compute the joint distribution $P(m_{1},m_{2})$ of mRNA counts by marginalizing over the gene states:

$$\begin{matrix} P(m_{1},m_{2})=\sum_{s_{1},s_{2}=0,1} P(s_{1},s_{2},m_{1},m_{2}) \end{matrix}$$

The experimental measurements in our study represent independent samples $(m_{1},m_{2})$ from this steady-state distribution.

### Model reduction

For individual genes, we find that the mRNA count distribution $P\left( m_{i} \right)$ can be captured by a negative binomial distribution (see **Fig. S7**) which is consistent with the bursting limit of the telegraph model (PMID: 38594901), where the gene is off for the majority of the time, but when it is on, it produces bursts of multiple transcripts. This bursting limit is achieved when there is a separation of timescales: $k_{on,i},d_{i}\ll k_{off,i},\mu_{i}$. The slow processes are mRNA degradation ($d_{i}$) and gene switching from off to on ($k_{on,i}$). Their rate ratio is defined as the burst frequency $\gamma_{i}=k_{on,i}/d_{i}$, which is the average number of bursts observed during the decay time of a transcript. The fast processes are transcription ($\mu_{i}$) and gene switching from on to off ($k_{off,i}$). Their rate ratio is defined by the burst size $\beta_{i}=\mu_{i}/k_{off,i}$, which is the average number of transcripts produced in a burst before the gene is turned off. Together, in the limit of timescale separation $\tau_{i}=k_{off,i}/d_{i}\gg1$, the steady-state distribution is determined by bursting frequency $\gamma$ and size $\beta$.

When our four-state model (**Fig. 5A**) operates in the bursting limit, the individual switching rates $k_{on,i}$ and $k_{off,i}$ (i.e., when the other gene is off) obey timescale separation as described above. However, $k_{\mathrm{on},i}^{'}$ (the on rate when ohe other gene is also on) can range from order ${O(k}_{on,i})$ when there is almost no interaction to order ${O(k}_{off,i})$ when the inter-gene coupling is substantial.

In the bursting limit, the distribution is insensitive to the exact value of $\tau_{i}$ as long as it is large. Hence, we can simplify the four-state model by assuming $\tau_{1}=\tau_{2}=10\gg1$ without affecting the quality of fits (larger values of $\tau_{i}$ also yield similar results). We also adopt equal degradation rates $d_{1}=d_{2}=1$ to further reduce the number of parameters and set the unit of time. These choices are not necessary for our model to capture the data but are introduced to prevent overfitting; the model can similarly fit the data with other choices of $\tau_{i}$ and $d_{i}$. As further described in the Discussion, measurements of gene expression dynamics will allow for a more precise determination of all model time scales.

The general four-state model can capture a wide range of interactions from strong activation to strong inhibition. However, in the bursting limit mutual, it is easier for the model to capture activating interaction than inhibitory interaction. This is because simultaneous activation of two uncoupled genes is already very rare, of order $O(1/k_{\mathrm{off}}^{2})$, so suppressing double activation has no discernible effect on the mRNA distribution. Since the data also show predominantly positive correlations, we restrict the model to activation-enhancing interactions, which we assume to modulate the on rates ($k_{\text{on},i}^{'}>k_{on,i}$) without affecting the off rates ($k_{off,i}=k_{\text{off},i}^{'}=\tau_{i}d_{i}$).

With these assumptions, we arrive at a reduced model with 6 parameters: two production rates $\mu_{i}$ and four on rates $k_{on,i}$ (when the other gene is off) and $k_{\text{on},i}^{'}$(when the other gene is on).

### Data selection and Model fitting

Our dataset contains cells at various cell cycle stages, including those with replicated chromosomes. Since our model predicts the mRNA counts for a single copy of each gene, we filter the data to isolate short cells (<1.7 μm), which most likely has only a single chromosome copy based on our RNA puncta measurements (**Figs. 3C, S5, S6**). Restricting to this population of cells, the measured mRNA counts correspond to independent samples of the steady-state distribution $P(m_{1},m_{2})$ predicted by our model.

To fit the model to data, we use a maximum likelihood approach; we minimize the loss function $L(\theta)$ defined to be the negative log likelihood of the data given model parameters $\theta$:

$$\begin{aligned} \begin{aligned} L\left( \theta\right)=-\sum_{i=1}^{N} \ln P\left( \text{data}_{i} | \theta\right), \end{aligned}\#\left( 3 \right) \end{aligned}$$

where $i$ sums over cells, and $P$ is the steady-state transcript distribution $P(m_{1},m_{2})$ computed from the model above. We minimize this loss function with respect to model parameters $\theta$ using the L-BFGS-B algorithm implemented in jaxopt 0.8.5. The uncertainty of fitting parameters is obtained by computing the inverse Hessian of the negative log likelihood using jax automatic differentiation. As noted above, we restrict the fits to activation-enhancing interactions $k_{\text{on},i}^{'}>k_{on,i}$, since inhibitory interactions produce a distribution that is identical to a model with constant on rates $k_{\text{on},i}^{'}=k_{on,i}$.

The simplified 6-parameter model fits very well with the joint distribution $P(m_{1},m_{2})$ and low order moments across all twelve experiment conditions, including 4 constructs (codirectional, codirectional-loop, divergent, divergent-loop) across 3 treatments (native, novobiocin, seconeolitsine), see **Fig. 5, S10** for the *G1-G2* pair, and **Fig. S12** for the *G2-G3* and *G1-G3* pairs.

### Amplification ratio $\boldsymbol{\Gamma}$ and temporal correlations

As mentioned in the Discussion, a key question about the coupling between neighboring genes on the DNA is the directionality of the interaction: does *G1* have more influence on *G2* or vice versa? Our model contains an intuitive measure for the dominant interaction direction, $\Gamma=\alpha_{1}/\alpha_{2}$, where $\alpha_{i}=k_{\text{on},i}^{'}/k_{on,i}$ is the fold amplification of the of the gene $i$ on-rate while the other gene is active. For example, $\Gamma\neq1$ indicates an asymmetry in the interaction, while $\Gamma=1$ indicates a symmetric interaction. An asymmetry might arise if, for example, activating *G1* strongly enhances the activation rate of *G2*, but the state of *G2* only weakly enhances *G1* dynamics ($\Gamma<1$ in this scenario). The amplification ratio $\Gamma$ also has a thermodynamic interpretation: $k_{B}Tln\Gamma$ is the free energy dissipated per activation/deactivation cycle through the four gene states, with $\Gamma=1$ corresponding to equilibrium, non-dissipative, dynamics.

The parameters in our model are well constrained by the measured data with one primary exception: we can vary $\Gamma$ within a moderate range and achieve comparable fits. To demonstrate this property, we fit the measured distributions as described in the preceding section with an additional constraint fixing the value of $\Gamma$. As shown in **Fig. S10B1**, for $-2⪅ln\Gamma⪅2$, the mean square error of the distribution fits (averaged over all conditions) is comparable to that of the fit with unconstrained $\Gamma$. Despite this combination of parameters being somewhat flexible, below we show that our results (in particular, the non-monotonicity of inter-gene coupling with supercoiling) are robust to variations in $\Gamma$. The boundaries of the best fit region have an intuitive physical interpretation: they correspond to an approximately unidirectional interaction, as shown **in Fig. S10B2**. For example, in the best fit with $ln\Gamma=2$, *G2* only effects *G1* and not vice versa: there is a $\alpha_{1}=e^{2}$ fold-enhancement in the *G1* activation rate and no enhancement $\alpha_{2}\approx1$ for *G2*. The opposite sign $ln\Gamma=-2$ corresponds to the same scenario with the directionality of interaction reversed. Since we restrict to activation-enhancing interactions, increasing $|\Gamma|$ further leads to uni-directional interaction with the dominant interaction $\alpha_{i}\approx|\Gamma|$ (**Fig. S10B2**), which produces gene correlations larger than those observed.

As stated in the Discussion, $\Gamma$ is not fully constrained by our fits because the measured data are sampled from a steady-state distribution. We have shown that these measurements are sufficient to constrain the gene interaction, but we cannot resolve whether there is a dominant direction in the interaction (i.e. $\Gamma>1$ or $\Gamma<1$). We proposed that temporal measurements can be used to more precisely determine $\Gamma$ as well as various other timescales underlying gene expression dynamics. Here we provide exact analytical expressions for the temporal correlations predicted by our model. Defining

$$\mathbf{P}(m_{1},m_{2},t)=[P(0,0,m_{1},m_{2},t),P(1,0,m_{1},m_{2},t),P(0,1,m_{1},m_{2},t),P(1,1,m_{1},m_{2},t)],$$

diagonal production matrices $M_{1}=diag(0,\mu_{1},0,\mu_{1})$ and $M_{2}=diag(0,0,\mu_{2},\mu_{2})$, and $4\times4$ gene switching matrix $\Omega_{\mathrm{switch}}$ containing the transitions in Eq. (2), we can solve the master equation, Eq. (1) for the mRNA count autocorrelation and cross-correlation using an approach similar to (PMID: 38594901, Appendix E). We find that the autocorrelation for gene $i$, $C_{ii}(t)=\langle m_{i}(t_{0})m_{i}(t_{0}+t)\rangle_{t_{0}}-\langle m_{i}^{2}\rangle$, is given by:

$$C_{ii}(t)=\left[ e^{-d_{i}t}-1 \right]\langle m_{i}^{2}\rangle+\mathbf{1}^{\top}D_{i}\left[ e^{\Omega_{\mathrm{switch}}t}-e^{-d_{i}t} \right](d_{i}^{2}-\Omega_{\mathrm{switch}}^{2})^{-1}D_{i}\mathbf{P}_{0},$$

while the temporal cross-correlation is $C_{ij}(t)=\langle m_{i}(t_{0})m_{j}(t_{0}+t)\rangle_{t_{0}}-\langle m_{i}\rangle\langle m_{j}\rangle$, is given by:

$$C_{ij}(t)=e^{-d_{i}t}\langle m_{i}m_{j}\rangle-\langle m_{i}\rangle\langle m_{j}\rangle+\mathbf{1}^{\top}M_{j}\left[ e^{\Omega_{\mathrm{switch}}t}-e^{-d_{j}t} \right](\Omega_{\mathrm{switch}}+d_{j})^{-1}(d_{i} -\Omega_{\mathrm{switch}})^{-1}M_{i}\mathbf{P}_{0}.$$

Here $\mathbf{P}_{0}$ is the steady-state distribution on the gene-state space, satisfying $\Omega_{\mathrm{switch}}\mathbf{P}_{0}=0$, $\mathbf{1}$ is the vector containing all 1’s, and we have written the expressions in terms of the equal-time moments,

$$\begin{aligned} \langle m_{i}\rangle&=\mathbf{1}^{\top}(d_{i}-\Omega_{\mathrm{switch}})^{-1}M_{i}\mathbf{P}_{0}, \\ \langle m_{i}^{2}\rangle&=2\mathbf{1}^{\top}(2d_{i}-\Omega_{\mathrm{switch}})^{-1}M_{i}(d_{i}-\Omega_{\mathrm{switch}})^{-1}M_{i}\mathbf{P}_{0}+\langle m_{i}\rangle, \\ \langle m_{i}m_{j}\rangle&=\mathbf{1}^{\top}(d_{i}+d_{j}-\Omega_{\mathrm{switch}})^{-1}\left[ M_{j}(d_{i}-\Omega_{\mathrm{switch}})^{-1}M_{i}+M_{i}(d_{j}-\Omega_{\mathrm{switch}})^{-1}M_{j} \right]\mathbf{P}_{0} \end{aligned}$$

The autocorrelation and cross-correlations for G1 and G2 predicted by the model fit to the codir-scn measurements are shown in **Fig. S10B7**. We further find that the correlation asymmetric $\Delta C_{12}(t)=C_{12}(t)-C_{21}(t)$ depends sensitively on $\Gamma$ (**Fig. S10B8**).

### Coupling non-monotonicity and robustness to variations in $\boldsymbol{\Gamma}$

In our model, the covariance of the expression level of the two genes is captured by switching rates that depend on the state of the other gene. As discussed in the main text, we quantify the strength of this gene-gene interaction with interaction parameter $J_{ij}$,

$$\begin{matrix} J_{ij}=\frac{k_{on,i}}{k_{on,i}+k_{\mathrm{off}}}\left( \frac{k_{\text{on},j}^{'}\left( k_{\mathrm{off}}+k_{on,j} \right)}{k_{on,j}\left( k_{\mathrm{off}}+k_{\text{on},j}^{'} \right)}-1 \right). \end{matrix}$$

As shown in the main text **Fig. 5F**, the impact of supercoiling on coupling is construct-dependent. Compared to the native condition, novobiocin treatment increases positive supercoiling by inhibiting gyrase activity and seconeolitsine treatment promotes negative supercoiling accumulation by inhibiting topoisomerase I. However, these treatments can either increase or decrease coupling depending on the construct. We hypothesized that the differing dependence of coupling on drug treatment could be explained by a non-monotonic relationship between supercoiling $\sigma$ and inter-gene coupling $J$, which takes the form of a bell-shaped curve.

To examine this hypothesis, we assume each construct has a native supercoiling level $\sigma=x_{c}$ and that novobiocin and seconeolitsine treatments increase/decrease the supercoiling level to $\sigma=x_{c}+1$ and $\sigma=x_{c}-1$ respectively. We consider a bell curve,

$$\begin{matrix} J(\sigma)=J_{\max}e^{-\frac{\sigma^{2}}{2\sigma_{c}^{2}}}, \end{matrix}$$

where $J_{\max}$ and $\sigma_{c}$ give the height and width respectively. We fit $x_{c}$ for each construct along with $J_{\max}$ and $\sigma_{c}$ (assumed constant across constructs), finding that the data nicely collapsed onto the bell curve. As described further in the main text, these results support a non-monotonic relation between supercoiling and coupling and the hypothesis that gene expressions are only highly correlated when the supercoiling is at an intermediate level, while strongly positive or negative supercoiling leads to weak or negligible correlations.

While we have used the bell curve as an illustrative example, we note that a broader class of peaked curves could be used to capture our fit couplings $J$. For instance, there may be an asymmetry around the peak as supercoiling is increased or decreased and the change in supercoiling due to drug treatments may also be asymmetric. Though our experiments cannot more precisely constrain the supercoiling-coupling relationship, the fit clearly indicates the presence of non-monotonicity and a peak since coupling increases with supercoiling in some constructs but decreases in others.

The coupling non-monotonicity and the collapse onto the bell curve are robust to variations in $\Gamma$ within the best-fit regime. **Fig. S10B5** and **B6** show the dependence of total coupling on drug treatments for each construct and the bell curve collapse for $ln\Gamma=2$. The overall trends we identified are preserved, including (1) whether increasing supercoiling increases or decreases coupling and (2) the relative locations $x_{c}$ of constructs on the bell curve.

The intricate dependence of coupling on construct and supercoiling also holds for the model fit to G1-G3 (**Fig. S11A**) and G2-G3 statistics (**Fig.S11B**), though the fits had larger deviations with the underlying bell curves. The deviations perhaps indicate the complexity of long-range coupling through another gene and/or a topological domain boundary.

### Bounding extrinsic noise using long-cell measurements

Our four-state model captures the correlated statistics of transcription measured in our experiments but makes a simplifying assumption that all cell-to-cell variability comes from the stochasticity of the expression dynamics (intrinsic noise). In reality, cell-cell variability caused, for example, by long-timescale fluctuations of RNAP concentration and other cellular components, may contribute to the observed fluctuations and correlations in mRNA counts, which is termed extrinsic noise. To interrogate how extrinsic noise affects the fitting result, we must first quantify its magnitude and correlation structure. This section presents the tools for estimating the extrinsic noise structure from long cells (> 2 μm), and the next section incorporates it into the four-state model.

For steady-state mRNA distribution $P(m|E)$ depending on extrinsic variables $E$, we can always decompose the total noise (as quantified by the squared coefficient of variation $\mathrm{CV}^{2}(m)=Var(m)/Mean(m)^{2}$) into extrinsic and intrinsic components,

$$\eta_{\mathrm{tot}}^{2}=\mathrm{CV}^{2}\left( m \right)=\frac{\overline{\left\langle m^{2} \right\rangle}-\left( \overline{\left\langle m \right\rangle} \right)^{2}}{\left( \overline{\left\langle m \right\rangle} \right)^{2}}=\underset{\eta_{\mathrm{int}}^{2}}{\underbrace{\frac{\overline{\langle m^{2}\rangle-\langle m\rangle^{2}}}{\left( \overline{\left\langle m \right\rangle} \right)^{2}}}}+\underset{\eta_{\mathrm{ext}}^{2}}{\underbrace{\frac{\overline{\langle m\rangle^{2}}-\left( \overline{\left\langle m \right\rangle} \right)^{2}}{\left( \overline{\left\langle m \right\rangle} \right)^{2}}}}.$$

Here, the brackets $\langle X\rangle$ denote an average over the steady-state distribution with extrinsic variables $E$ fixed, while the overbar $\overline{X}$ is the average over the extrinsic variability distribution $P(E)$.

Swain et al. showed that the extrinsic-intrinsic decomposition can be computed using two independent samples of expression within each cell,

$$\eta_{\mathrm{int}}^{2}=\frac{\overline{\langle\left( m^{\left( a \right)}-m^{\left( b \right)} \right)^{2}\rangle}}{2\left( \overline{\langle m\rangle} \right)^{2}}, \eta_{\mathrm{ext}}^{2}=\frac{\overline{\langle m^{\left( a \right)}m^{\left( b \right)}\rangle}-\overline{\langle m^{\left( a \right)}\rangle} \overline{\langle m^{\left( b \right)}\rangle}}{\left( \overline{\langle m\rangle} \right)^{2}},$$

where the superscript *a* or *b* indicates the two samples taken within each cell.^7^

We find that a similar decomposition exists for the covariance between two genes, $m_{1}$ and $m_{2}$: there are both extrinsic and intrinsic components as follows,

$$\mathrm{cov}\left( m_{1},m_{2} \right)=\overline{\left\langle m_{1}m_{2} \right\rangle}-\overline{\left\langle m_{1} \right\rangle} \overline{\left\langle m_{2} \right\rangle}=\underset{\mathrm{cov}_{\mathrm{int}}\left( m_{1},m_{2} \right)}{\underbrace{\left( \overline{\left\langle m_{1}m_{2} \right\rangle-\left\langle m_{1} \right\rangle\left\langle m_{2} \right\rangle} \right)}}+\underset{\mathrm{cov}_{\mathrm{ext}}\left( m_{1},m_{2} \right)}{\underbrace{\left( \overline{\left\langle m_{1} \right\rangle\left\langle m_{2} \right\rangle}-\overline{\left\langle m_{1} \right\rangle} \overline{\left\langle m_{2} \right\rangle} \right)}}.$$

Once again we can compute this decomposition in practice using duplicate independent samples from each cell (here subscript denote gene identity while superscript denotes the gene copies on the two sister chromosomes *a* or *b*),

$$\begin{matrix} \mathrm{cov}_{\mathrm{int}}\left( m_{1},m_{2} \right) & =\frac{1}{2}\overline{\langle m_{1}^{\left( a \right)}m_{2}^{\left( a \right)}+m_{1}^{\left( b \right)}m_{2}^{\left( b \right)}-m_{1}^{\left( a \right)}m_{2}^{\left( b \right)}-m_{1}^{\left( b \right)}m_{2}^{\left( a \right)}\rangle} \\ \mathrm{cov}_{\mathrm{ext}}\left( m_{1},m_{2} \right) & =\frac{1}{2}\left[ \overline{\langle m_{1}^{\left( a \right)}m_{2}^{\left( b \right)}\rangle}+\overline{\langle m_{1}^{\left( b \right)}m_{2}^{\left( a \right)}\rangle}-\overline{\langle m_{1}^{\left( a \right)}\rangle} \overline{\langle m_{2}^{\left( b \right)}\rangle}-\overline{\langle m_{1}^{\left( b \right)}\rangle} \overline{\langle m_{2}^{\left( a \right)}\rangle} \right]. \end{matrix}$$

While we do not have a well-controlled measurement of transcription from two independent gene copies in each cell, the two copies of the chromosome in long cells allow us to estimate a bound on the extrinsic contributions to the observed noise and covariance. In particular, the bimodal puncta spatial distribution in long cells suggests that transcripts originating from the two chromosomal copies are minimally mixed. Thus, puncta observed in the two halves of long cells serve as a reasonable proxy for independent samples of gene expression from a single chromosome. While some mixing may occur (transcripts originating on one half of the cell may diffuse to the other half) we find that artificial mixing of the measured transcripts considerably diminished the intrinsic-component of the decomposition, which was in contrast to our experimental observations and further supported the claim that the RNA molecules from the two cell halves were minimally mixed. A further analysis of the extrinsic-intrinsic decomposition indicates that binomial mixing generically increases observed extrinsic noise for a broad range of intrinsic stochastic processes. Therefore, estimation of extrinsic noise from the observed data serves as an upper bound on the true extrinsic noise.

Finally, we used the estimated bounds on extrinsic noise and covariance to investigate the impact of parameter variability on the predictions of our model. We made the simplifying assumption that all extrinsic variability arises from the mRNA production rates $\mu_{1},\mu_{2}$. First note that the means in our four-state model take on the form: $\langle m_{i}\rangle=A_{i}\mu_{i}$ for coefficient $A_{i}$ depending on other parameters $k_{\mathrm{off}},k_{on,i},k_{\text{on},i}^{'},d_{i}$. It follows that the extrinsic noise and (normalized) covariance in mRNA counts directly determines the variability in $\mu_{i}$:

$$\eta_{\mathrm{ext}}^{2}=\frac{\overline{\mu_{i}^{2}}-{\overline{\mu_{i}}}^{2}}{{\overline{\mu_{i}}}^{2}}=\frac{Var(\mu_{i})}{{\overline{\mu_{i}}}^{2}}=\mathrm{CV}_{\mu_{i}}^{2}, \frac{\mathrm{cov}_{\mathrm{ext}}(m_{1},m_{2})}{\overline{\langle m_{i}\rangle}\cdot\overline{\langle m_{j}\rangle}}=\frac{cov(\mu_{i},\mu_{j})}{\overline{\mu_{i}}\cdot\overline{\mu_{j}}}.$$

The next section describes how we incorporated this extrinsic variation in $\mu_{i}$ into our model to study its impact on the observed steady-state distributions and fit parameters.

### Impact of mixing on apparent noise decomposition

In the intrinsic/extrinsic noise and covariance described above, we used the half-cell location of each puncta as a proxy for its chromosomal origin. However, the observed RNA counts may involve diffusive mixing between the two cell halves. Indeed, puncta were observed in near the cell centerline, suggesting that transcripts can diffuse to the other half of the cell within their lifetime. While the extent of mixing cannot be directly quantified from existing experiments, here we develop a theory to study how it affects the intrinsic-extrinsic decomposition and how the mixed statistics relate to the unmixed statistics.

First considering a single gene, let $m^{(L)},m^{(R)}\sim P(m|E)$ be the true transcript counts produced by the two chromosome copies (with extrinsic parameters $E$). The observed transcripts on the left and right sides of the cell, $m_{L},m_{R}$ can be decomposed as follows,

$$m_{L}=m_{L}^{(L)}+m_{L}^{(L)}=m_{L}^{(L)}+m^{(R)}-m_{R}^{(R)} m_{R}=m_{R}^{(L)}+m_{R}^{(R)}=m^{(L)}-m_{L}^{(L)}+m_{R}^{(R)}.$$

In these equations, the subscript represents where the transcript is observed and the superscript represents where it was produced. Assuming random diffusive mixing, we have $m_{L}^{(L)}\sim Binom(m^{(L)},1-p)$ and similar for $m_{R}^{(R)}$. Here $p$ is the probability of the transcript switching sides. With cell length $L$, transcript diffusion constant $D$, and degradation rate $d$, we have that for $\mathcal{l}_{D}=\sqrt{D/d}\ll L/2$, $p\approx0$ and for $\mathcal{l}_{D}\gg L$, $p\approx1/2$.

Averaging over the binomial distribution due to diffusive mixing, the apparent extrinsic and intrinsic noise are

$$\eta_{int,eff}^{2}=\eta_{\mathrm{int}}^{2}-4p(1-p)\left[ \eta_{\mathrm{int}}^{2}-\frac{1}{\overline{\langle m\rangle}} \right] \eta_{ext,eff}^{2}=\eta_{\mathrm{ext}}^{2}+2p(1-p)\left[ \eta_{\mathrm{int}}^{2}-\frac{1}{\overline{\langle m\rangle}} \right].$$

This calculation demonstrates that mixing will increase extrinsic noise and decrease intrinsic noise if the underlying process satisfies $\eta_{\mathrm{int}}^{2}>1/\overline{\langle m\rangle}$, i.e. the process is super-poissonian on average. Thus, computing the intrinsic-extrinsic decomposition on the measured data yields an *upper bound* on extrinsic noise. We note that this bound holds for a broad range of gene expression dynamics, including the MMPP/M/$\infty$ class of queues to which telegraph and our 4-state model belong.^8^ This class of models has production dynamics driven by a Markov Modulated Poisson Process (MMPP) that describes the transitions between gene activation states. These queues are therefore equivalent to Poisson mixtures over the dynamics of the gene-state and hence are always super-Poissonian for any given parameters: $Var(m|E)\geq\langle m|E\rangle\to\overline{\langle m\rangle}^{2}\eta_{\mathrm{int}}^{2}\geq\overline{\langle m\rangle}$. We find that our fit 4-state model is often strongly super-Poissonian, meaning even a small amount of mixing can amplify the apparent extrinsic noise in the data. Thus, the level of extrinsic noise estimated from the long cells form a loose upper bound of the true (unmixed) extrinsic noise. For this reason, we only use half of the maximum extrinsic noise level when fitting the 4-state model to the data.

An analogous calculation considering multiple genes leads to the following mixing-dependence of the covariance,

$$\begin{aligned} \mathrm{cov}_{int,eff}(m_{1},m_{2})&=(1-2p)^{2}\mathrm{cov}_{\mathrm{int}}(m_{1},m_{2}) \\ \mathrm{cov}_{ext,eff}(m_{1},m_{2})&=\mathrm{cov}_{\mathrm{ext}}(m_{1},m_{2})+2p(1-p)\mathrm{cov}_{\mathrm{int}}(m_{1},m_{2}). \end{aligned}$$

Mixing decreases the magnitude of the apparent intrinsic covariance, reaching 0 when the data are fully shuffled ($p=1/2$). Mixing increases the extrinsic covariance when the intrinsic covariance is positive, as is the case for most conditions in our dataset.

While we cannot directly estimate the extent of mixing in our data, by artificially shuffling the left-right label on measured transcripts (**Fig. S9**), we demonstrate that the data are far from fully mixed. In particular, we see considerable increases, often by 50-100% in extrinsic noise due to mixing (**Fig. S9A**). Similarly, mixing suppresses the observed intrinsic covariances (**Fig. S9B**). Both results are consistent with the analysis in this section.

### Four-state gene expression model with extrinsic noise

Having determined the magnitude of extrinsic noise (in terms of the variance and covariance of the production rate $\mu$) in long cells, we can now incorporate the estimated extrinsic noise into our joint distribution fits of the number of transcripts in short cells (< 1.7 μm).

For the sake of simplicity, we assume that $\mu$ is log-normal distributed: $\ln\mu\mathcal{\sim N(}\nu,\sigma)$ with mean $\langle\mu\rangle=e^{\nu+\sigma^{2}/2}$ and $var(\mu)=e^{2\nu+\sigma^{2}}(e^{\sigma^{2}}-1)$. Thus, the parameter $\sigma$ can be directly determined from the coefficient of variation:

$$\begin{matrix} \sigma=\sqrt{ln(1+\mathrm{CV}_{\mu}^{2})} \end{matrix}$$

with $\mathrm{CV}_{\mu}$ determined from long cells.

To compute the transcript distribution $P(m)$ in the model, we approximate the log-normal distribution with a three-point stencil at $\mu=(e^{\nu-\delta},e^{\nu},e^{\nu+\delta})$, with probabilities $(p,1-2p,p)$, respectively. The distribution of transcript number $P(m)$ is given by:

$$\begin{matrix} P(m)=pP(m|\mu=e^{\nu-\delta})+(1-2p)P(m|\mu=e^{\nu})+pP(m|\mu=e^{\nu+\delta}), \end{matrix}$$

with $\nu=\ln\left\langle\mu\right\rangle-\frac{\sigma^{2}}{2}$. Parameters $\delta$ and $p$ are determined by matching the mean and variance of $\mu$:

$$\begin{matrix} p=\frac{1}{2e^{\frac{\sigma^{2}}{2}}+e^{\sigma^{2}}+3}, \delta=2\mathrm{arccosh}\left( \frac{1}{2}\sqrt{\frac{e^{2\sigma^{2}}-1}{e^{\frac{\sigma^{2}}{2}}-1}} \right), \end{matrix}$$

which only depends on $\sigma$ and therefore $\mathrm{CV}_{\mu}$.

For two genes with correlated extrinsic noise, we use a nine-point stencil in the space spanned by $\mu_{1}=(e^{\nu_{1}-\delta_{1}},e^{\nu_{1}},e^{\nu_{1}+\delta_{1}})$ and $\mu_{2}=(e^{\nu_{2}-\delta_{2}},e^{\nu_{2}},e^{\nu_{2}+\delta_{2}})$, with $\nu_{1,2}$ and $\delta_{1,2}$ determined by the single-gene three-point stencil as constructed above. The joint distribution $P(m_{1},m_{2})$ is thus given by a linear combination of those of 9 models with different production rates $(\mu_{1})_{i}$ and $(\mu_{2})_{j}$,

$$\begin{matrix} P(m_{1},m_{2})=\sum_{i=1}^{3} \sum_{j=1}^{3} w_{i,j}P(m_{1},m_{2}|\mu_{1}=(\mu_{1})_{i},\mu_{2}=(\mu_{2})_{j}), \end{matrix}$$

where the weights $w_{i,j}$ follow marginal distributions as defined by the single-gene three-point stencils. Thus, with a suitable stencil $w_{i,j}$, we can account for correlated extrinsic noise in our fits by computing the negative log likelihood loss, Eq. (3), using $P(m_{1},m_{2})$ defined above, with parameters and weights determined by the extrinsic noise.

We choose the stencil $w$ to be a linear combination of the independent stencil $w_{\mathrm{ind}}$, the maximally correlated stencil $w_{\mathrm{corr}}$, and the maximally anticorrelated stencil $w_{\mathrm{anti}}$:

$$\begin{matrix} w=\lambda_{+}w_{\mathrm{corr}}+\lambda_{-}w_{\mathrm{anti}}+(1-\lambda_{+}-\lambda_{-})w_{\mathrm{ind}}, \end{matrix}$$

where

$$\begin{matrix} w_{\mathrm{ind}}=\left( \begin{matrix} p_{1}p_{2} & p_{1}(1-2p_{2}) & p_{1}p_{2} \\ (1-2p_{1})p_{2} & (1-2p_{1})(1-2p_{2}) & (1-2p_{1})p_{2} \\ p_{1}p_{2} & p_{1}(1-2p_{2}) & p_{1}p_{2} \end{matrix} \right), \end{matrix}$$

and

$$\begin{matrix} w_{\mathrm{corr}}=\left\{ \begin{matrix} \left( \begin{matrix} p_{1} & 0 & 0 \\ p_{2}-p_{1} & 1-2p_{2} & p_{2}-p_{1} \\ 0 & 0 & p_{1} \end{matrix} \right) & p_{1}<p_{2} \\ \left( \begin{matrix} p_{2} & p_{1}-p_{2} & 0 \\ 0 & 1-2p_{1} & 0 \\ 0 & p_{1}-p_{2} & p_{2} \end{matrix} \right) & p_{1}\geq p_{2} \end{matrix} \right. , \end{matrix}$$

$$w_{\mathrm{anti}}=\left\{ \begin{matrix} \left( \begin{matrix} 0 & 0 & p_{1} \\ p_{2}-p_{1} & 1-2p_{2} & p_{2}-p_{1} \\ p_{1} & 0 & 0 \end{matrix} \right) & p_{1}<p_{2} \\ \left( \begin{matrix} 0 & p_{1}-p_{2} & p_{2} \\ 0 & 1-2p_{1} & 0 \\ p_{2} & p_{1}-p_{2} & 0 \end{matrix} \right) & p_{1}\geq p_{2} \end{matrix} \right.$$

where $p_{1}$ and $p_{2}$ are the probabilities for the three-point stencil for individual genes. These stencils are designed such that the marginals always match the mean and variance of individual genes, and the linear combination independently adjusts the covariance:

$$\begin{matrix} \mathrm{cov}_{w}\left( \mu_{1},\mu_{2} \right)=\lambda_{+}\mathrm{cov}_{\mathrm{corr}}\left( \mu_{1},\mu_{2} \right)+\lambda_{-}\mathrm{cov}_{\mathrm{anti}}\left( \mu_{1},\mu_{2} \right) \end{matrix}$$

Since the observed extrinsic covariance is mostly positive, we can set $\lambda_{-}=0$, and adjust $\lambda_{+}\in(0,1)$ to match the normalized covariance ($\frac{\mathrm{cov}_{\mathrm{ext}}(m_{1},m_{2})}{\overline{\langle m_{i}\rangle}\cdot\overline{\langle m_{j}\rangle}})$ of extrinsic noise, as determined in the previous section.

In the model, we fitted the model considering extrinsic noise at half of the estimated strength since the extrinsic noise estimate from the last section is likely an overestimate. However, fitting the model with extrinsic noise at maximum strength (**Fig. S12D**) and without extrinsic noise (**Fig. S12D**), we find that the model still successfully captured the measured gene expression statistics as well as the non-monotonic relation between gene-gene interaction strength and overall supercoiling level.

| **Table S1. Chromosomally integrated DNA construct dimensions** | |
| --- | --- |
| **Codirectional (*codir*) Construct Dimensions** |  |
| *3xO_L_123* to Gene 1 Promoter start | 229 bp |
| Gene 1 Promoter start to TSS | 38 bp |
| Gene 1 TSS to start of Gene 1 Terminator (*rrnB* T1 + *rrnD*) | 3429 bp |
| Start of Gene 1 Terminator to end of Terminator (*rrnB* T1 + *rrnD*) | 276 bp |
| End of Gene 1 Terminator to start of Gene 2 Promoter | 30 bp |
| Gene 2 Promoter start to TSS | 38 bp |
| Gene 2 Promoter to start of Terminator (*rrnD*) | 1684 bp |
| Start of Gene 2 Terminator to end of Terminator (*rrnD*) | 55 bp |
| End of Gene 2 Terminator to 2x*O_R_123* | 303 bp |
| Length of 2x*O_R_123* | 158 bp |
| *2xO_R_123* to Gene 3 Promoter | 595 bp |
| Gene 3 Promoter to end of Construct | 934 bp |
| End of Construct to next chromosomal Terminator | 3292 bp |
| **Divergent (Div) Construct Dimensions** |  |
| *3xO_L_123* to Gene 1 Terminator end (*rrnB* T1 + *rrnD*) | 185 bp |
| Start of Gene 1 Terminator to end of terminator (*rrnB* T1 + *rrnD*) | 276 bp |
| Start of Gene 1 Terminator to Gene 1 TSS | 3429 bp |
| Gene 1 Promoter start to TSS | 38 bp |
| Start of Gene 1 Promoter to start of Gene 2 Promoter | 74 bp |
| Gene 2 Promoter start to TSS | 38 bp |
| Gene 2 Promoter to start of Terminator (*rrnD*) | 1684 bp |
| Start of Gene 2 Terminator to end of Terminator (*rrnD*) | 55 bp |
| End of Gene 2 Terminator to 2x*O_R_123* | 303 bp |
| Length of 2x*O_R_123* | 158 bp |
| 2x*O_R_123* to Gene 3 Promoter | 595 bp |
| Gene 3 Promoter to end of Construct | 934 bp |
| End of Construct to next chromosomal Terminator | 3292 bp |

| **Table S2. E coli strains used for smFISH experiments.** | | |  |
| --- | --- | --- | --- |
| **Strain Name** | **Background** | **Description** | **Source** |
| XF001 | MG1655 | Landing pad strain: *lacI* replaced by the LP1-*tet*^R^-LP2 fragment using λ RED recombination | ^4^ |
| NY45 | XF001 | Landing pad insertion of the codirectional construct | This study |
| NY50 | NY45 | Codirectional construct strain expressing λ repressor CI from plasmid pAL18_CI | This study |
| NY51 | NY45 | Codirectional construct strain with control plasmid pAL18_no_λCI | This study |
| NY89 | XF001 | Landing Pad insertion of the divergent construct | This study |
| NY91 | NY89 | Divergent construct strain expressing λ repressor CI from plasmid pAL18_CI | This study |
| NY92 | NY89 | Divergent construct strain with control plasmid pAL18_no_λCI | This study |

| **Table S3: smFISH Probe sequences (5’ to 3’)** | | |
| --- | --- | --- |
| Gene 1 - Fluorescein (FAM) | Gene 2 - Cal Fluor Red 590 | Gene 3 - Quasar 670 |
| gattaagttgggtaacgcca | ggccttttttgatgtttttt | tcttgttcaatggccgatc |
| aaagggggatgtgctgcaag | aacgtttcattgctttgtgc | ggagaacctgcgtgcaatc |
| aactgttgggaagggcgatc | gatgtgtgcgtcggtgaatg | aatagcctctccacccaag |
| tgaggggacgacgacagtat | ttctgcgtaggtgatgttta | ctgttgtgcccagtcatag |
| tgtagatgggcgcatcgtaa | cagacgtacggacatttcga | gcatcagagcagccgattg |
| gtaatgggataggtcacgtt | cgtaacgtttcattgcttct | cgctgacagccggaacacg |
| gggaacaaacggcggattga | atacggtggttggtgttcag | acaaaaagaaccgggcgcc |
| atgtgagcgagtaacaaccc | ggagttttcggagcatacta | tgcagttcattcagggcac |
| tagccagctttcatcaacat | cagtaccggcatgaagaact | cacagctgcgcaaggaacg |
| aaaataattcgcgtctggcc | tactgctacgccgatgaaca | gcttcagtgacaacgtcga |
| ttgcaccacagatgaaacgc | gttcgttgtagatgtcgttt | gcacttcgcccaatagcag |
| ttcagacggcaaacgactgt | atgttcatggagttcagcag | agatgacaggagatcctgc |
| cgcgtaaaaatgcgctcagg | atacgaatactacggtcggc | ctttctcggcaggagcaag |
| tcctgatcttccagataact | tacgttcaggattttctgca | attgcatcagccatgatgg |
| gagacgtcacggaaaatgcc | tggatgatcggcagtttttt | cggatcaagcgtatgcagc |
| tgtgtagtcggtttatgcag | gttttggagtccatgatgat | tggtcgaatgggcaggtag |
| ggcaacatggaaatcgctga | cgaaggtgtacatggactgg | cgatgcgatgtttcgcttg |
| cgcggctgaaatcatcatta | aagtcgtattcgttgaagcc | cttccatccgagtacgtgc |
| cacatctgaacttcagcctc | ttttgtcacggtcgaaggat | atcctgatcgacaagaccg |
| gtaggtagtcacgcaactcg | gagttcatgatcagtgcgat | tgatgctcttcgtccagat |
| caccctgccataaagaaact | agaaacgtacgcatgcggta | ttgagcctggcgaacagtt |
| ctcatcgataatttcaccgc | atgatctggttgccgaagat | gggtcacgacgagatcatc |
| agacgtagtgtgacgcgatc | cggtactacggacaggattg | tgatattcggcaagcaggc |
| cacagtttcgggttttcgac | ggtacatcagtactacacgg | aaaagcggccattttccac |
| gcacgatagagattcgggat | aacgcaggaacagttcttct | cggccacagtcgatgaatc |
| caggcttctgcttcaatcag | gactggattttgtagtcctg | ccaacgctatgtcctgata |
| agcagcagaccattttcaat | aaggagaacagggtcggtac | gccaagctcttcagcaata |
| taacgcctcgaatcagcaac | tatttgtcgatcagggtgga | aagcacgaggaagcggtca |
| tgaccatgcagaggatgatg | atttcgtgcaggttggacag | gaatcgggagcggcgatac |
| ttcatcagcaggatatcctg | tacttctttggacagcggtg | gatagaaggcgatgcgctg |
| cacggcgttaaagttgttct | ggaaacgttttgctactgct |  |
| agcggatggttcggataatg | tgatcaggattgcggaggtg |  |
| gggtttcaatattggcttca | aacggtactactttgcctac |  |
| gatcatcggtcagacgattc | gtctactacttttgcttcga |  |
| gatcacactcgggtgattac | tttacgtagccggacatgat |  |
| atacagcgcgtcgtgattag | ctttgtcgatcagtgcgttg |  |
| ggatcgacagatttgatcca | gaagaagtgttcgtcttcgt |  |
| cgcgtacatcgggcaaataa | agggatttcagacggtctac |  |
| aagccattttttgatggacc | ctggtagcctttgtatttga |  |
| tattcgcaaaggatcagcgg | caggatggattccagttctg |  |
| aaaccgccaagactgttacc | tcgaagatgttcgggtgctg |  |
| aaacgcctgccagtatttag | gtgttccagtactactactg |  |
| cctgtaaacggggatactga | ctttttcggtcatggttttg |  |
| gttgccgttttcatcatatt | gatgctacgtagtctacgat |  |
| ttcggcgtatcgccaaaatc | agtttttttgcggtggttac |  |
| cgttcatacagaactggcga | ttcggtacttcgtctacgaa |  |
| aaactgctgctggtgttttg | tttcacggattttacgtgcg |  |
| tttgcccggataaacggaac | cctttttttgctttgatcag |  |

| **Table S4. smFISH mean RNA copy number measurements.** | | | | |
| --- | --- | --- | --- | --- |
| **Condition** | **Number of Cells (N)** | **RNA1**  ***(Mean ± SEM)*** | **RNA2**  ***(Mean ± SEM)*** | **RNA3**  ***(Mean ± SEM)*** |
| Codirectional | 2791 | 3.58 ± 0.06 | 2.55 ± 0.06 | 5.64 ± 0.10 |
| Codirectional + nov | 2673 | 3.71 ± 0.07 | 2.34 ± 0.06 | 1.03 ± 0.04 |
| Codirectional + scn | 1081 | 3.15 ± 0.10 | 1.80 ± 0.07 | 3.28 ± 0.13 |
| Codirectional + loop | 2523 | 1.73 ± 0.04 | 1.71 ± 0.05 | 3.03 ± 0.10 |
| Codirectional + loop + nov | 1999 | 1.52 ± 0.04 | 1.16 ± 0.06 | 0.42 ± 0.04 |
| Codirectional + loop + scn | 618 | 2.43 ± 0.10 | 2.04 ± 0.10 | 3.72 ± 0.19 |
| Divergent | 2743 | 2.73 ± 0.06 | 1.25 ± 0.04 | 3.53 ± 0.08 |
| Divergent + nov | 3055 | 2.68 ± 0.06 | 1.15 ± 0.03 | 1.19 ± 0.04 |
| Divergent + scn | 1046 | 3.53 ± 0.12 | 2.43 ± 0.09 | 3.97 ± 0.16 |
| Divergent + loop | 2051 | 1.18 ± 0.04 | 1.01 ± 0.05 | 1.34 ± 0.04 |
| Divergent + loop + nov | 2301 | 1.28 ± 0.04 | 0.75 ± 0.03 | 0.54 ± 0.02 |
| Divergent + loop + scn | 605 | 2.51 ± 0.14 | 1.66 ± 0.10 | 3.07 ± 0.11 |

**Table S5: KS2 test *p* values for pairwise comparisons of all conditions (see attached Excel file)**

| **Table S6. Three-way ANOVA analysis statistics. The most significant factor (largest F value) for each RNA species is bolded.** | | | | | | |
| --- | --- | --- | --- | --- | --- | --- |
| **Condition** | **RNA** | **SumSq** | **DF** | **MeanSq** | **F** | **pValue** |
| Gene Orientation | RNA 1 | 1889 | 1 | 1889 | 237 | 4E-53 |
| **Looping State** | **RNA 1** | **14991** | **1** | **14991** | **1877** | **0E+00** |
| Drug | RNA 1 | 684 | 2 | 342 | 43 | 3E-19 |
| Gene Orientation: Looping State | RNA 1 | 271 | 1 | 271 | 34 | 6E-09 |
| Gene Orientation: Drug | RNA 1 | 738 | 2 | 369 | 46 | 9E-21 |
| Looping State: Drug | RNA 1 | 530 | 2 | 265 | 33 | 4E-15 |
| Gene Orientation: Looping State: Drug | RNA 1 | 193 | 2 | 97 | 12 | 6E-06 |
| **Gene Orientation** | **RNA 2** | **3509** | **1** | **3509** | **573** | **4E-125** |
| Looping State | RNA 2 | 2072 | 1 | 2072 | 338 | 5E-75 |
| Drug | RNA 2 | 900 | 2 | 450 | 73 | 2E-32 |
| Gene Orientation: Looping State | RNA 2 | 300 | 1 | 300 | 49 | 3E-12 |
| Gene Orientation: Drug | RNA 2 | 1134 | 2 | 567 | 93 | 9E-41 |
| Looping State: Drug | RNA 2 | 184 | 2 | 92 | 15 | 3E-07 |
| Gene Orientation: Looping State: Drug | RNA 2 | 494 | 2 | 247 | 40 | 3E-18 |
| Gene Orientation | RNA 3 | 3172 | 1 | 3172 | 243 | 2E-54 |
| Looping State | RNA 3 | 10562 | 1 | 10562 | 809 | 6E-175 |
| **Drug** | **RNA 3** | **40749** | **2** | **20375** | **1560** | **0E+00** |
| Gene Orientation: Looping State | RNA 3 | 1 | 1 | 1 | 0 | 8E-01 |
| Gene Orientation: Drug | RNA 3 | 6155 | 2 | 3077 | 236 | 5E-102 |
| Looping State: Drug | RNA 3 | 5094 | 2 | 2547 | 195 | 9E-85 |
| Gene Orientation: Looping State: Drug | RNA 3 | 449 | 2 | 225 | 17 | 3E-08 |

| **Table S7. smFISH measurements of mean RNA puncta per cell and RNA copy number per punctum across all genes and conditions.** | | | | | | | | |
| --- | --- | --- | --- | --- | --- | --- | --- | --- |
|  | **RNA1** | |  | **RNA2** | |  | **RNA3** | |
| **Condition** | **Number of Puncta/Cell (Mean ± SEM)** | **Number of RNA/Punctum (Mean ± SEM)** |  | **Number of Puncta/Cell (Mean ± SEM)** | **Number of RNA/Punctum (Mean ± SEM)** |  | **Number of Puncta/Cell (Mean ± SEM)** | **Number of RNA/Punctum (Mean ± SEM)** |
| codir | 1.31 ± 0.02 | 2.74 ± 0.03 |  | 1.12 ± 0.02 | 2.28 ± 0.04 |  | 1.39 ± 0.01 | 4.07 ± 0.05 |
| codir-loop | 0.86 ± 0.02 | 2.02 ± 0.03 |  | 0.82 ± 0.02 | 2.10 ± 0.05 |  | 0.84 ± 0.02 | 3.62 ± 0.08 |
| codir-loop-nov | 0.75 ± 0.02 | 2.02 ± 0.03 |  | 0.52 ± 0.02 | 2.25 ± 0.07 |  | 0.15 ± 0.01 | 2.78 ± 0.22 |
| codir-loop-scn | 1.13 ± 0.03 | 2.16 ± 0.06 |  | 1.23 ± 0.05 | 1.66 ± 0.04 |  | 1.15 ± 0.04 | 3.24 ± 0.12 |
| codir-nov | 1.33 ± 0.02 | 2.79 ± 0.04 |  | 0.91 ± 0.02 | 2.57 ± 0.04 |  | 0.56 ± 0.02 | 1.82 ± 0.04 |
| codir-scn | 1.22 ± 0.03 | 2.57 ± 0.05 |  | 1.01 ± 0.03 | 1.79 ± 0.04 |  | 1.22 ± 0.03 | 2.69 ± 0.07 |
| div | 1.19 ± 0.02 | 2.30 ± 0.03 |  | 0.63 ± 0.02 | 1.96 ± 0.03 |  | 1.25 ± 0.02 | 2.84 ± 0.05 |
| div-loop | 0.73 ± 0.02 | 1.62 ± 0.03 |  | 0.53 ± 0.02 | 1.92 ± 0.05 |  | 0.80 ± 0.02 | 1.67 ± 0.03 |
| div-loop-nov | 0.76 ± 0.02 | 1.69 ± 0.03 |  | 0.40 ± 0.01 | 1.89 ± 0.04 |  | 0.39 ± 0.01 | 1.38 ± 0.03 |
| div-loop-scn | 1.07 ± 0.04 | 2.35 ± 0.09 |  | 0.94 ± 0.04 | 1.77 ± 0.06 |  | 1.40 ± 0.04 | 2.19 ± 0.05 |
| div-nov | 1.08 ± 0.02 | 2.48 ± 0.03 |  | 0.61 ± 0.01 | 1.88 ± 0.03 |  | 0.60 ± 0.01 | 2.00 ± 0.04 |
| div-scn | 1.18 ± 0.03 | 2.98 ± 0.07 |  | 1.25 ± 0.04 | 1.94 ± 0.05 |  | 1.19 ± 0.03 | 3.34 ± 0.09 |

**Table S8A: Intrinsic-extrinsic noise decomposition of the total noise (**${\boldsymbol{\sigma}^{\boldsymbol{2}}}/{\boldsymbol{\mu}^{\boldsymbol{2}}}$**) for each gene in long cells (>2000 nm). Reported values are noise** $\boldsymbol{\pm}$ **bootstrapped errors.**

| **Condition** | ***G1* Intrinsic** | ***G1* Extrinsic** | ***G2* Intrinsic** | ***G2* Extrinsic** | ***G3* Intrinsic** | ***G3* Extrinsic** |
| --- | --- | --- | --- | --- | --- | --- |
| *codir*-scn | 0.80 ± 0.08 | 1.61 ± 0.16 | 0.93 ± 0.23 | 0.46 ± 0.07 | 0.60 ± 0.13 | 1.05 ± 0.12 |
| *codir* | 0.87 ± 0.05 | 1.62 ± 0.14 | 0.60 ± 0.05 | 0.24 ± 0.05 | 0.36 ± 0.07 | 0.44 ± 0.04 |
| *codir*-nov | 0.94 ± 0.07 | 2.00 ± 0.16 | 1.75 ± 0.25 | 0.27 ± 0.04 | 0.51 ± 0.08 | 2.03 ± 0.24 |
| *codir*-loop-scn | 1.49 ± 0.19 | 1.43 ± 0.29 | 1.10 ± 0.23 | 0.23 ± 0.08 | 0.73 ± 0.18 | 0.98 ± 0.16 |
| *codir*-loop | 1.67 ± 0.09 | 2.79 ± 0.28 | 2.18 ± 0.23 | 0.40 ± 0.07 | 0.84 ± 0.15 | 1.52 ± 0.14 |
| *codir*-loop-nov | 1.97 ± 0.14 | 3.69 ± 0.47 | 16.59 ± 3.16 | 0.28 ± 0.08 | 1.67 ± 0.40 | 8.78 ± 1.90 |
| *div*-scn | 1.17 ± 0.11 | 1.54 ± 0.21 | 1.27 ± 0.18 | 0.59 ± 0.09 | 0.54 ± 0.09 | 0.98 ± 0.12 |
| *div* | 1.14 ± 0.11 | 2.99 ± 0.25 | 0.76 ± 0.07 | 0.38 ± 0.06 | 0.59 ± 0.11 | 0.83 ± 0.06 |
| *div*-nov | 1.10 ± 0.09 | 3.23 ± 0.24 | 2.30 ± 0.36 | 0.46 ± 0.05 | 0.55 ± 0.13 | 1.54 ± 0.15 |
| *div*-loop-scn | 1.73 ± 0.30 | 1.67 ± 0.25 | 0.86 ± 0.10 | 1.00 ± 0.27 | 1.34 ± 0.29 | 0.39 ± 0.13 |
| *div*-loop | 2.24 ± 0.19 | 5.02 ± 0.51 | 1.87 ± 0.15 | 0.55 ± 0.10 | 1.59 ± 0.33 | 0.99 ± 0.20 |
| *div*-loop-nov | 2.05 ± 0.18 | 5.47 ± 0.48 | 5.10 ± 0.77 | 0.56 ± 0.11 | 1.29 ± 0.39 | 0.94 ± 0.19 |

**Table S8B. Intrinsic-extrinsic Fano factor decomposition of the total Fano factor (**$\boldsymbol{\sigma}^{\boldsymbol{2}}\boldsymbol{/}\boldsymbol{\mu}$**) for each gene in long cells (>2 µm). Reported values are Fano factor** $\boldsymbol{\pm}$ **bootstrapped errors.**

| **Condition** | ***G1* Intrinsic** | ***G1* Extrinsic** | ***G2* Intrinsic** | ***G2* Extrinsic** | ***G3* Intrinsic** | ***G3* Extrinsic** |
| --- | --- | --- | --- | --- | --- | --- |
| *codir*-scn | 1.55 ± 0.15 | 0.89 ± 0.14 | 1.75 ± 0.17 | 0.65 ± 0.14 | 1.78 ± 0.44 | 2.01 ± 0.22 |
| *codir* | 1.93 ± 0.12 | 0.53 ± 0.10 | 2.52 ± 0.21 | 0.55 ± 0.11 | 2.03 ± 0.15 | 1.46 ± 0.12 |
| *codir*-nov | 2.19 ± 0.17 | 0.62 ± 0.10 | 2.95 ± 0.24 | 0.76 ± 0.12 | 1.27 ± 0.18 | 1.47 ± 0.17 |
| *codir*-loop-scn | 1.99 ± 0.25 | 0.30 ± 0.11 | 1.60 ± 0.32 | 0.81 ± 0.20 | 2.33 ± 0.48 | 2.07 ± 0.34 |
| *codir*-loop | 1.72 ± 0.09 | 0.41 ± 0.07 | 2.85 ± 0.29 | 0.85 ± 0.15 | 3.78 ± 0.40 | 2.63 ± 0.25 |
| *codir*-loop-nov | 1.80 ± 0.13 | 0.25 ± 0.07 | 2.81 ± 0.36 | 1.27 ± 0.30 | 4.71 ± 0.90 | 2.49 ± 0.54 |
| *div*-scn | 2.28 ± 0.21 | 1.16 ± 0.17 | 2.10 ± 0.29 | 0.73 ± 0.12 | 2.85 ± 0.39 | 2.20 ± 0.26 |
| *div* | 1.92 ± 0.18 | 0.64 ± 0.10 | 2.25 ± 0.19 | 0.44 ± 0.08 | 1.62 ± 0.15 | 1.76 ± 0.14 |
| *div*-nov | 1.86 ± 0.15 | 0.78 ± 0.08 | 2.16 ± 0.16 | 0.37 ± 0.09 | 1.58 ± 0.25 | 1.06 ± 0.10 |
| *div*-loop-scn | 2.72 ± 0.47 | 1.58 ± 0.42 | 1.61 ± 0.24 | 1.29 ± 0.28 | 1.51 ± 0.18 | 0.68 ± 0.23 |
| *div*-loop | 1.54 ± 0.13 | 0.38 ± 0.07 | 2.96 ± 0.30 | 0.94 ± 0.20 | 1.41 ± 0.12 | 0.75 ± 0.15 |
| *div*-loop-nov | 1.56 ± 0.14 | 0.43 ± 0.08 | 2.42 ± 0.21 | 0.57 ± 0.17 | 1.55 ± 0.23 | 0.29 ± 0.06 |

| **Table S9: Spearman correlation coefficients (rho) and the associated bootstrapped errors (err) of all experimentally measured and computationally scrambled (ctrl) RNA pairs. Bolded values indicate significant correlations.** | | | | | | |
| --- | --- | --- | --- | --- | --- | --- |
| **Condition** | **G1 -G2**  **(rho ± err)** | **G1 -G2_ctrl**  **(rho ± err)** | **G1 -G3**  **(rho ± err)** | **G1 -G3_ctrl**  **(rho ± err)** | **G2 -G3**  **(rho ± err)** | **G2 -G3_ctrl**  **(rho ± err)** |
| codir | **0.19 ± 0.03** | 0.00 ± 0.03 | **0.30 ± 0.03** | 0.00 ± 0.03 | **0.25 ± 0.03** | 0.00 ± 0.03 |
| codir-loop | **0.10 ± 0.04** | 0.00 ± 0.04 | **0.19 ± 0.04** | 0.00 ± 0.04 | **0.31 ± 0.04** | 0.00 ± 0.04 |
| codir-loop-nov | **0.09 ± 0.04** | 0.00 ± 0.04 | **0.05 ± 0.04** | 0.00 ± 0.04 | **0.34 ± 0.05** | 0.00 ± 0.04 |
| codir-loop-scn | **0.23 ± 0.10** | -0.01 ± 0.10 | -0.03 ± 0.10 | 0.00 ± 0.10 | **0.18 ± 0.09** | 0.00 ± 0.10 |
| codir-nov | **0.23 ± 0.03** | 0.00 ± 0.03 | **0.22 ± 0.03** | 0.00 ± 0.03 | **0.30 ± 0.03** | 0.00 ± 0.03 |
| codir-scn | 0.08 ± 0.06 | 0.00 ± 0.06 | **0.53 ± 0.05** | 0.00 ± 0.06 | **0.10 ± 0.06** | 0.00 ± 0.06 |
| div | **0.18 ± 0.04** | 0.00 ± 0.04 | **0.37 ± 0.03** | 0.00 ± 0.04 | **0.11 ± 0.04** | 0.00 ± 0.04 |
| div-loop | **0.24 ± 0.05** | 0.00 ± 0.05 | **0.34 ± 0.05** | 0.00 ± 0.05 | **0.16 ± 0.05** | 0.00 ± 0.05 |
| div-loop-nov | **0.21 ± 0.05** | 0.00 ± 0.05 | **0.33 ± 0.05** | 0.00 ± 0.05 | **0.10 ± 0.05** | 0.00 ± 0.05 |
| div-loop-scn | -0.04 ± 0.09 | 0.00 ± 0.09 | **0.27 ± 0.08** | 0.00 ± 0.09 | **-0.12 ± 0.09** | -0.01 ± 0.09 |
| div-nov | **0.27 ± 0.03** | 0.00 ± 0.03 | 0.03 ± 0.04 | 0.00 ± 0.03 | 0.01 ± 0.03 | 0.00 ± 0.03 |
| div-scn | **0.10 ± 0.06** | 0.00 ± 0.07 | **0.47 ± 0.05** | 0.00 ± 0.07 | **0.31 ± 0.06** | 0.00 ± 0.07 |

**Table S10: Joint distribution fitting parameters and fit coupling constants** $\boldsymbol{J}_{\boldsymbol{S}}$ **for all conditions.** Parameters reported are from the $\Gamma=0$ ($\alpha_{1}=\alpha_{2}$) fits corresponding to the results in **Fig. 5**. Note that fitting steady-state data only constrains ratios of parameters. As described in model reduction section of STAR Methods, we fixed $d_{1}=d_{2}=1$ , measuring time in units of the average mRNA lifetime. Thus, the kinetic rate parameters $\mu$ and $k_{on}$ are measured in units of 1/mRNA lifetime. The parameters $\alpha$ and $J_{S}$ are dimensionless. We also fixed $k_{\text{off}}=10$ in fits; results are insensitive to $k_{\text{off}}$ in the bursting regime.

| **(*G1, G2*)** | | | | | | |
| --- | --- | --- | --- | --- | --- | --- |
| **Condition** | $\boldsymbol{\mu}_{\boldsymbol{1}}$ | $\boldsymbol{\mu}_{\boldsymbol{2}}$ | $\boldsymbol{k}_{\text{on}\boldsymbol{,1}}$ | $\boldsymbol{k}_{\text{on}\boldsymbol{,2}}$ | $\boldsymbol{\alpha}_{\boldsymbol{1}}\boldsymbol{,}\boldsymbol{\alpha}_{\boldsymbol{2}}$ | $\boldsymbol{J}_{\boldsymbol{S}}\boldsymbol{=}\boldsymbol{J}_{\boldsymbol{12}}\boldsymbol{+}\boldsymbol{J}_{\boldsymbol{21}}$ |
| *codir*-scn | 25.63 | 16.16 | 0.81 | 0.75 | 1.76 | 0.10 |
| *codir* | 18.81 | 25.05 | 1.35 | 0.67 | 3.32 | 0.33 |
| *codir*-nov | 21.12 | 32.47 | 1.28 | 0.37 | 4.25 | 0.39 |
| *codir*-loop-scn | 7.39 | 11.86 | 2.46 | 1.03 | 4.31 | 0.60 |
| *codir*-loop | 13.07 | 24.42 | 0.94 | 0.45 | 2.98 | 0.22 |
| *codir*-loop-nov | 20.71 | 41.31 | 0.60 | 0.21 | 2.45 | 0.10 |
| *div*-scn | 27.10 | 38.40 | 1.13 | 0.44 | 2.24 | 0.16 |
| *div* | 15.96 | 17.13 | 1.29 | 0.41 | 3.70 | 0.34 |
| *div*-nov | 20.86 | 18.86 | 0.77 | 0.27 | 10.09 | 0.65 |
| *div*-loop-scn | 7.39 | 2.04 | 2.61 | 12.01 | 1.00 | 0.00 |
| *div*-loop | 9.78 | 35.85 | 0.79 | 0.15 | 11.86 | 0.76 |
| *div*-loop-nov | 10.99 | 16.22 | 0.63 | 0.17 | 12.56 | 0.68 |
| **(*G1, G3*)** | | | | | | |
| **Condition** | $\boldsymbol{\mu}_{\boldsymbol{1}}$ | $\boldsymbol{\mu}_{\boldsymbol{2}}$ | $\boldsymbol{k}_{\boldsymbol{on,1}}$ | $\boldsymbol{k}_{\boldsymbol{on,2}}$ | $\boldsymbol{\alpha}_{\boldsymbol{1}}\boldsymbol{,}\boldsymbol{\alpha}_{\boldsymbol{2}}$ | $\boldsymbol{J}_{\boldsymbol{S}}\boldsymbol{=}\boldsymbol{J}_{\boldsymbol{12}}\boldsymbol{+}\boldsymbol{J}_{\boldsymbol{21}}$ |
| *codir*-scn | 19.36 | 26.67 | 0.62 | 0.39 | 23.17 | 1.02 |
| *codir* | 17.73 | 38.33 | 1.18 | 0.8 | 6.32 | 0.6 |
| *codir*-nov | 20.18 | 15.05 | 1.29 | 0.23 | 8.35 | 0.79 |
| *codir*-loop-scn | 7.81 | 44.15 | 2.99 | 0.65 | 1 | 0 |
| *codir*-loop | 13.2 | 90.93 | 0.91 | 0.24 | 5.75 | 0.42 |
| *codir*-loop-nov | 20.56 | 38.98 | 0.61 | 0.06 | 3.89 | 0.18 |
| *div*-scn | 24.4 | 45.95 | 0.92 | 0.39 | 13.78 | 0.91 |
| *div* | 13.73 | 20.76 | 1.11 | 0.81 | 7.69 | 0.68 |
| *div*-nov | 22.3 | 28.51 | 0.87 | 0.38 | 1.42 | 0.05 |
| *div*-loop-scn | 8.87 | 11.82 | 1.66 | 1.95 | 2.8 | 0.37 |
| *div*-loop | 7.7 | 8.34 | 0.51 | 0.74 | 20 | 1.02 |
| *div*-loop-nov | 9.35 | 8.41 | 0.6 | 0.15 | 36.52 | 1.47 |
| **(*G2, G3*)** | | | | | | |
| **Condition** | $\boldsymbol{\mu}_{\boldsymbol{1}}$ | $\boldsymbol{\mu}_{\boldsymbol{2}}$ | $\boldsymbol{k}_{\boldsymbol{on,1}}$ | $\boldsymbol{k}_{\boldsymbol{on,2}}$ | $\boldsymbol{\alpha}_{\boldsymbol{1}}\boldsymbol{,}\boldsymbol{\alpha}_{\boldsymbol{2}}$ | $\boldsymbol{J}_{\boldsymbol{S}}\boldsymbol{=}\boldsymbol{J}_{\boldsymbol{12}}\boldsymbol{+}\boldsymbol{J}_{\boldsymbol{21}}$ |
| *codir*-scn | 15.45 | 29.77 | 0.76 | 0.71 | 2.37 | 0.16 |
| *codir* | 25.31 | 39.9 | 0.63 | 0.97 | 4.84 | 0.42 |
| *codir*-nov | 30.59 | 14.7 | 0.39 | 0.28 | 14.64 | 0.6 |
| *codir*-loop-scn | 12.4 | 40.76 | 1.47 | 0.56 | 2.92 | 0.28 |
| *codir*-loop | 23.55 | 89.2 | 0.44 | 0.23 | 12.79 | 0.55 |
| *codir*-loop-nov | 30.97 | 24.21 | 0.22 | 0.04 | 66.75 | 1.24 |
| *div*-scn | 35.94 | 50.53 | 0.38 | 0.58 | 8.37 | 0.49 |
| *div* | 17.17 | 22.62 | 0.47 | 1.17 | 2.24 | 0.16 |
| *div*-nov | 20.1 | 28.63 | 0.42 | 0.4 | 1 | 0 |
| *div*-loop-scn | 1.12 | 10.86 | 2870.39 | 2.68 | 1 | 0 |
| *div*-loop | 36.82 | 11.58 | 0.18 | 0.92 | 6.22 | 0.45 |
| *div*-loop-nov | 16.78 | 9.99 | 0.25 | 0.34 | 4.98 | 0.2 |

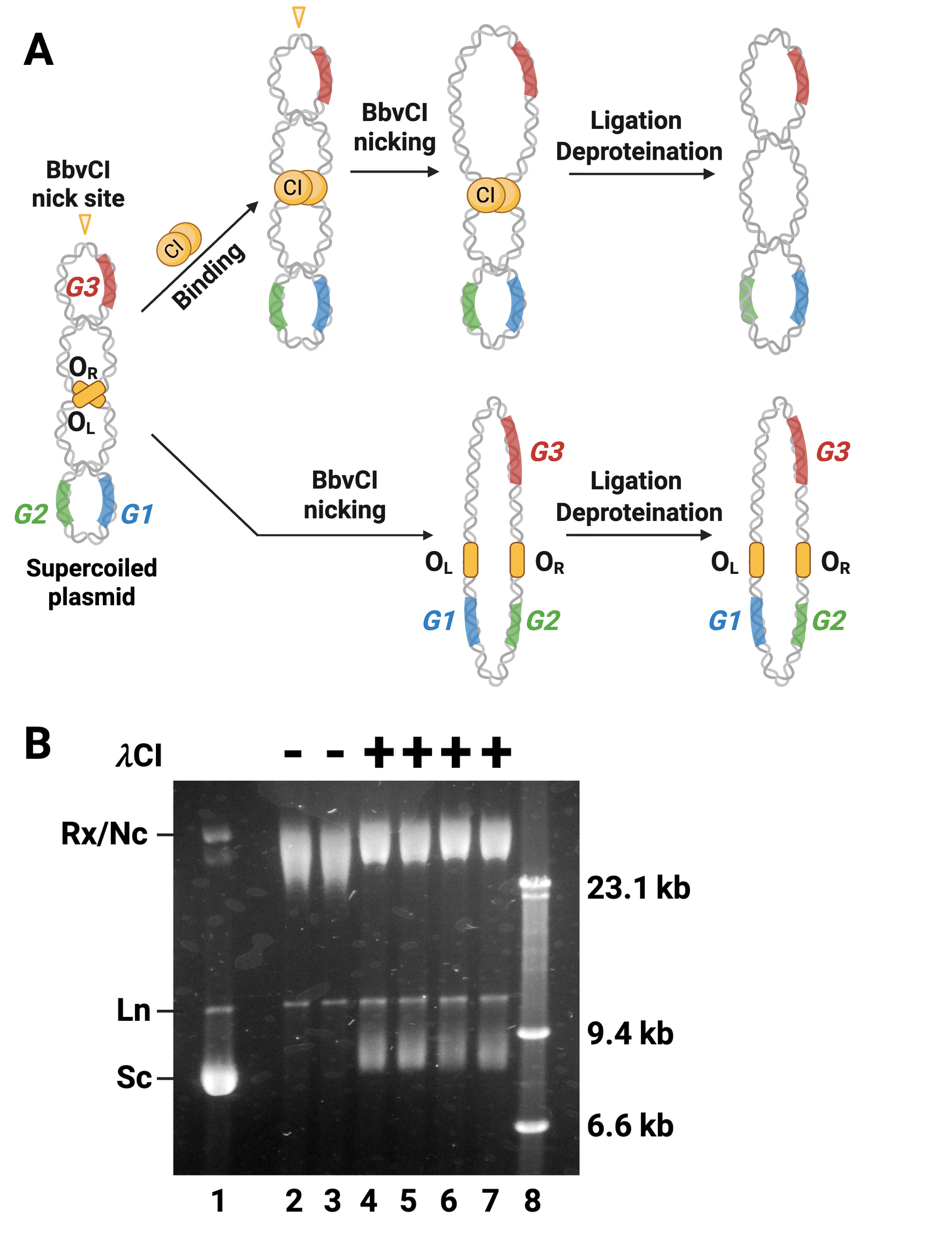
**Figure S1. The binding of λ repressor CI divided supercoiled plasmids into two independent topological domains for plasmid pNY1**.

**(A)** Schematics of the plasmid nicking topology assay. Plasmid pNY1 (~11 Kbp) contains three genes (G1–G3), with the λ repressor operator sites (3×O_L_123 and 2×O_R_123) flanking G1 and G2, and a unique nicking site (Nt.BbvCI) between G1/G2 and G3. The plasmid is naturally negatively supercoiled when purified form cell cultures. In the nicking assay, the plasmid prep is divided into two batches. One batch (top) is incubated with purified λ CI protein, followed by incubation with the nicking enzyme Nt.BbvCI. The binding and octamerization of CI on the operators restricts the relaxation caused by nicking to only the nicking site-containing domain but not the other domain. After ligation and deproteination, the plasmid DNA still contained some supercoils but had a lower supercoiling density compared to that in the beginning. The second batch (bottom) is incubated directly with the nicking enzyme Nt.BbvCI, which relaxes the entire plasmid and hence the plasmid should have minimal supercoils after ligation and deproteination. **(B)** An representative agarose gel showing results of the DNA-nicking topology assay performed as described in **A** (Materials and Methods). Lanes 1-7 contain 0.156 nM of pNY1 plasmid each. Lane 1 is purified plasmid pNY1, which showed typical supercoiled (Sc), linear (Ln), and relaxed/nicked (Rx/Nc) conformers. Lanes 2 and 3 are replicates of pNY1 treated with Nt.BbvCI (20 units) in the absence of λ cI. They contained mainly relaxed or nicked conformers. Lanes 4-7 are replicates of pNY1 first inculcated with 170 nM λ cI and then treated with Nt.BbvCI (20 units). The presence of reduced but still supercoiled plasmids in these lanes (bottom bands), compared to the absence of supercoiled plasmids in Lanes 2-3, confirms that λ CI constrained the topological domain and prevented the propagation of relaxation form one domain to another domain. Lane 8 is λ DNA digested with Hind III as a molecular size marker.

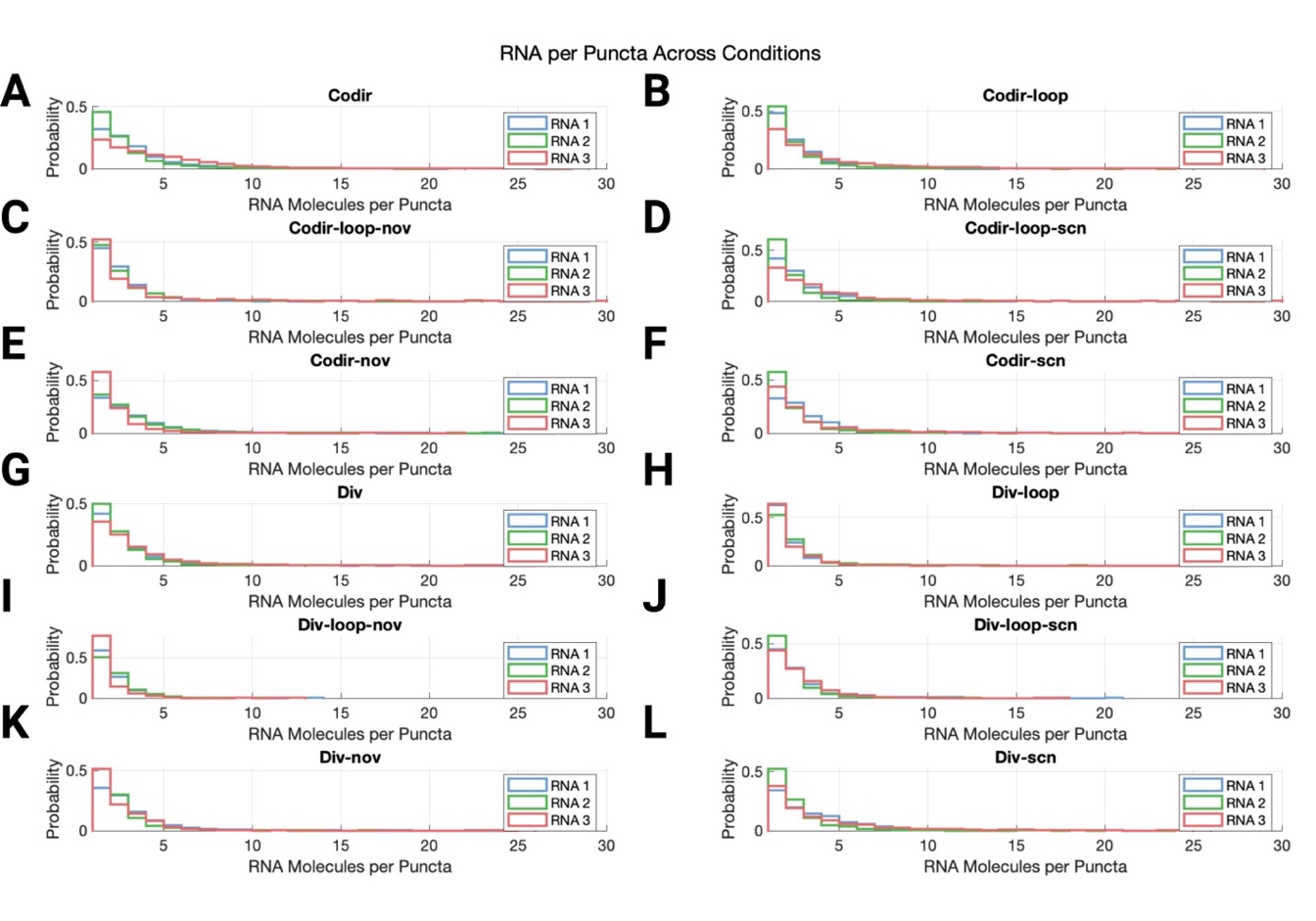

**Figure S2. Distributions of RNA molecules per punctum across genes and conditions**.

Probability distributions of RNA copy number per punctum for RNA1 (blue), RNA2 (green), and RNA3 (red) across all twelve experimental conditions are shown from **A** to **L.** Specifically: (**A**) Codirectional, unlooped, no drug; (**B**) Codirectional, looped, no drug; (**C**) Codirectional, looped, novobiocin-treated; (**D**) Codirectional, looped, seconeolitsine-treated; (**E**) Codirectional, unlooped, novobiocin-treated; (**F**) Codirectional, unlooped, seconeolitsine-treated; (**G**) Divergent, unlooped, no drug; (**H**) Divergent, looped, no drug; (**I**) Divergent, looped, novobiocin-treated; (**J**) Divergent, looped, seconeolitsine-treated; (**K**) Divergent, unlooped, novobiocin-treated; (**L**) Divergent, unlooped, seconeolitsine-treated.

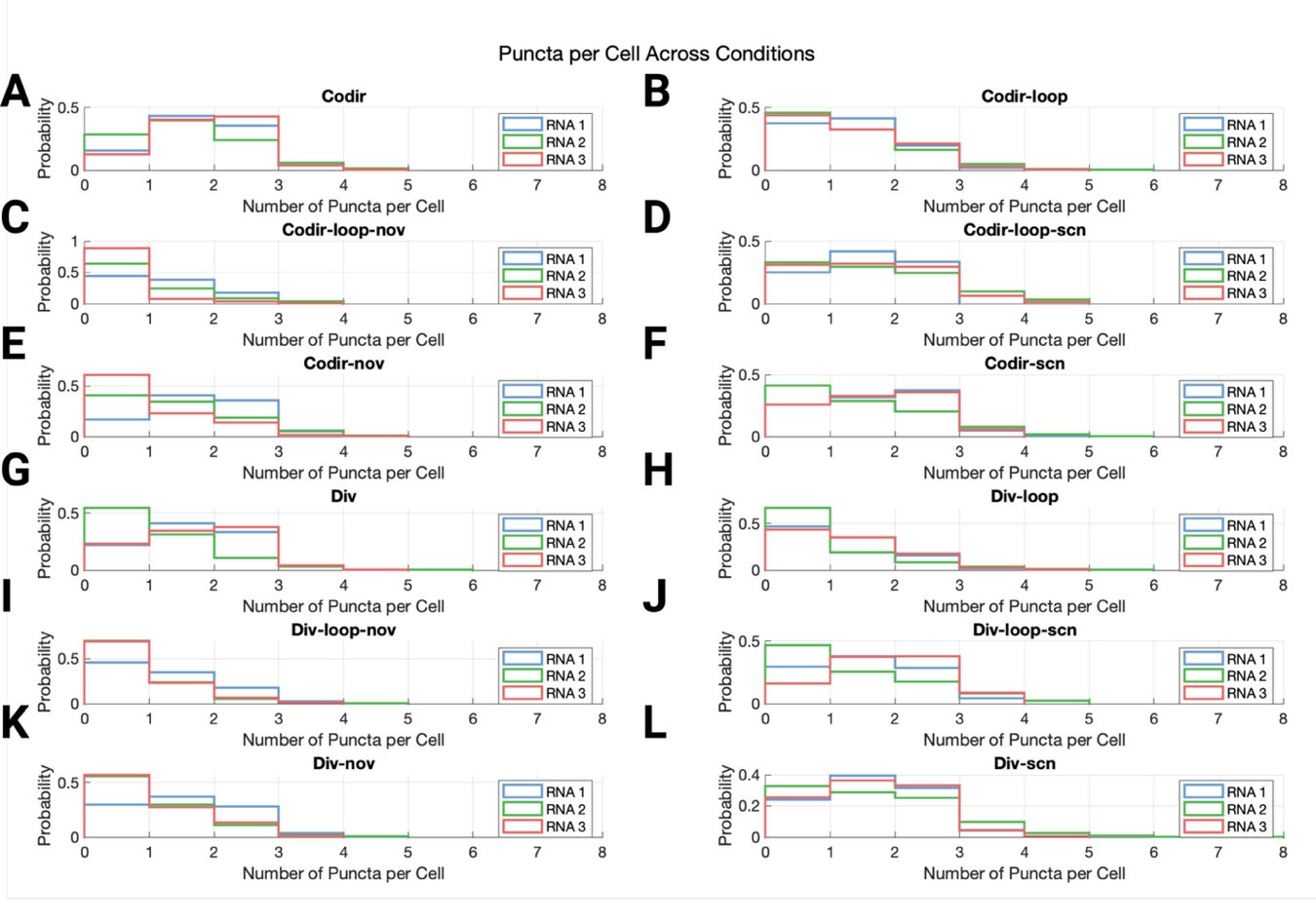

**Figure S3. Distributions of RNA puncta number per cell across genes and conditions**.

Probability distributions of RNA puncta number per cell for RNA1 (blue), RNA2 (green), and RNA3 (red) across all twelve experimental conditions are shown from **A** to **L.** Specifically: (**A**) Codirectional, unlooped, no drug; (**B**) Codirectional, looped, no drug; (**C**) Codirectional, looped, novobiocin-treated; (**D**) Codirectional, looped, seconeolitsine-treated; (**E**) Codirectional, unlooped, novobiocin-treated; (**F**) Codirectional, unlooped, seconeolitsine-treated; (**G**) Divergent, unlooped, no drug; (**H**) Divergent, looped, no drug; (**I**) Divergent, looped, novobiocin-treated; (**J**) Divergent, looped, seconeolitsine-treated; (**K**) Divergent, unlooped, novobiocin-treated; (**L**) Divergent, unlooped, seconeolitsine-treated.

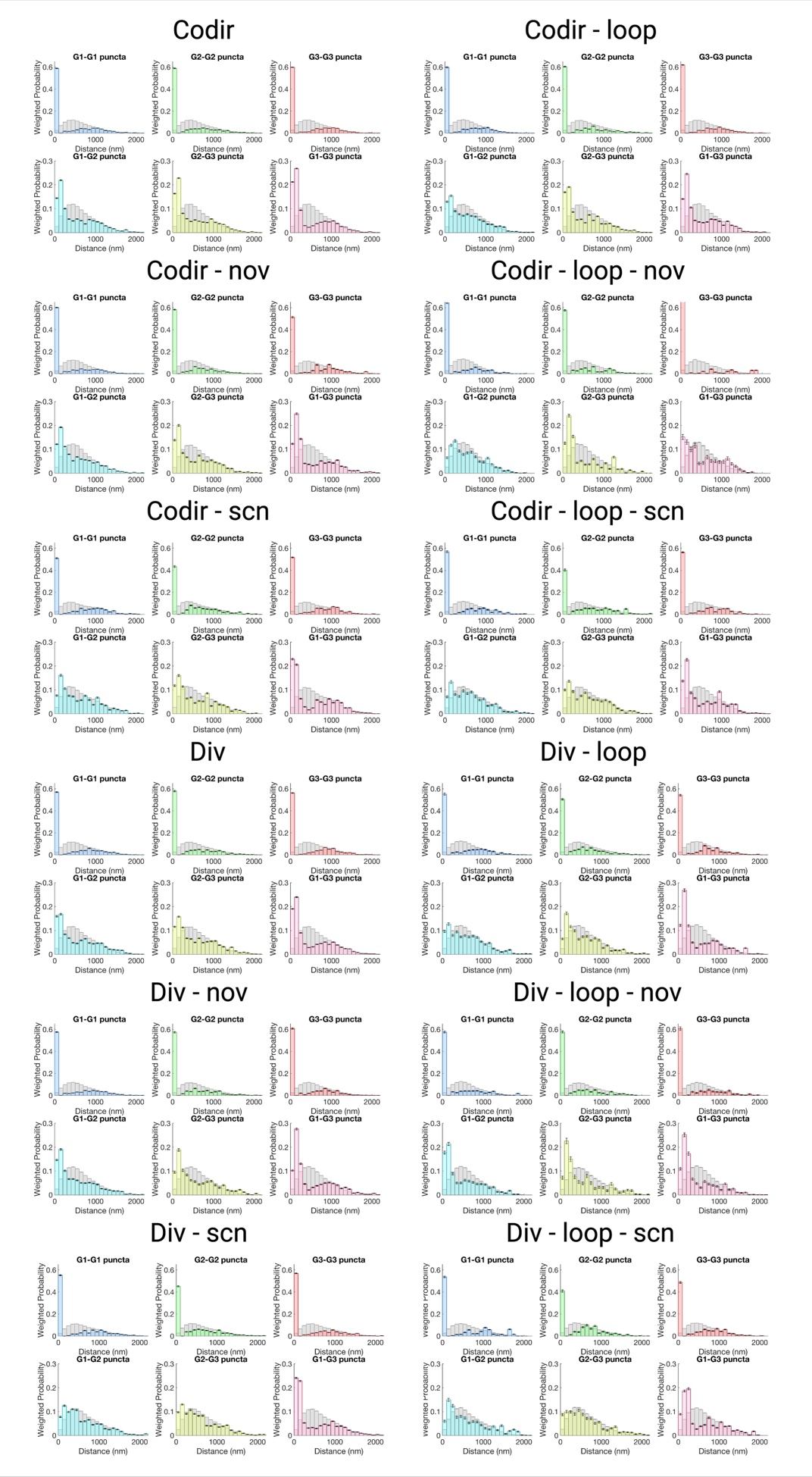

**Figure S4. Pairwise distance distributions of RNA molecules from the same or different genes under all experimental conditions**.

Each panel shows both intragenic (G1–G1, G2–G2, G3–G3; top row) and intergenic (G1–G2, G2–G3, G1–G3; bottom row) pairwise distance distributions, where distances are weighted by the number of RNA molecules in each punctum. Grey bars are pairwise distance distributions calculated in the same way but using computationally scrambled RNA molecule coordinates from each punctum in the same cells.

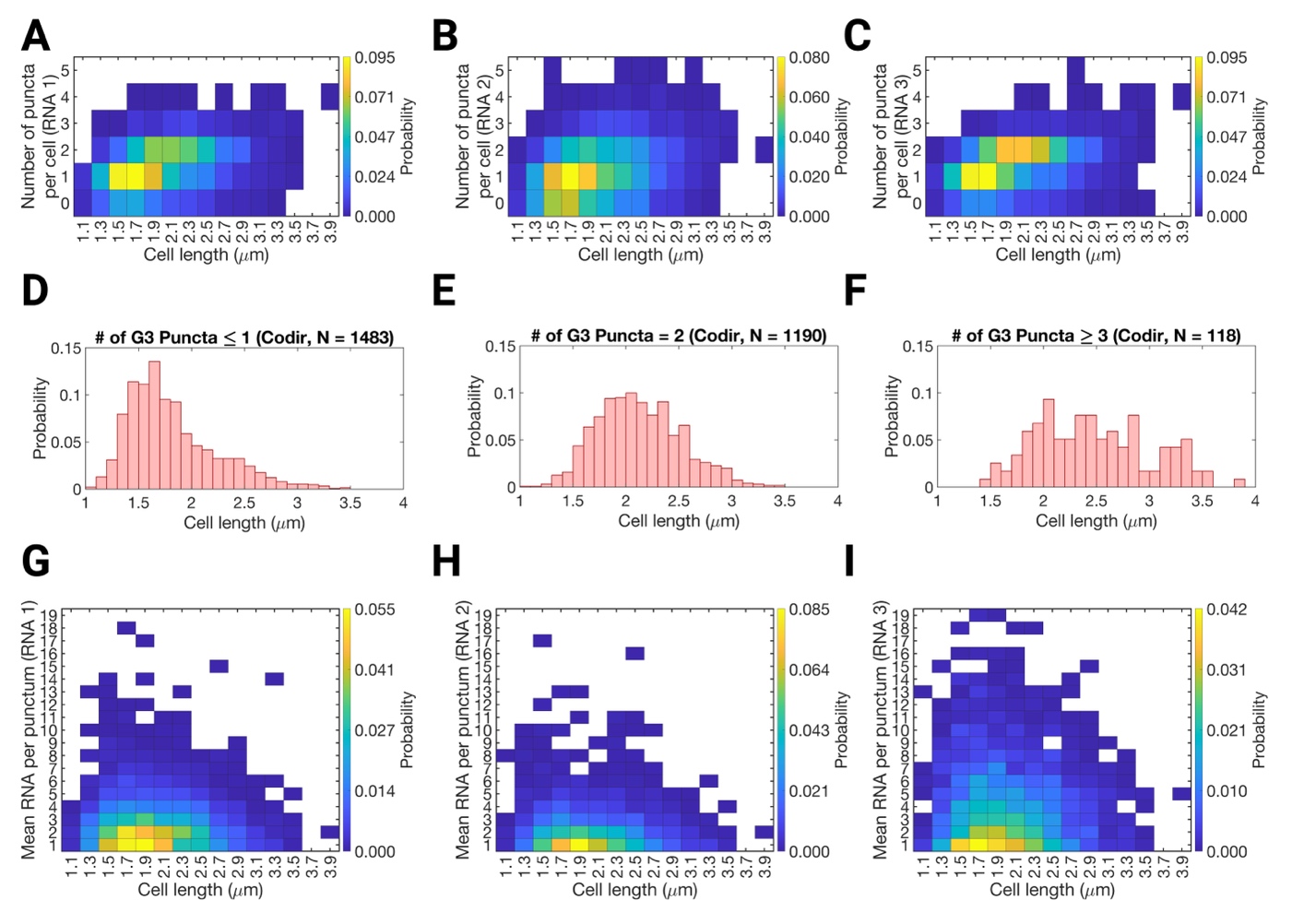

**Figure S5. Relationship between cell length, puncta number, and RNA content in the codirectional construct.**
**(A–C)** Two-dimensional histograms showing the number of puncta per cell as a function of cell length for RNA1 (**A**), RNA2 (**B**), and RNA3 (**C**) in the codirectional construct. Shorter cells typically contained one or no punctum, whereas longer cells more frequently contained multiple puncta, consistent with an increase in chromosome copy number as the cell cycle progresses. **(D–F)** Distributions of cell lengths for cells containing ≤ 1 (**D**), 2 (**E**), or ≥ 3 (**F**) puncta (using G3 as an example), showing that cells with multiple puncta are biased toward longer lengths, indicative of replicated nucleoids. **(G–I)** Two-dimensional histograms showing the mean number of RNA molecules per punctum as a function of cell length for RNA1 (**G**), RNA2 (**H**), and RNA3 (**I**) in the codirectional construct. In contrast to puncta count, RNA copy number per punctum shows little dependence on cell length.

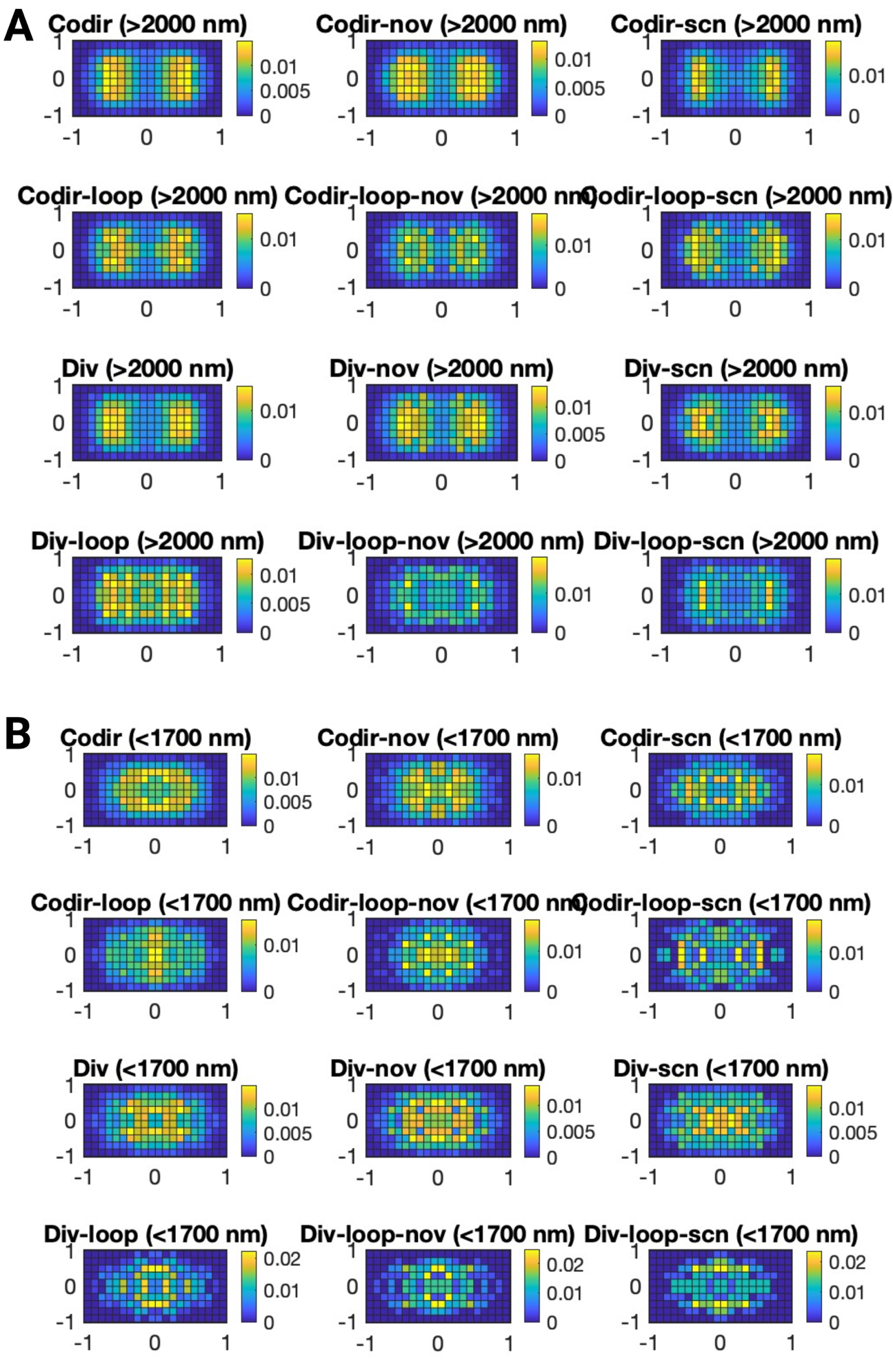

**Figure S6.** Two-dimensional heatmaps of combined RNA puncta localizations compiled from long (> 2.0 µm; **A**) and short cells (< 1.7 µm; **B**) at each condition.

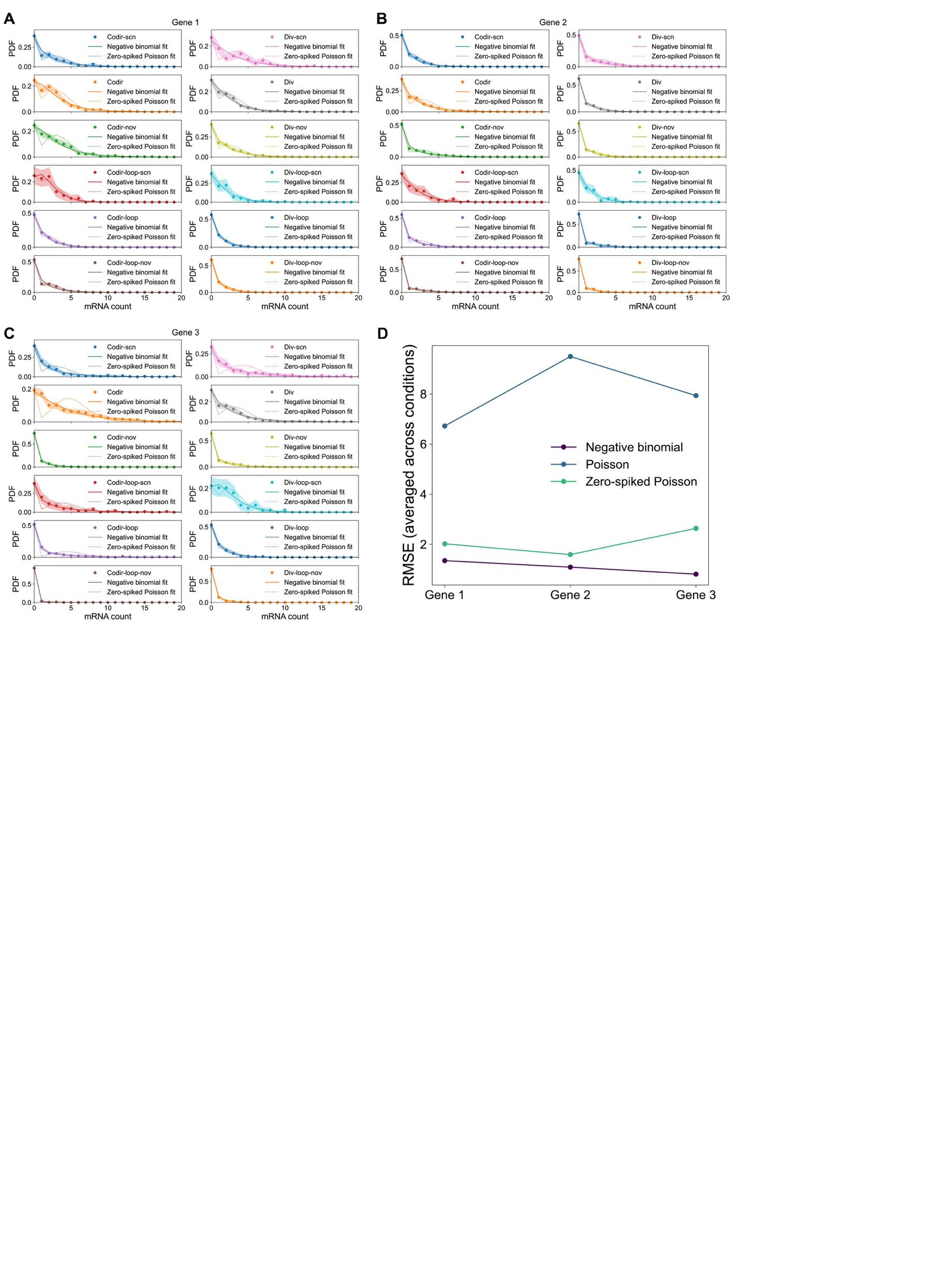

**Figure S7. Distributions of RNA copy number per cell for short cells (< 1.7 µm) across genes and conditions.**

Probability distributions of RNA copy numbers per cell for short cells (< 1.7 µm) for *G1* (**A**), *G2* (**B**) and *G3* (**C**) across the twelve experimental conditions. Each distribution is fit with either a Poisson (not shown for simplicity), a zero spike+ Poisson (dashed curve), and a negative binomial distribution (solid curves). The shaded areas are boot-strapped errors of the distribution. (**D**) Comparison of the summed root mean square errors (RMSE) of the three genes distributions fit with Poisson (blue, Zero-spike Poisson (green), and negative binomial (purple) distributions. For all the three genes, the negative binomial fitting has the least RMSE, indicating the best fit.

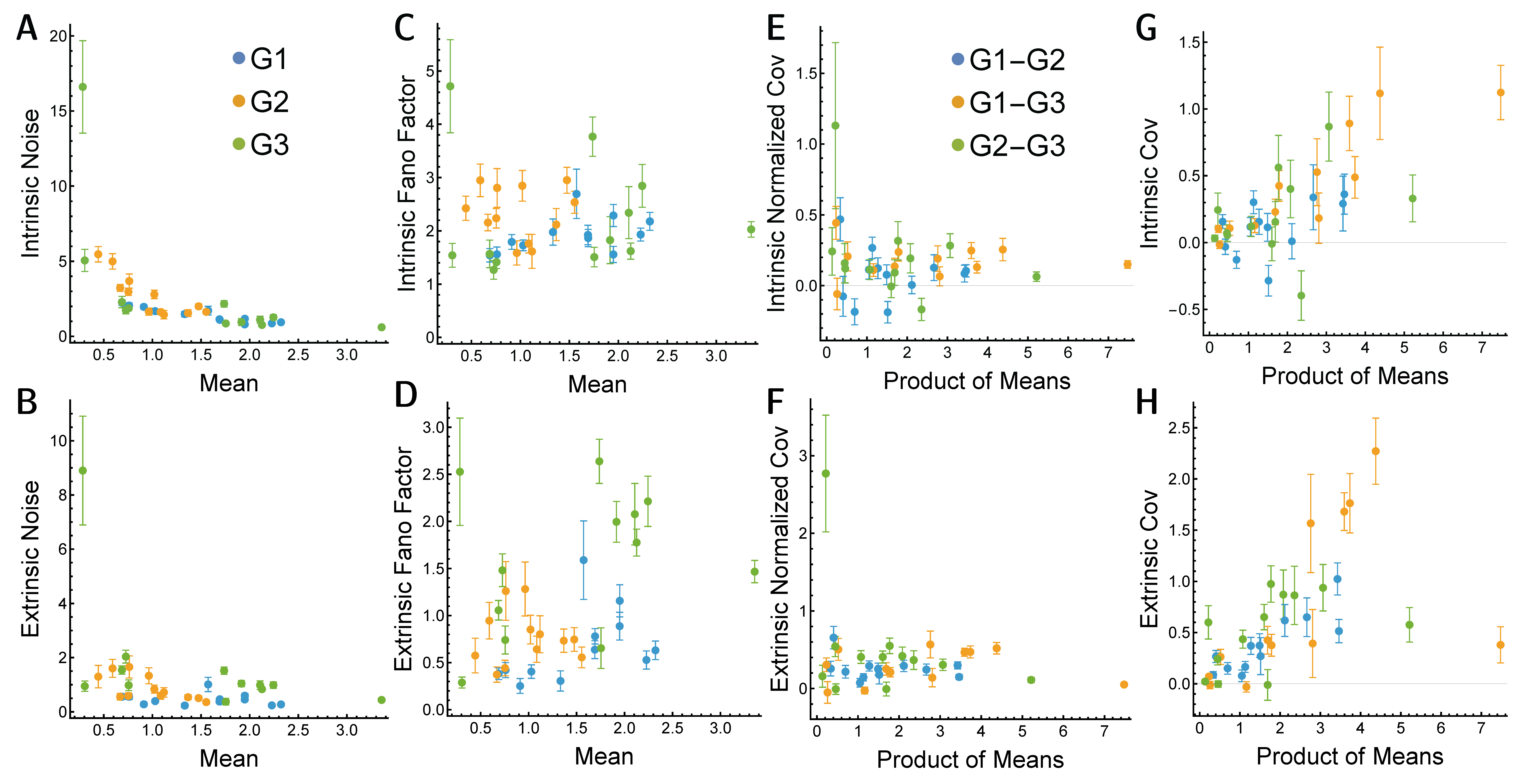

**Figure S8**. Dependence of decomposed intrinsic noise (**A**), extrinsic noise (**B**), intrinsic Fano factor (**C**) and extrinsic Fano Factor (**D**) on the mean RNA expression levels, and the dependence of decomposed intrinsic covariance (**E**), extrinsic covariance (**F**), intrinsic covariance (**G**), and extrinsic covariance (**H**) on the product of the corresponding means of two RNA expression levels.

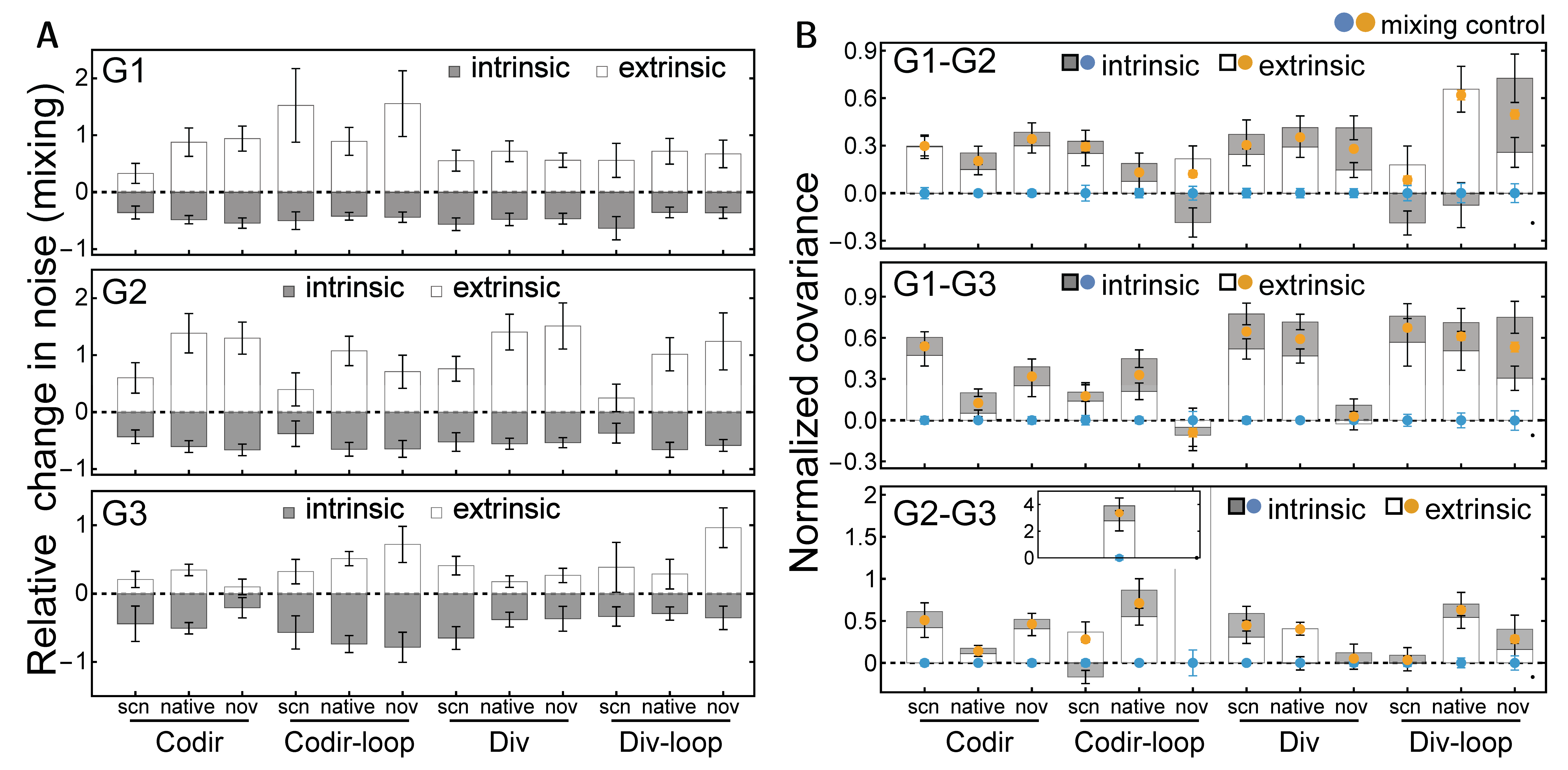

**Fig.S9. Mixing control of intrinsic-extrinsic decompositions.**

(**A**) Recomputed relative changes in intrinsic and extrinsic noise when RNA copy numbers between the two halve cells were computationally scrambled show that the intrinsic noise (gray bars) was significantly diminished while extrinsic noise (white bars) increased.

(**B**) Normalized intrinsic (blue dots) covariances between each gene pair were nearly completely diminished when RNA copy numbers between the two halve cells were computationally scrambled. The intrinsic covariance (gray bars) and extrinsic covariances (white bars) computed from the original data were shown as a comparison. Note the order of bars is switched compared to **Fig. 4** to aid the comparison between scrambled extrinsic covariances (yellow dots) and the original data.

The changes in apparent intrinsic/extrinsic statistics under this control demonstrate that measured data are not well mixed within the cell.

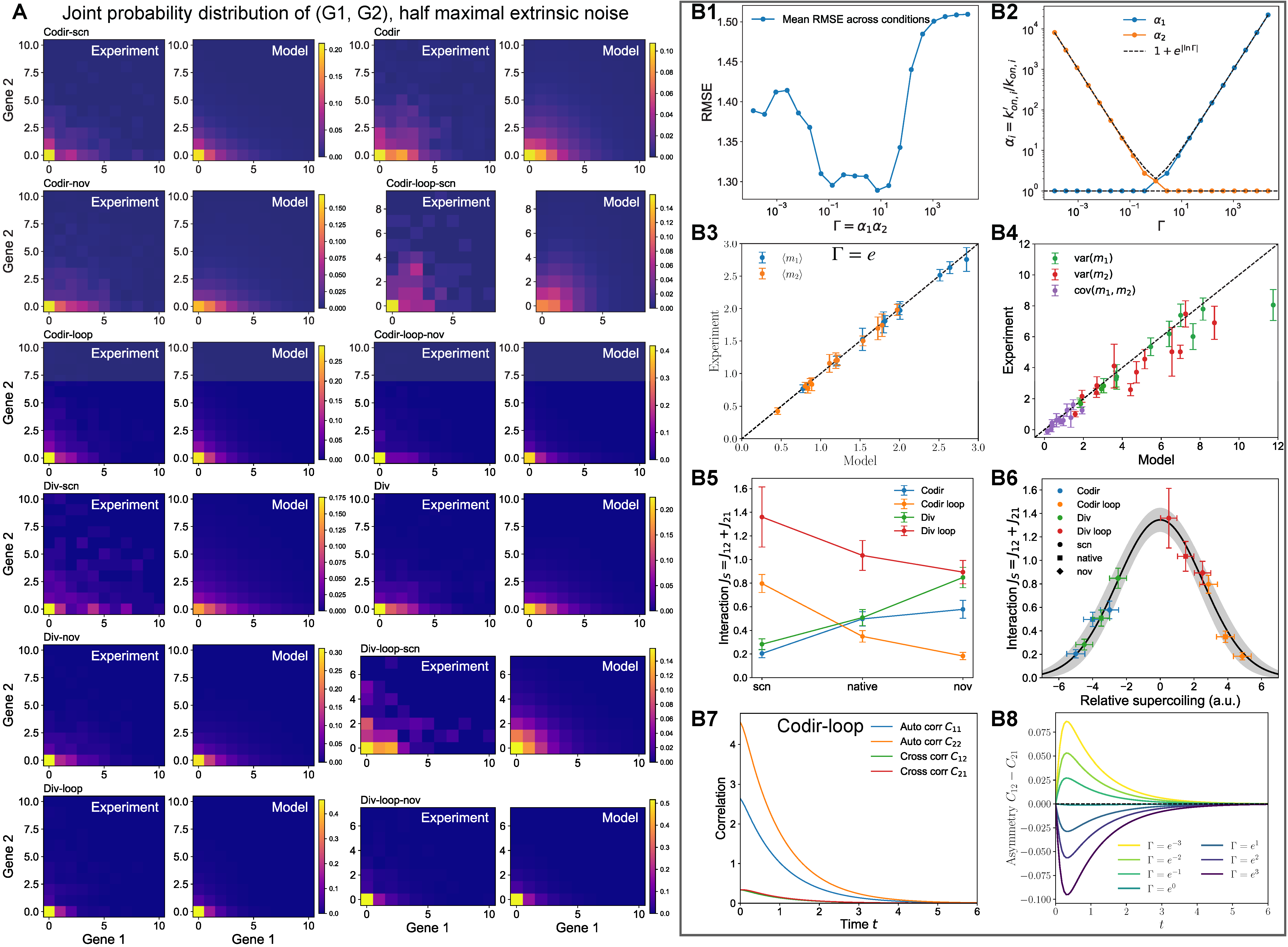

**Figure S10. Fitting the 4-state stochastic gene expression model of joint distribution of *G1* and *G2* when considering extrinsic noise at half maximum strength.**

(**A**) Joint mRNA count distribution $P(m_{1},m_{2})$ for experiments (left) and model fit (right) across all 12 experimental conditions, for considering extrinsic noise at half maximum strength (same as Fig.5 in the main text)

(**B**) Dependence of the fit on the amplification ratio $\Gamma=\alpha_{1}/\alpha_{2}$.

(**B1**) The fit accuracy, quantified by the root mean square error (RMSE) in the joint distribution averaged over all conditions, versus amplification ratio $\Gamma=\alpha_{1}/\alpha_{2}$, $\alpha_{i}=k_{{on,i}^{'}}/k_{on,i}$. Fit quality is roughly invariant for $-2⪅ln\Gamma⪅2$. (b2) The fit amplification factors $\alpha_{i}$ versus $\Gamma$. Outside the best fit range $|\Gamma|>2$, the interaction is unidirectional, with one $\alpha\approx1$ and the other $\approx|\Gamma|$ (dashed line). (**B3-B4**) The model captures both the mean (b3) and the variance and covariance (b4) of transcript numbers if $\Gamma$ is held constant in fits, in this case to $\Gamma=e^{2}$. (**B5-B6**) For fits with constrained $\Gamma$ (here $\Gamma=e^{2}$), the construct-dependence of the changes in total interaction parameter $J_{S}=J_{12}+J_{21}$ with drug treatment (**B5**) and the collapse of the fit interaction parameter onto a non-monotonic bell curve (**B6**) are qualitatively similar to within the best fitting regime $-2⪅ln\Gamma⪅2$ (c.f. **Fig. 5E, F**). (**B7**) The autocorrelation $C_{ii}(t)=\langle m_{i}(t^{'})m_{i}(t^{'}+t)\rangle-\langle m_{i}\rangle^{2}$ and cross correlations $C_{ij}(t)=\langle m_{i}(t^{'})m_{j}(t^{'}+t)\rangle-\langle m_{i}\rangle\langle m_{j}\rangle$. The decay in correlations can be used to further constrain timescales in expression kinetics. (**B8**) The asymmetry in cross correlation $C_{12}-C_{21}$ depends sensitively on $\Gamma$ and thus can be used to determine the dominant direction of the gene-gene interaction.

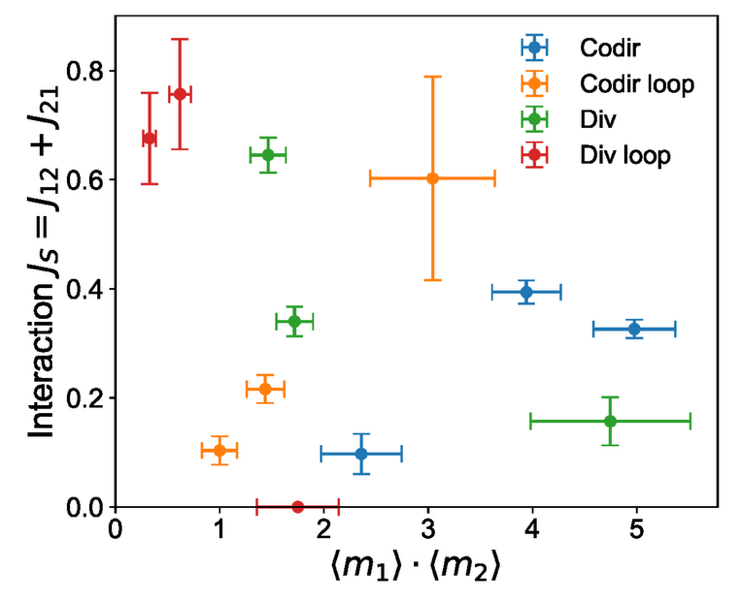

**Figure S11. The interaction strength** $\boldsymbol{J}_{\boldsymbol{S}}$**calculated at half maximum extrinsic noise strength had no clear dependence on the product of the mean expression levels of two genes.**

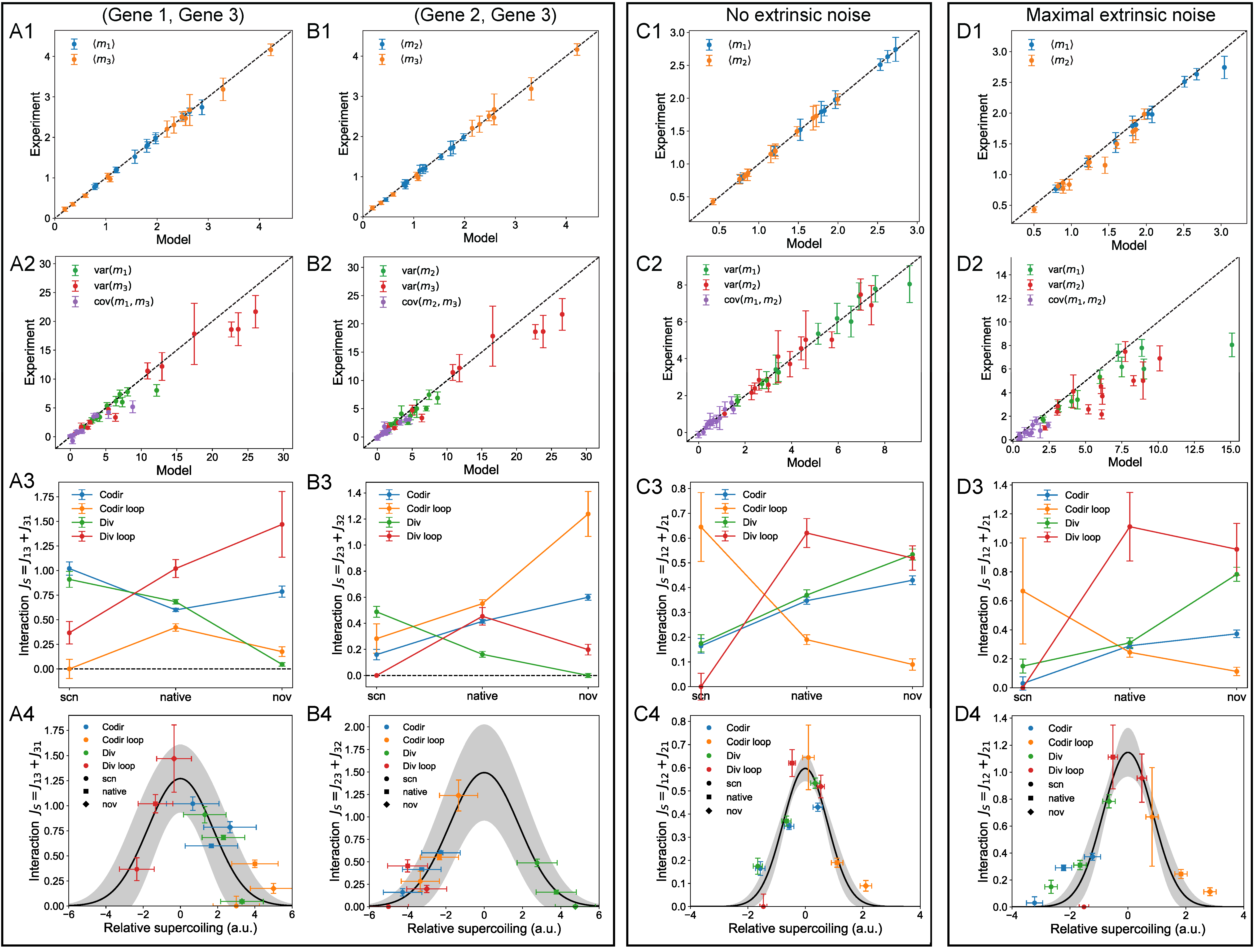

**Figure S12.** (**A**) Joint statistics and fitting results for G1 and G3 expression. The model captures both the mean (**A1**) and the variance and covariance (**A2**) of transcript numbers for the joint distribution of G1 and G3 expression. The changes in total interaction parameter $J_{S}$ with drug treatment are construct-dependent (**A3**) and the fit interaction parameter can be collapsed onto a non-monotonic bell curve (**A4**).

(**B**) Same for G2 and G3 expression.

(**C-D**) Joint statistics and fitting results for G1 and G2 expression, for considering zero extrinsic noise (**C**) and maximum amount of extrinsic noise (**D**). The model still captures the measured first (**C1, D1**) and second (**C2, D2**) moments of gene expression statistics, as well as a non-monotonic relation between the interaction strength $J_{S}$ (**C3, D3**) and the relative supercoiling (**C4, D4**).
